# An opioid efficacy switch for reversible optical control of peripheral analgesia

**DOI:** 10.1101/2024.12.16.628735

**Authors:** Luca Posa, Giovanna Romano, Xiang Ji, Saif Khan, Bruno Matos Paz, Gye Won Han, Antonina L. Nazarova, Saheem A. Zaidi, Mohsen Ranjbar, Kristen Pleil, Vsevolod Katritch, Cornelius Gati, Dirk Trauner, Joshua Levitz

## Abstract

The mu-opioid receptor (MOR) is a major target for the treatment of pain. However, opioids are prone to side effects which limit their effectiveness as analgesics and can lead to opioid use disorders or, even, lethal overdose. The systemic administration of opioid agonists makes it both very difficult to decipher their underlying circuit mechanisms of action and to limit drug action to specific receptor subpopulations to isolate therapeutic effects from adverse side effects. Here we design, synthesize, and characterize a reversibly photoswitchable morphinan agonist termed “azo-morphine-3” (**AM-3**) which interconverts from low to high efficacy in response to different wavelengths of light to enable optical control of MOR signaling. Cryo-EM structures of the low efficacy “*trans*” and high efficacy “*cis*” states of **AM-3** bound to the MOR reveal distinct binding modes of the photoswitchable azobenzene moiety, each inducing unique structural dynamics, providing insight into the molecular basis of agonist efficacy. In mice, **AM-3** drives reversible and repeatable optical control of anti-nociception with a reduced side effect profile owing to its restriction to the periphery and its ability to be locally activated at the site of pain.

## Main

The MOR, a Gi/o-coupled family A G protein-coupled receptor (GPCR), is the major target of opioid analgesics, including morphine (**Fig. 1a**), oxycodone, and fentanyl^1^. The MOR is expressed broadly and has been found to contribute to various forms of analgesia in both the central and peripheral nervous system^2^. The effectiveness of opioids for the treatment of pain has led to the desire for new opioid-based strategies that can maintain analgesic properties while minimizing adverse side effects. This has included extensive efforts to develop G protein-biased and/or low intrinsic efficacy agonists^3–5^. Alternatively, given that many opioid-driven side effects, including respiratory depression and addiction, are centrally mediated in the brain, peripherally-restricted opioids have emerged as an attractive alternative approach^6,7^. However, gastrointestinal side effects and peripheral tolerance occur outside of the central nervous system^8^, highlighting the limitations of this approach and emphasizing the need for more advanced targeting mechanisms.

**Figure 1.**
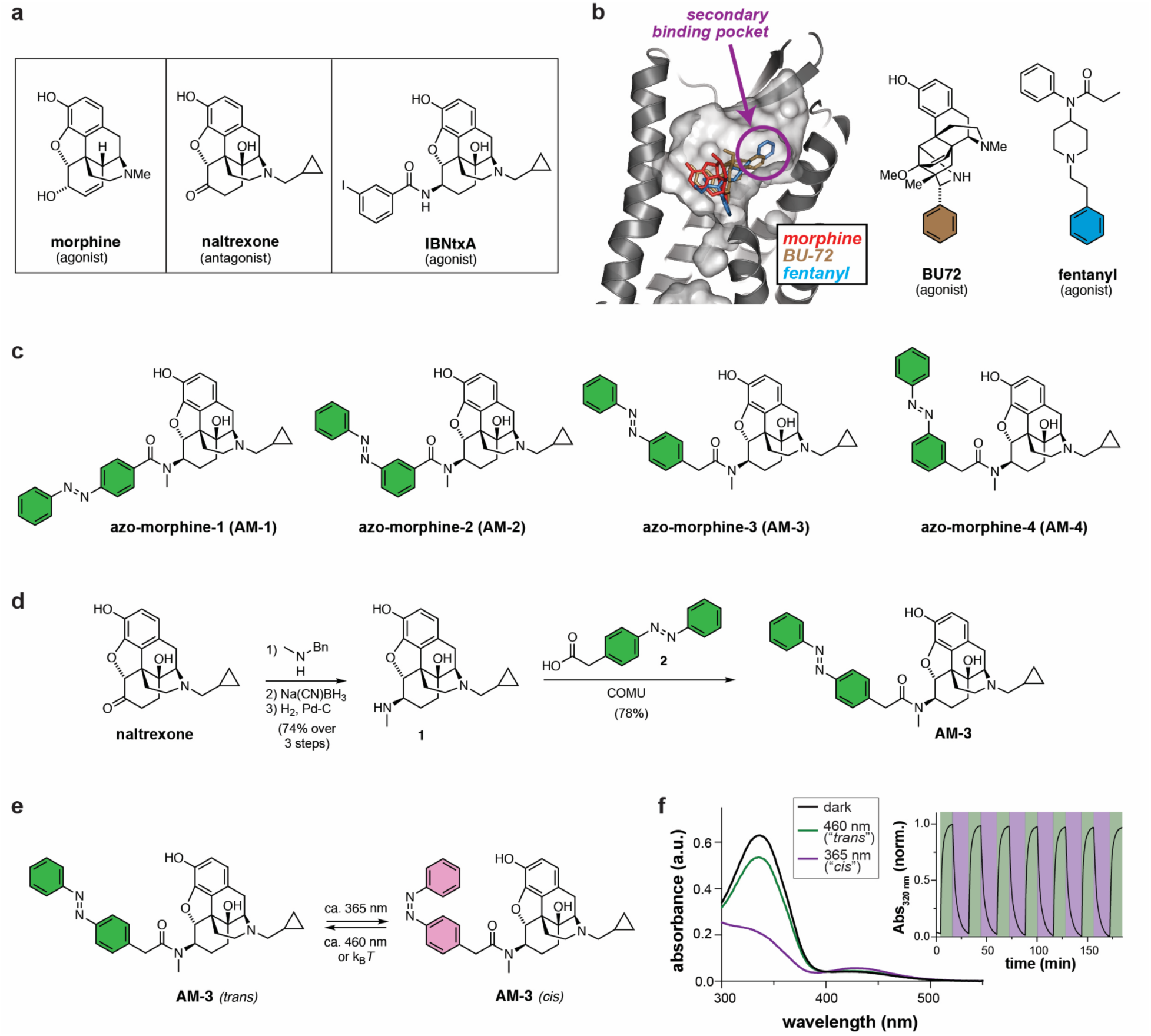
Structure-guided design and synthesis of azo-morphines. **a,** Chemical structures of morphinan ligands. **b,** Left, alignment of structures showing binding poses of morphine (PDB: 8EF6), BU-72 (PDB: 5C1M), and fentanyl (PDB: 8EF5). Right, chemical structures of BU-72 and fentanyl with rings that extend into the secondary subpocket colored. **c,** Chemical structures of azo-morphine variants tested in this study. **d,** Synthetic scheme for AM-3. **e-f,** Isomerization of AM-3 between *trans* and *cis* showing chemical structures (e) and UV/vis spectroscopy following different illumination conditions (f). Inset in (f) shows reversible photoswitching of AM-3 following alternating 460 nm (green) and 365 nm (magenta) illumination.

Photopharmacology, the use of light-sensitive ligands, represents a powerful approach to enable spatiotemporally precise control of drug action^9,10^, and may be a viable strategy for local opioid-driven pain relief. Recently reported photopharmacological designs for *in vivo* optical control of opioid agonists have employed caged approaches that irreversibly release ligands in response to illumination^11–13^. In contrast, photochromic ligands, which employ reversible photoswitchable moieties, such as azobenzenes, may provide a higher degree of spatiotemporal control to enable reversible and repeatable control of ligand action. To date, reversibly photoswitchable azobenzene-based opioid agonists using fentanyl scaffolds have exhibited tonic MOR activation in the relaxed *trans* state, with decreased activation in the excited *cis* form^14–16^. Morphinan-based photochromic agonists with enhanced activity in the thermodynamically less stable *cis* state would be ideal tools for both the mechanistic interrogation and precise therapeutic targeting of opioid action.

A major challenge in the design of reversible, photoswitchable ligands for GPCRs, including MORs, is a limited understanding of the molecular basis of ligand efficacy. In principle, the development of efficacy photoswitches^17^, ligands which change their maximal efficiency of target activation with light, would provide powerful tools for dissecting the structural and biophysical basis of GPCR agonism. Here, we use a structure-guided approach to develop a family of azobenzene-conjugated “azo-morphines”. Subtle tweaks in linker length and attachment geometry produce dramatic differences, with one variant “**AM-3**” showing a large difference in efficacy between *trans* and *cis* following illumination with blue versus UV light. Patch clamp electrophysiology experiments reveal the ability of **AM-3** to provide reversible, repeatable, and bistable control of MOR-driven G protein signaling in live cells. We probe the basis of this light-dependent effect on ligand efficacy by determining cryo-EM structures of *cis*-**AM-3** and *trans*-**AM-3** bound to MOR, revealing that a distinct azobenzene moiety pose is associated with high versus low efficacy agonism. Application of **AM-3** *in vivo* in mice enables local, light-dependent analgesic action in both acute and chronic pain with reduced central and peripheral side effects. Together these features provide a new approach to opioid pharmacology and open the door for further design and application of targeted, reversible GPCR photopharmacology.

### Design and Synthesis of Azo-Morphines

To produce potential photoswitchable morphinan agonists we examined agonist-bound MOR X-ray crystallography^18,19^ and cryo-EM structures^4,20–23^ and noted the presence of a secondary binding subpocket that accommodates aromatic moieties of a variety of agonists including fentanyl and BU-72 (**Fig. 1b**). We reasoned based on the orientation of morphinans in G protein-bound structures^18,19,21^, that chemical extension from the morphinan core may enable differential engagement of this subpocket via bent *cis* and extended *trans* azobenzene configurations. Notably, the agonist IBNtx employs a naltrexone-like core with a similar extension (**Fig. 1a**), which enables mode-switching of the common scaffold from an antagonist to an agonist^24–26^. We thus designed a small library of “azo-morphine” compounds using a naltrexone scaffold and variable length linkers to an azobenzene conjugated in the meta or para position (**Fig. 1c**). Molecular docking analysis supported our hypothesis that such compounds can bind with the morphinan group in the primary binding pocket and azobenzenes at positions toward the extracellular face of the receptor. Intriguingly, while the azobenzenes consistently occupied the secondary subpocket in *cis*, in *trans* various positions were predicted with similar binding scores (**Extended Data Fig. 1**).

Synthesis of the azo-morphines started with reductive amination of naltrexone with methyl benzyl amine, followed by debenzylation, to yield the known secondary amine **1** (**Fig. 1d**)^27^. Amide coupling with a variety of azobenzene carboxylic acids, such as **2**, then yielded the azo-morphines, specifically **AM-3** (see Supplementary Information for a detailed description of all syntheses). Photophysical characterization showed that **AM-3** behaves as a regular azobenzene, with maximum *cis* content achieved at 360 nm (*trans*:*cis* = 28:72) and maximum *trans* content at 460 nm (*trans*:*cis* = 84:16) (**Fig. 1e, f; Extended Data Fig. 2**). Thermal relaxation to the *trans* form was found to be slow (t_1/2_ = > 24 h in PBS:DMSO = 9:1) (**Extended Data Fig. 2b**). Similar photophysical properties were observed across all azo-morphine variants tested here (**Extended Data Fig. 2a, b**).

### Functional characterization of Azo-Morphines

We tested **AM-1, AM-2, AM-3,** and **AM-4** using patch clamp electrophysiology in HEK 293T cells expressing the rat MOR and the G protein-coupled inward rectifier potassium channel (GIRK) as a reporter of G protein activation. Each ligand was applied at 100 nM in the presence of 460 nm light to maintain the maximal *trans* state and then light was switched between 365 nm and 460 nm to toggle the ligand between primarily *cis* and primarily *trans*, respectively. Following at least two rounds of switching, the azo-morphine was removed and the full agonist DAMGO was applied to enable normalization of the current amplitudes induced by *trans* versus *cis* azo-morphine (**Fig. 2a; Extended Data Fig. 3**). The four ligands showed strikingly different behavior. AM-1 produced clear activation in *trans*, which was maintained but not altered by photoswitching to the *cis* state. In contrast, AM-2, AM-3, and AM-4 all showed some activation in *trans* followed by enhanced activation in *cis*. However, the extent of *trans* activation was substantially less with AM-3, pointing to a more pronounced difference in the ability of *cis* and *trans* forms of the ligand to activate the receptor. **Fig. 2b** shows a summary of the *cis*/*trans* activation ratio across all four ligands, highlighting the enhanced photoswitching of AM-3 relative to AM-2 and AM-4. Based on these results, we focused on AM-3 as the most promising azo-morphine variant.

**Figure 2.**
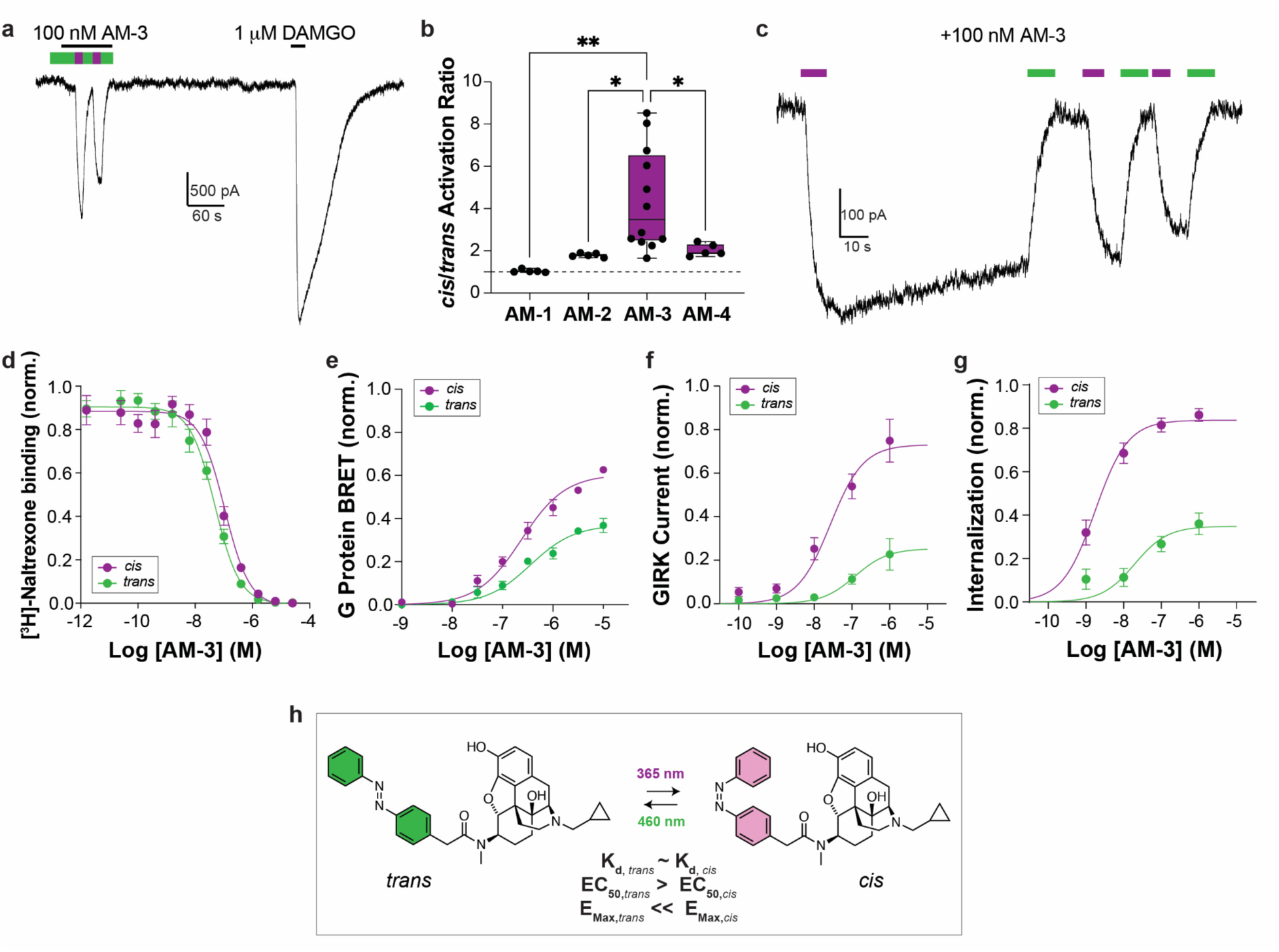
AM-3 is a photochromic MOR agonist which enables reversible efficacy switching. **a,** Representative patch clamp electrophysiology trace showing GIRK currents induced by *trans* or *cis*-AM-3 relative to DAMGO in cells expressing the MOR. Green=460 nm illumination; Magenta=365 nm illumination. **b,** Summary graph (box plot shows median and error bars show min and max) of relative activation by *cis* (365 nm illumination) versus *trans* (460 nm illumination) across azo-morphine variants at 100 nM. * p<0.05; ** p<0.001; 1-way ANOVA with multiple comparisons. **c,** Representative trace showing bistable, reversible, and repeatable photoactivation of the MOR by AM-3. **d-g,** Dose-response curves showing the difference between *cis* (365 nm illumination) and *trans* (460 nm illumination or dark-adapted) forms of AM-3 across assays that measure orthosteric binding (d), G protein activation (e), GIRK current activation (f), and receptor internalization (g). Values in e-g are normalized to the response to 1 μM DAMGO. **h,** Summary of AM-3 pharmacological data.

AM-3 produced robust photocurrents in response to 365 nm that were repeatable and were maintained in the dark prior to 460 nm illumination (**Fig. 2c**). This bistability is consistent with our UV/Vis spectroscopy results (**Extended Data Fig. 2**) and enables sustained MOR activation without the need for ongoing, potentially harmful UV illumination. We characterized the spectral properties of AM-3 and found that longer wavelengths than 365 nm were unable to efficiently drive photocurrents (**Extended Data Fig. 4a-b**). Importantly, comparable *cis* photo-activation via AM-3 was observed for rat, mouse, and human MOR subtypes (**Extended Data Fig. 5a, b**).

To better understand the mechanism of the enhanced activity in *cis* versus *trans*, we performed a series of dose-response titrations of AM-3 in either form. Radioligand binding experiments, using *cis*-AM-3 and *trans*-AM-3 against tritiated [^3^H]-Naltrexone, showed robust competition, with an estimated K_i_ of 35 nM and 16 nM, respectively (**Fig. 2d**). To detect MOR-mediated G protein activation, we used the ONE-GO G𝛂i3 bioluminescence resonance energy transfer (BRET) sensor, which detects agonist-evoked GPCR-mediated release of G𝛂i3-GTP^28^ (**Fig. 2e**). Both *trans*-AM-3 and *cis*-AM-3 produced clear responses of comparable potencies (EC_50,*cis*_=199 nM; EC_50,*trans*_=293 nM), but the maximal activation evoked by *cis*-AM-3 was substantially higher (E_max,*cis*_=0.59; E_max, *trans*_=0.36), suggesting that AM-3 serves as an efficacy photoswitch. To evaluate the dose-dependence of downstream signaling activity initiated by AM-3, we performed our photoswitching patch clamp GIRK current experiment across a range of concentrations to produce a dose-response curve for *cis* and *trans* (**Extended Data Fig. 6a, b**). *cis*-AM-3 showed a higher potency (EC_50,*cis*_=27 nM; EC_50,*trans*_=121 nM) and a ∼3 fold increase in Emax (E_max,*cis*_=0.73; E_max, *trans*_=0.25) (**Fig. 2f**), further supporting the efficacy switch model. Finally, we measured MOR internalization as a means of assessing the ability of AM-3 to initiate receptor desensitization and downregulation. Using a SNAP-tag based surface labeling assay, we found that extended treatment with both *cis*-AM-3 and *trans*-AM-3 can drive MOR internalization, but with increased potency (EC_50,*cis*_=1.7 nM; EC_50,*trans*_=12 nM) and efficacy for *cis* (E_max,*cis*_=0.85; E_max, *trans*_=0.32) (**Fig. 2g**). Live cell imaging confirmed that *cis*-AM-3 produced substantially more intracellular puncta from surface-labeled SNAP-tagged MOR compared to *trans*-AM-3 (**Extended Data Fig. 6c**). Notably, *cis*-AM-3 showed a lower Emax than DAMGO but was comparable to morphine across assays (**Extended Data Fig. 6d-g**). Altogether, these data show that both forms of AM-3 serve as partial MOR agonists, but that the *cis* form contains a modestly higher potency and a substantially enhanced efficacy relative to *trans* (**Fig. 2h**). Finally, we tested the two other major opioid receptor subtypes and found that AM-3 potently activates both the kappa and delta opioid receptors, but without any light dependence (**Extended Data Fig. 7**).

### Structural Basis of AM-3 Efficacy Switching

To probe the structural basis of the distinct efficacies of *cis* and *trans*-AM-3, we performed single particle cryo-EM experiments of the ternary complex consisting of either *cis*-AM-3 or *trans*-AM-3, wild type human MOR, wild type human Gαi1 and Gβ1 and Gγ2. We obtained high resolution reconstructions of the complex with the compound in either state, with an overall estimated resolution of 3.2 Å (*cis*-AM-3) and 3.1 Å (*trans*-AM-3) (**Fig. 3a, b**; **Extended Data Table 1**). For *cis*-the consensus refinement was followed by local refinement of the receptor and G protein heterotrimer separately and merged into a composite map, which was used for modeling and further structural analysis (**Extended Data Fig. 8**). For *trans*-AM-3, additional classification was performed using 3D Variability Analysis^33^, to select for subsets of particles with improved density for the AM-3 azobenzene group (**Extended Data Fig. 9)**. Our refined models closely resemble previously published active-state MOR structures (**Extended Data Fig. 10a, b**). Comparing *cis*-AM-3 and *trans*-AM-3 structures to the highest resolution MOR structure without the use of the receptor:G protein complex-stabilizing scFv16 (Mitragynine pseudoindoxyl MP, PDB ID: 7T2G), the receptors align well, with an overall RMSD of 0.79 Å and 0.62 Å for *cis*- and *trans*-AM-3, respectively. General structural hallmarks of GPCR activation, including the DR^3.50^Y, CW^6.48^xP^6.50^, P^5.50^-I^3.40^-F^6.44^ and NP^7.50^xxY^7.53^ motifs (values indicate Ballesteros–Weinstein numbering for GPCRs^29^), are identical to those derived from structures with prototypical agonists DAMGO, fentanyl and morphine (PDB ID: 8EFQ, 8EF5, 8EF6^21^). The interface between the nucleotide-free G protein and MOR are also very similar, representing a canonical MOR:Gi1 coupling state. One noteworthy difference between *cis*- and *trans*-AM-3 bound states is the relative angle between the N-terminus of G𝛂i, which is approximately 6 degree tilted towards the receptor, suggesting a closer engagement of the receptor with the G protein in the *cis* state, which could support its higher efficacy (**Fig. 3c**).

**Figure 3.**
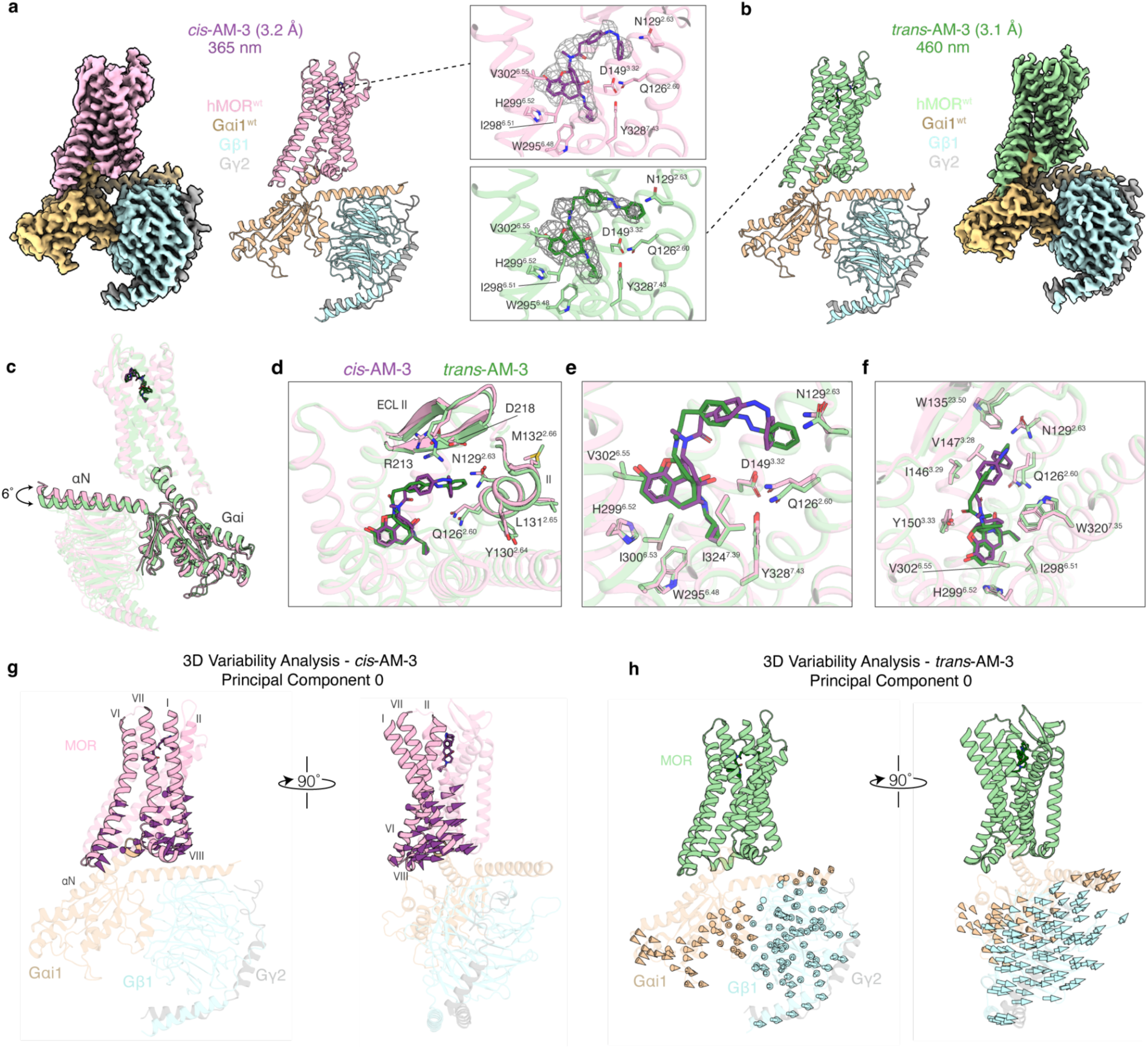
Cryo-EM reconstructions of *cis*-AM-3 and *trans*-AM-3 in complex with MOR:G protein heterotrimer. **a-b,** Cryo-EM reconstructions of the MOR:Gi complex (MOR: pink/light green, Gαi: beige, Gβ1: light blue, γ2: grey) bound to (a) *cis*-AM-3 (purple) and (b) *trans*-AM-3 (dark green), highlighting individual ligand densities (grey mesh). **c,** Superimposed structures showing that the Gαi αN helix is tilted 6° towards the *cis*-AM-3 bound MOR (pink), relative to *trans*-AM-3 (light green), suggesting stronger receptor-Gαi engagement. **d,** Overview of structural rearrangements in TM2 and ECL2, as a result of AM-3 binding in the *cis* or *trans* state. **e-f,** Detailed comparisons of the MOR orthosteric pocket from the (e) side and (f) top views. The terminal benzene of *trans*-AM-3 penetrates deeper towards TM2, resulting in differences in rotamer conformations of Q^2.60^ and N^2.63^. **g-h,** 3D Variability Analysis of the (g) *cis*-AM-3, and (h) *trans*-AM-3 bound MOR:Gi complexes. Vectors (arrows, color coded by chain) represent trajectories of motion along principal component 0 that are larger than 1 Å, highlighting distinct conformational dynamics between the *cis*-AM-3 and *trans*-AM-3 bound MOR:Gi complexes.

Our initial attempts of structure refinement for both datasets produced maps with strong and unambiguous density for the naltrexone scaffold. While a complete ligand density for *cis*-azobenzene was readily obtained (**Extended Data Fig. 8**), the *trans*-azobenzene group initially appeared somewhat dynamic, and an unambiguous ligand density was only obtained after extensive classification attempts (**Extended Data Fig. 9**). For both *cis*- and *trans*-AM-3 the azobenzene moiety protrudes into the secondary subpocket, located between TM1 and TM2, analogous to fentanyl and other synthetic opioids^21,23^ (**Fig. 3a, d-f**). The naltrexone scaffold is located in the canonical, orthosteric binding pocket, with a similar pose to the previously published structure of the irreversible morphinan antagonist beta-FNA (PDB ID: 4DKL)^18^ (**Extended Data Fig. 10c**). Previous work has suggested that the morphinan scaffold of agonists sits deeper in the orthosteric binding pocket, compared to antagonists^21,30,31^. Using the tertiary amine in beta-FNA as a reference point, the naltrexone moiety of *trans*-AM-3 sits 0.6 Å deeper, *cis*-AM-3 by 0.9 Å, and morphine by 1.1 Å, which, while subtle, shows a correlation with the efficacy of the respective ligand (**Extended Data Fig. 10c**).

The terminal benzene ring of *cis*-AM-3 shows hydrophobic interactions with side-chains Q^2.60^, N^2.63^, W^23.50^, V^3.28^ and I^3.29^, analogous to fentanyl-like scaffolds shown previously^21,23^. The *trans*-AM-3 structure occupies the same overall binding pocket, but due to isomerization around the azo group, the terminal benzene moiety protrudes 2.5 Å further towards TM2 (**Fig. 3a-f**). This results in a modest rearrangement of the extracellular half of TM2 moving towards TM1 by approximately 1.5 Å. In addition to this global change, we observe differences in rotamer conformations of Q^2.60^, N^2.63^, Y^2.64^, all of which are proximal to the azo-benzene groups, and have been previously linked to ligand efficacy^23^ (**Fig. 3d-f; Extended Data Fig. 10d-j**). Interestingly, occupation of the fentanyl subpocket in MOR by itself is not clearly predictive of agonist efficacy. For instance, the benzyl moiety of alvimopan, a non-morphinan MOR-specific antagonist, also penetrates this subpocket, and extends similarly towards TM2^32^. However, the precise location of the phenyl group of fentanyl more closely matches *cis*-AM-3, while *trans*-AM-3 penetrates the subpocket similarly to alvimopan (**Extended Fig. 10d-j**). Additionally, in *trans*-AM-3, both the linker and the proximal benzene protrude towards extracellular loop 2 (ECL2), which is similarly occupied by carboxylic acid in alvimopan (**Extended Fig. 10 e, i**). Both beta-strands in ECL2 point away from the core of the receptor by approximately 1.8 Å in *cis*, compared to the *trans* structure, in which the beta hair-pin in ECL2 is stabilized by an additional interaction between R213 and D218 (**Fig. 3d**). ECL2 repositioning has previously been seen upon agonist binding suggesting that this interaction may contribute to ligand efficacy^33^. Finally, while we observed distinct binding poses of the ligands at MOR in their respective states, in light of both our docking results and the exclusion of a substantial particle subset by 3D Variability Analysis for the *trans*-AM-3 structure, we cannot rule out that either alternative conformations of azobenzene in the bound ligand, or binding kinetics, contribute to the reduced efficacy of *trans-*AM-3.

In addition to a standard cryo-EM data processing pipeline, we performed 3D Variability Analysis (3DVA) to visualize structural dynamics within our respective datasets^34^. Strikingly, while the average conformation obtained from structural refinement only showed modest differences between *cis-* and *trans*-AM-3, we observed substantial differences in terms of conformational dynamics **(Fig. 3g, h)**. Most notably, 3DVA from the *cis* dataset shows a twisting motion of the receptor, relative to the G protein, which is routinely observed in nucleotide free GPCR:G protein complex structures^34–36^. The twisting motion in the *cis* structure is accompanied by a correlated motion within the receptor, mainly driven by a rotation of the intracellular portion of TM7/helix 8, which coincides with the appearance of a density for the C-terminal loop of the Gai subunit (**Supplementary Movie 1**). Analogous findings from MD simulations^21,23^, as well as NMR studies^33^, were previously described, where the degree of this rotation correlated with the efficacy of an agonist at MOR. On the other hand, 3DVA for the *trans* dataset shows a relatively rigid receptor, while we observed a surprising degree of dynamics within the G protein and its displacement relative to the receptor. It is noteworthy that we observe similar movements across all three determined principal components, suggesting that these dynamics are dominant within the respective datasets. Qualitatively, these dynamic movements within the ternary complex suggest a less stable interaction between MOR and the G protein in the *trans* state, while *cis* results in a more tightly coupled ternary complex.

### *In vivo* local control of pain with AM-3

Motivated by the robust efficacy switching observed in cultured cells, we asked if AM-3 may be employed *in vivo* for light-dependent antinociception in mice. As the MOR is strongly expressed in heat-sensitive nociceptors^37,38^, we turned to the Hargreaves test as an assay of thermal pain sensitivity in the hind paw. We injected either relaxed *trans*-AM-3 or pre-illuminated *cis*-AM-3 directly into the plantar surface of the hind paw (i.pl.) (**Fig. 4a**) and measured the paw withdrawal latency (PWL) over 45 minutes. *cis*-AM-3 produced an increase in PWL that peaked at 20 minutes, was dose-dependent, and was observed in the injected paw but not the contralateral paw (**Fig. 4b; Extended Data Fig. 11a-c**). At the same i.pl. injection dose of 5 μg/5 μL, morphine and *cis*-AM-3 showed comparable effects (**Fig. 4c; Extended Data Fig. 11b**). However, across all doses tested, *trans*-AM-3 showed no effect relative to vehicle (**Fig. 4b,c; Extended Data Fig. 11d-f**). This suggests that the modest efficacy observed for *trans*-AM-3 in cultured cells (**Fig. 2e-g**) is insufficient to drive a behavioral effect in this assay. The antinociceptive effect of *cis*-AM-3 was blocked by pre-injection of the MOR antagonist naloxone (20 μg/5 μL), confirming opioid receptor-dependence (**Fig. 4d**).

**Figure 4.**
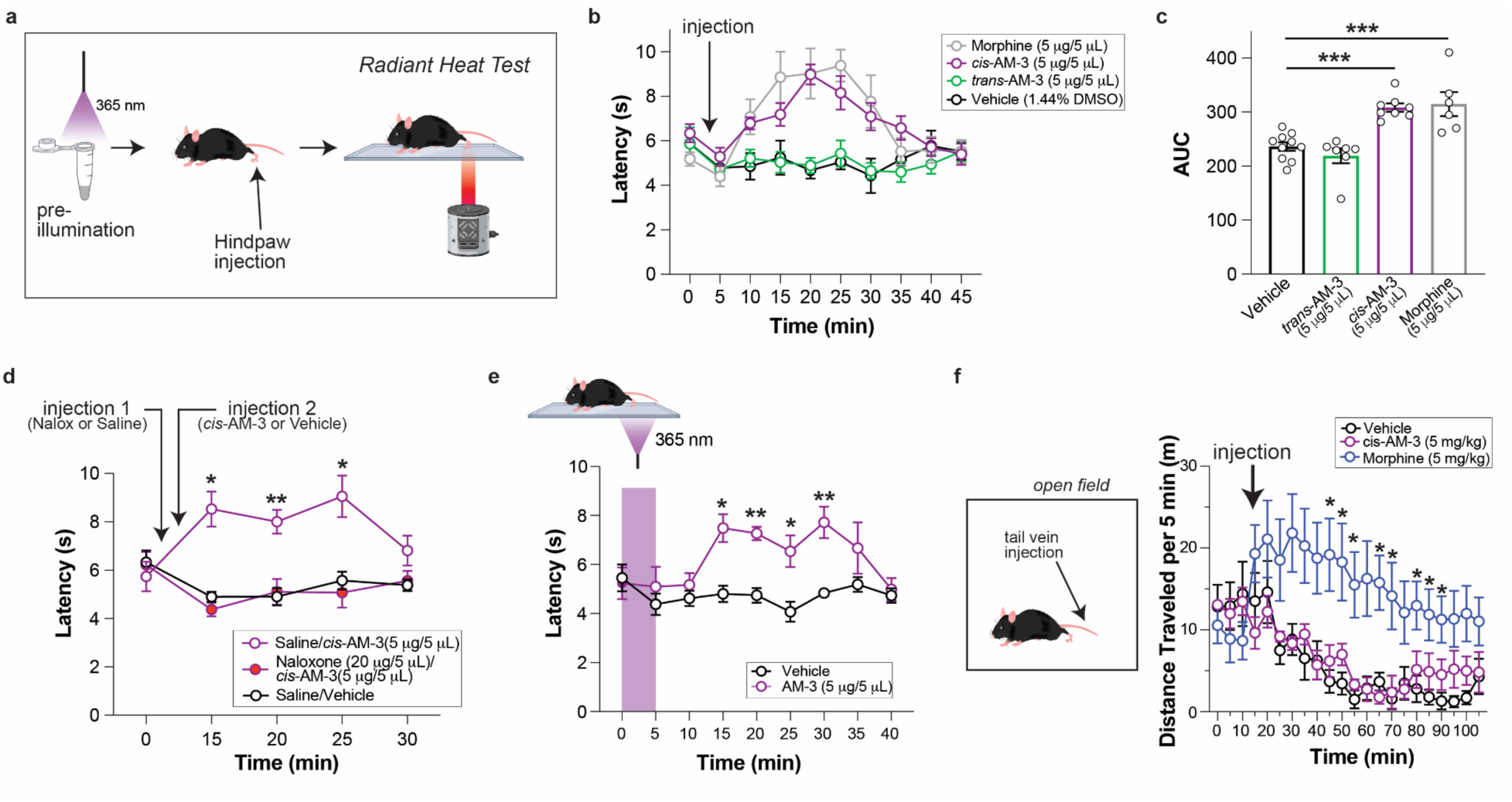
AM-3 enables optical control of peripheral antinociception. **a,** Schematic showing timing of pre-illuminated i.pl. drug administration prior to the Radiant Heat Test. **b,** Time course showing that *cis*-AM-3, but not trans-AM-3, increases the PWL and has a comparable time course to morphine. **c,** Area under the curve (AUC) analysis of cumulative analgesic effect of i.pl. *cis*-AM-3, *trans*-AM-3, morphine, and vehicle over 45 min post drug administrations. **d,** i.pl. pretreatment with naloxone 10 min prior to i.pl. administration of pre-illuminated *cis*-AM-3 completely blocked the antinociceptive effect. **e,** Time course showing that PWL increases after hind paw exposure to 365 nm light (purple bar). **f,** Locomotor activity showing that, unlike morphine, pre-illuminated cis-AM-3 does not promote hyperlocomotion in the open field test. All data are shown as mean±s.e.m. with n> 5 mice/condition; for (c) points represent individual mice; ∗p>0.05; ∗∗p<0.01; ∗∗∗p<0.001 vs vehicle (c, e, and f) or saline/vehicle (d). 1-way ANOVA was used for (c) and 2-way ANOVA was used for (d-f).

We next tested if AM-3 can be locally activated to its *cis* form *in vivo* to initiate antinociception. We i.pl. injected *trans*-AM-3 and used 365 nm light illumination through the bottom of the behavioral chamber to photoswitch to the *cis* form, which led to a large increase in PWL (**Fig. 4e**). In contrast, UV light had no effect in control mice that received a vehicle injection (**Fig. 4e**). We further showed that *in vivo* optical deactivation of AM-3 blocks its analgesic effect, as 465 nm light illumination following i.pl. injection of the pre-activated *cis* form of AM-3 prevented the observed increase in PWL in the absence of light illumination (**Extended Data Fig. 12**). Together these experiments demonstrate the ability of the antinociceptive activity of AM-3 to be controlled *in vivo* with light.

We next asked if AM-3 can accumulate in the central nervous system (brain and spine) as this is a key determinant of its ability to drive centrally-mediated side effects. We first tested this using the tail-flick assay, in which the latency to the tail reflex from the heat source is driven by MOR populations in the spine and brain^39,40^. Whereas systemic (intravenous, i.v.) tail-vein injection of morphine produced a robust increase in tail-flick latency, no tested doses of *cis*-AM-3 had an effect (**Extended Data Fig. 13**). In addition, *cis*-AM-3 did not produce hyperlocomotion in the open field test (**Fig. 4f**), a well-established acute effect of morphine mediated by mesolimbic dopaminergic systems in the brain related to reinforcement processes^2,41^. These data show that AM-3 does not have a central effect, which would enable it to serve as a peripherally restricted analgesic. To further test this, we conducted an *in vitro* permeability assay to evaluate AM-3 brain penetration assessing the role of drug transporters in its uptake into the brain. Both *cis* and *trans* forms of AM-3 exhibited an efflux ratio of ∼7 (**Extended Data Table 2**), indicating that it is a substrate for the P-glycoprotein efflux transporter, a multidrug-resistance membrane protein expressed at the blood-brain-barrier (BBB) that actively pumps various foreign substances out of cells^42^. These findings align with our behavioral data which show that AM-3 does not produce centrally-mediated effects.

Finally, we assessed the ability of AM-3 to alleviate chronic pain symptoms using the spared nerve injury (SNI) model of neuropathic pain (**Fig. 5a**). Three weeks post-surgery, mice developed thermal hyperalgesia in the ipsilateral paw (**Fig.5b-d**) and systemic AM-3 administration reversed this condition (**Fig. 5b-d; Extended Data Fig. 14**). Injection of pre-illuminated *cis*-AM-3 into the paw on the injured side relieved the thermal hyperalgesia and alternating illumination with 460 nm and 365 nm light enabled reversible control of this analgesic effect (**Fig. 5b; Extended Data Fig. 14c**), demonstrating the ability of AM-3 to undergo multiple rounds of photoswitching *in vivo*.

**Figure 5.**
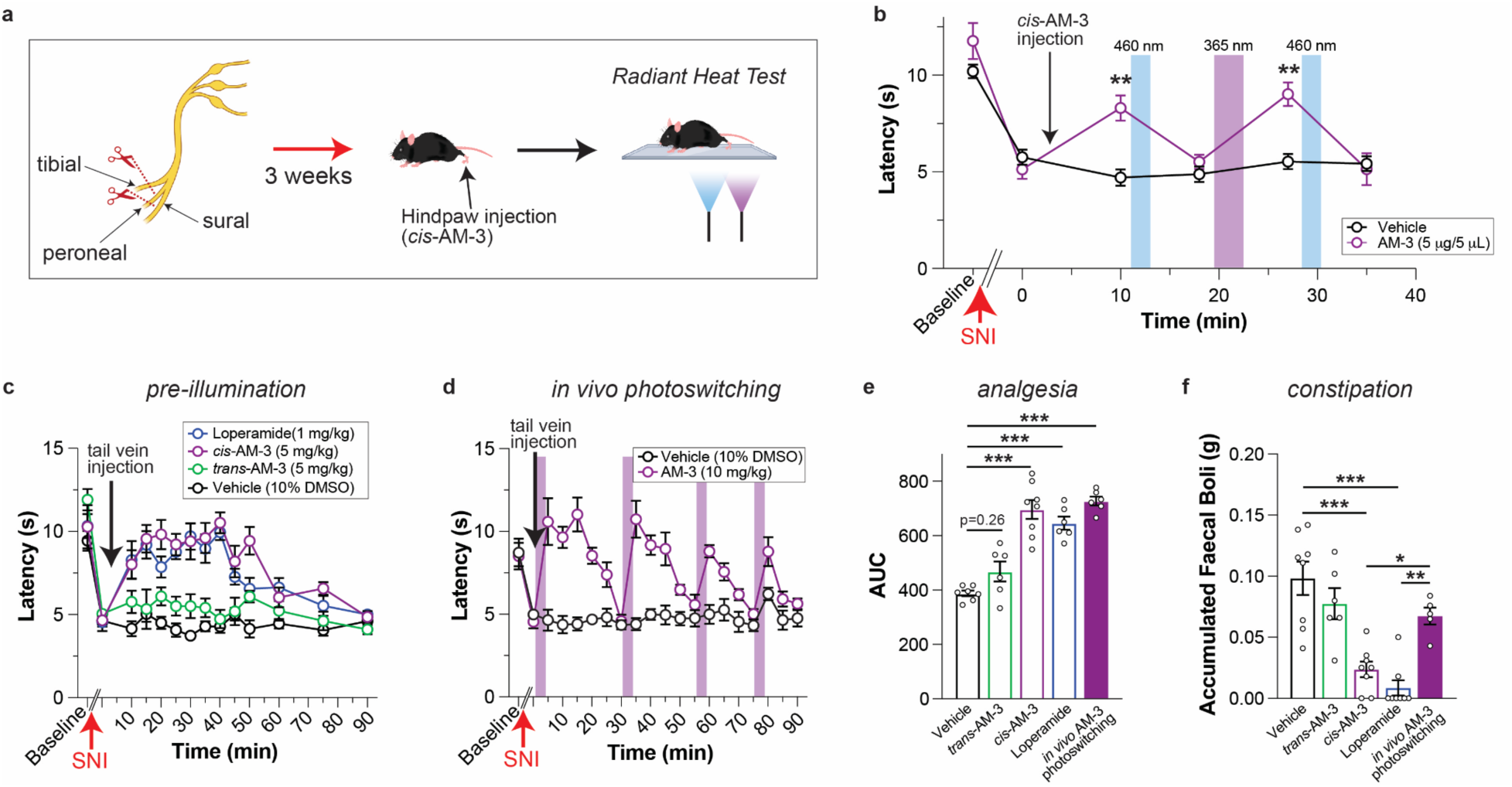
Alleviation of chronic neuropathic pain with reduced side effects via AM-3. **a,** Schematic showing spared nerve injury (SNI) model of chronic neuropathic pain followed by radiant heat test. **b,** Time course showing hyperalgesic effect of SNI followed by light-controlled analgesia via AM-3 following hindpaw injection. **c,** Time course showing analgesic effect of pre-illuminated *cis*-AM-3 and loperamide compared to *trans*-AM-3 and vehicle following tail vein injection. **d,** Time course showing in vivo photoactivation of AM-3 following tail vein injection in trans. **e,** Summary AUC analysis of analgesic effect of loperamide and AM-3 following pre-illumination (“*cis*-AM-3”) or in vivo photoswitching over the course of 90 min post-injection. **f,** Summary of accumulated faecal boli in SNI mice for all drug treatments during the 90 min time course used to assess hyperalgesia. All data in this figure are for the paw on the ipsilateral side relative to SNI. All data are shown as mean±s.e.m. with n> 5 mice/condition; for (e, f) points represent individual mice; ∗p>0.05; ∗∗p<0.01; ∗∗∗p<0.001 vs vehicle (b) or as depicted by lines (e, f). 2-way ANOVA was used for (b) and 1-way ANOVA was used for (e-f).

To test if systemic AM-3 application is also effective in the SNI model, we i.v. injected either relaxed *trans*-AM-3 or pre-illuminated *cis*-AM-3 at the dose of 5 mg/kg, and measured the PWL over 90 minutes. *cis*-AM-3, but not *trans*-AM-3, reversed the thermal hyperalgesia of the ipsilateral paw without affecting the contralateral one (**Fig. 5c; Extended Data Fig. 14b**). As a control we used the peripherally restricted MOR agonist loperamide^43,44^, which showed a similar effect to *cis*-AM-3 (**Fig. 5c; Extended Data Fig. 14b**). We next asked if AM-3 may be locally activated to its *cis* form *in vivo* following systemic treatment to promote a local anti-hyperalgesic effect. We i.v. injected 10 mg/kg *trans*-AM-3 and locally photoactivated to the *cis* form with 365 nm light illumination of the paw, which rapidly increased PWL that returned to baseline within 30 min (**Fig. 5d**). Subsequent 365 nm exposures produced similar anti-hyperalgesic effects, supporting the ability of repeated photoactivation of AM-3 to successfully provide robust analgesia.

To assess the ability of local, optically-targeted peripheral opioid action to drive analgesia with reduced gastrointestinal (GI) side effects, we assessed constipation by measuring faecal boli accumulation over 95 min and comparing it to loperamide^45,46^. Strikingly, despite a similar net analgesic effect for loperamide, systemic *cis*-AM-3, and local photoactivation of AM-3 (**Fig. 5e**), only loperamide or systemic *cis*-AM-3 produced substantial constipation (**Fig. 5f**). Together these data support the potential of AM-3 to enable local analgesia as a means of limiting both central and peripheral GI side effects.

## Discussion

Peripheral opioids are promising compounds for effective analgesia with a better safety profile than systemic opioids^7,47^. While chemical strategies have been reported to achieve peripheral restriction, here we use both chemical modification and light to target peripheral opioid analgesia, providing further precision to minimize off-target effects. The primary compound developed in this study, AM-3, shows minimal accumulation in the brain, minimal activity in the relaxed *trans* state, and robust analgesic effects in the active *cis* state following illumination. Importantly, and in contrast to caged ligand approaches^11^, AM-3 can be reversibly and repeatedly photo-activated and photo-deactivated, providing a high degree of spatiotemporal control both for basic and clinical application. Our work complements previous non-opioid azobenzene-based photopharmacological approaches to preclinical pain modulation^48–50^, supporting this novel mode of drug action as a promising alternative to traditional analgesic drugs.

We report two ways of applying AM-3 as a local analgesic: first, AM-3 may be “pre-activated” by illuminating prior to local application, as demonstrated with hindpaw injection. Local administration has the advantage of delivering high drug concentration at the site of injection and has previously been harnessed for topical opioid formulations^51,52^. In this mode of application, visible light illumination or thermal relaxation of azobenzenes can drive deactivation of AM-3 to further limit sustained or systemic MOR activation compared to classical opioids. Second, AM-3 can be applied systemically in its inactive *trans* form and locally activated via targeted illumination. While many peripheral sites are optically accessible, implant-based light delivery techniques may be required depending on the application. Notably, AM-3 should be well-suited for probing MOR signaling in the brain using optogenetic light fiber techniques following local delivery^53^.

We find that AM-3 serves as an efficacy switch such that *cis* and *trans* forms show comparable binding affinities for the MOR but *cis* is more effective as an agonist. While affinity switches represent the traditional mode of photochromic GPCR ligands^54–56^, they have emerged as an alternative mechanism^57–59^. Efficacy switches offer a clear advantage in terms of therapeutic window. Since optical illumination does not lead to 100% population of either *cis* or *trans* states, competition between the two forms will occur under most conditions. If *cis* had a substantially higher affinity than trans, at high doses strong effects would be seen following visible light illumination due to the 10-20% of molecules in the cis *state* which would outcompete the 80-90% in *trans*. Interestingly, despite clear partial agonism of *trans*-AM-3 in cultured cell experiments, no or minimal effects of *trans*-AM-3 were observed *in vivo*. A simple and plausible explanation is that the high expression seen in heterologous studies leads to an overestimation of partial agonism compared to *in vivo*, endogenous systems which may have a higher threshold for a biologically impactful activation. The absence of noticeable activity of *trans*-AM-3 *in vivo* could also be due, at least in part, to a pharmacokinetic effect. *Trans*-AM-3 is more lipophilic than its *cis* isomer and therefore more prone to be sequestered, e.g., by serum albumin^60^, and thus may be less bioavailable in the blood circulatory system.

We report cryo-EM structures of both *cis*- and *trans*-AM-3 bound to the MOR which provide a structural explanation for the efficacy differences between these states. In both forms, the naltrexone moiety binds in the canonical major pocket with a similar pose to prior structures. However, the azobenzene extension occupies the fentanyl subpocket in both *cis* and *trans*, with substantial differences which likely reposition key transmembrane helices, ultimately modulating conformational dynamics within the ternary complex, which results in low or high efficacy activation. Similarly to IBNtxA^26^, AM-3 is able to serve as an agonist despite the naltrexone antagonist scaffold, demonstrating the steep sensitivity of the MOR to subtle changes in ligand composition. Our functional and structural data support a simple model where the morphinan core serves to produce binding affinity while efficacy is controlled by the precise pose of the ligand in extracellular subpockets. Our structural analysis provides a framework for rational engineering of AM-3 variants with improved properties including red-shifted activation spectra, accelerated thermal relaxation, fine-tuned efficacy, opioid receptor subtype selectivity, and, possibly, G protein or arrestin bias.

## Methods

### Molecular docking

The active-state BU-72-bound structure of the µ-opioid receptor (PDB ID: 5C1M)^19^ was retrieved from the Protein Data Bank. The G-protein mimetic camelid antibody fragment was removed, retaining the receptor subunit, three crystallographic water molecules (wa* 8, 25, and 46), and the crystallized BU-72. Residues 52–63 at the N-terminus were deleted. Hydrogen atoms were added and optimized, and side-chain conformations were refined to improve structural accuracy. The inactive-state, beta-FNA-bound structure of the µ-opioid receptor (PDB ID: 4DKL)^18^ was similarly retrieved and processed retaining the receptor subunit, two crystallographic water molecules (wa* 18 and 19), and the crystallized ligand. Residues 263–269, which are part of the T4 lysozyme (T4L) fusion protein, were deleted. As with the active-state receptor, hydrogen atoms were added and optimized, and side-chain conformations were refined.

Prior to docking, the azo-morphine (AM-1, AM-2, AM-3, AM-4) in both *cis* and *trans* isomers of the azo group underwent chiral definition and formal charge assignment. Two-dimensional molecular representations were converted into three-dimensional models, which were energy optimized using the MMFF94 force field^61^. A biased-probability Monte Carlo (BPMC) optimization approach was applied for docking simulations, where compound conformations in internal coordinates were sampled using pre-calculated grid energy potentials of the receptor^62^. The grid potentials accounted for receptor limited flexibility through “soft” van der Waals potentials, while maintaining the receptor’s overall conformational state.

All-atom docking simulations were conducted with flexible models of azo-morphines in *cis* and *trans* isomers, with effort value of 5. To avoid major compound displacement for the suboptimal AM-x isomers, the common morphinan substructure from the naltrexone binding pose was used as a template, where the ligand atoms were tethered to the corresponding common substructure template atoms by soft harmonic restraints. The docking box was defined to encompass the extracellular half of the receptor. A minimum of 15 independent docking runs, each generating three top scoring conformations, were performed starting from random ligand conformations. The best ten docking results were analyzed for consistency by comparing ligand poses. Top-scoring docking solutions were refined iteratively using energy minimization and Monte Carlo sampling. Refinements focused on ligand conformations and receptor side chains within a 5 Å radius of the binding site. All molecular modeling operations were performed in the ICM-Pro v3.9-2b molecular modeling and drug discovery suite (Molsoft LLC).

### Chemical synthesis and characterization

See Supplementary Methods.

### Molecular biology

The rat SNAP-MOR (N-terminal HA followed by SNAP-tag followed by full length MOR) was previously described^63^. SNAP-tagged human MOR (cloned from addgene #66464), SNAP-tagged human KOR (Addgene #66462), SNAP-tagged human DOR (Addgene #66461) and SNAP-tagged mouse MOR (cloned from SSF-MOR, a gift from the Von Zastrow lab) were produced using Gibson Assembly Cloning Kits (NEB) by placing a SNAP tag between the signal sequence/FLAG tag and the N-terminus of the receptor. For SNAP-DOR, SNAP-KOR, and SNAP-MOR (human), the mGluR5 signal sequence was added and the C-terminal TANGO domain was removed. The previously described GIRK1-F137S hometetramerization mutant ^64^ was used for patch clamp recordings and the ONE-Go Gαi3 biosensor (Addgene kit #1000000224) was used for BRET measurements. For recombinant protein expression, a full length human MOR construct was subcloned into a modified pFastBac1 vector with an N-terminal HA-signal peptide, FLAG-tag, 10xHis-tag and thermostabilized b_562_RIL (BRIL) and a C-terminal TwinStrep tag and an additional 10xHis-tag. Both N- and C-terminal tags are cleavable by PreScission sites (pFastBac1-BRIL-hMOR).

### Cell culture and transfection

HEK 293 and HEK 293T cells purchased from ATCC (CRL-1573) and tested routinely for mycoplasma were used. Cells were cultured in Dulbecco’s modified Eagle’s medium (DMEM, Gibco) supplemented with 10% Fetal Bovine Serum (FBS) and maintained at 37°C in a 5% CO2 humidified incubator. For patch clamp and live cell imaging experiments, cells were seeded 24 hr prior to transfection on poly-L-lysine-coated glass coverslips (18 mm) in a 12-well plate. For BRET measurements, cells were seeded on a 10 cm non-treated tissue culture dish. Lipofectamine 2000 (Invitrogen) was used for transfection of DNA plasmids. For whole-cell patch clamp, cells were plated at low density and transfected with 0.7 μg SNAP-tagged mouse, rat, or human MOR, 0.7 μg GIRK1, and 0.15 μg tdTomato (as a transfection marker) per well. For cell surface labeling and internalization assays, cells were transfected with 0.35 μg rat SNAP-tagged MOR per well. For BRET measurements, cells were transfected with the following plasmids with 1 µg of the ONE-GO Gαi3 biosensor (1 µg) and 5 µg of SNAP-rMOR per dish.

### Surface labeling internalization assay and widefield fluorescence imaging

A previously described surface labeling-based internalization assay was used^63,65^. 24–36 hours post-transfection, coverslips containing cells expressing SNAP-rMOR were washed three times in 1.5 mL of extracellular (EX) solution composed of (in mM): 135 NaCl, 5.4 KCl, 10 HEPES, 2 CaCl2, and 1 MgCl2 (pH 7.4) and incubated in either no drug (media, control), 1 µM DAMGO (positive control), or *cis* or *trans* AM-3 (1 nM to 1 µM), in EX for 40-45 min at 37 degrees. AM-3 photoactivation was performed using a handheld 365 nm UV Flashlight Torch Light (HQRP 365 nm 12 LED). Cells were washed three times prior to labeling by incubation in 1 µM BG-Alexa-546 (New England BioLabs) in EX for 20 min at room temperature. Following fluorophore labeling, live cells were washed and imaged on an inverted microscope (Olympus IX83) with a 60x 1.49 NA objective. Alexa-546 was excited using a 560 nm laser and snapshots were acquired to measure the mean fluorescence intensity of labeled surface receptors using ImageJ. Fluorescence was normalized to the average fluorescence of the no drug control. Raw fluorescence values were normalized to 1 µM DAMGO for side-by-side comparison to the electrophysiology. The decrease in surface fluorescence was calculated as 100 * (1 - normalized surface fluorescence). Data was calculated across at least three experimental days per condition.

To observe the internalization of SNAP-tagged surface receptors, cells were incubated in 1 µM BG-Alexa-546 for 30-45 min at 37°C in EX. To induce internalization of SNAP-tagged receptors, cells were incubated with agonist in EX solution for 30 min at 37°C and imaged. Alexa-546 was excited with a 560 nm laser and snapshots of randomized fields were acquired. The basal condition was run side-by-side with drug conditions with the same allotted time but in EX solution.

### Patch clamp electrophysiology

Whole-cell patch-clamp recordings were performed as previously described^66^. 24-36 hours after transfection in a high potassium extracellular solution containing (mM): 120 KCl, 25 NaCl, 10 HEPES, 2 CaCl2, and 1 MgCl2 (pH 7.4). Solutions were delivered to a recording chamber using a gravity-driven perfusion system with exchange times of ∼1 s. Cells were voltage-clamped at −60 mV using an Axopatch 200B amplifier (Molecular Devices). Patch pipettes with resistances of 3-8 MΩ were filled with an intracellular solution containing (in mM): 140 KCl, 10 HEPES, 3 Na2ATP, 0.2 Na2GTP, 5 EGTA, 3 MgCl2 (pH 7.4). Currents were sampled at 10 kHz and filtered using a low-pass Bessel (8-pole) at 2 kHz. Illumination was applied to the entire field of view using a CoolLED pE-4000 through a 40x objective. Light intensity in the sample plane was 1-2 mW/mm². pCLAMP software was used for data acquisition, control of illumination, and data analysis. All responses were quantified relative to 1 μM DAMGO responses. Recordings were analyzed using Clampfit (Molecular Devices) and Prism (GraphPad) software. Data was calculated across at least three experimental days per condition.

### ONE-GO BRET biosensor measurements

8 hrs after transfection, media was replaced and cells were harvested 24–36 hrs later. Cells were centrifuged for 5 min at 550 x g, and resuspended in BRET buffer (140 mM NaCl, 5 mM KCl, 1 mM MgCl2, 1 mM CaCl2, 0.37 mM NaH2PO4, 20 mM HEPES, pH 7.4, 0.1% glucose) at a concentration of approximately 1 million cells/mL. Forty to fifty thousand cells were added to a white opaque 96-well plate (Corning) and mixed with NanoLuciferase substrate Nano-Glo (Promega) for 2 min before measuring luminescence in a Promega GloMax Discover plate reader at 28°C. Luminescence was measured at 450 BP (±10 nm) and 540 SP (±10 nm), and the BRET signal was calculated as the ratio between the emission intensity at 540 nm divided by the emission intensity at 450 nm. Reagents were added to the wells during live measurements by hand. AM-3 was preactivated with a handheld UV Flashlight Torch Light (HQRP 365 nm 12 LED). BRET data are presented as the difference from baseline BRET signal (ΔBRET) by subtracting the average of the signal pre-stimulation from all data points. The BRET signal (540 nm luminescence/450 nm luminescence) was measured every minute for 3-5 min with a signal integration time of 0.3 s for each measurement. Where indicated, the EC50 and Emax values were determined using a 3-parameter sigmoidal curve-fit in Prism (GraphPad). Data was calculated across at least three experimental days per condition.

### MOR expression and purification

Recombinant expression of the pFastBac-BRIL-MOR construct bound to either *cis* or *trans* AM-3 was performed in Spodoptera frugiperda (Sf9) insect cells using the Bac-to-Bac expression system (Gibco). Briefly, 1% (v/v) baculovirus was transduced into Sf9 cells at a density of 4 x 10^6^ cells/mL in ESF921 medium, containing 1% (v/v) production boost additive. 10 µM naloxone was maintained in the insect cell culture to improve receptor surface expression. Following incubation for 48 h at 27 °C, cells were harvested, and resuspended in a hypotonic buffer containing 10mM HEPES (pH 7.5), 10mM MgCl₂, 20mM KCl, and an in-house protease inhibitor cocktail (2mM AEBSF, 14μM E-64, 1μM leupeptin, and 0.3μM aprotinin). The cell suspension was lysed using a glass homogenizer and then subjected to further homogenization in a hypertonic buffer (hypotonic buffer supplemented with 1M NaCl). The resulting mixture was centrifuged at 175,000 x g for 45 minutes at 4°C. This process was repeated twice, with the final two washes supplemented with 10μM of either *cis* or *trans* AM-3. Following the washes, membranes were diluted in buffer containing 40mM HEPES (pH 7.5), 100mM NaCl, 10 μM AM-3, 10% glycerol, 0.5% (w/v) LMNG (Anatrace), 0.05% (w/v) CHS (Anatrace), and 2 mg/mL iodoacetamide, and incubated at 4°C for 14 h. The insoluble fraction was cleared by centrifugation at 175,000 g for 1 h. MOR was purified using Strep-Tactin®XT 4Flow® resin (IBA), in buffer containing 40 mM HEPES pH 7.5, 100 mM NaCl, 5 mM MgCl2, 3 mM CaCl2, 0.1 mM TCEP, 0.001% (w/v) LMNG, 0.0001% (w/v) CHS, 5% glycerol and 10 μM AM-3, supplemented with 50 mM Biotin (IBA).

### Expression and purification of heterotrimeric G proteins

Wild-type Gαi1 was co-expressed with Gβ1 and Gγ2 subunits in Sf9 cells at a density of 2 × 10⁶ cells/mL in ESF921 medium, using P2 baculovirus at a multiplicity of infection (MOI) ratio of 10:5. After 48 hours of incubation at 27°C, cells were harvested by centrifugation, washed with ice-cold PBS, and stored at −80°C until further processing. Purification of Gαi(wt)βγ / Gαi(DN)βγ heterotrimer was performed following established protocols (Rasmussen, S. G. F. et al, 2011). Briefly, cells were thawed, homogenized once in hypotonic buffer supplemented with 10 µM GDP and 5 mM β-mercaptoethanol, and centrifuged at 100,000 x g. The pellet was then solubilized in a buffer containing 20 mM HEPES (pH 7.5), 100 mM NaCl, 1% sodium cholate, 0.05% DDM, 5 mM MgCl₂, 5 mM β-mercaptoethanol, 15 mM imidazole, 10 µM GDP, and protease inhibitor for 90 minutes at 4°C. Insoluble material was removed by centrifugation at 150,000 x g for 45 minutes at 4°C. The supernatant was applied to Ni-NTA resin in the same buffer, and the protein was eluted with 300 mM imidazole. The eluate was concentrated using a 50-kDa cutoff concentrator (Amicon) and further purified via anion exchange chromatography on a 1 mL HiTrap Q FF column (Cytiva).

### Cryo-EM sample preparation and data collection

MOR, purified in the presence of either 10 µM *cis* or *trans* AM-3, was incubated with Gαiβγ at a 1:1.5 molar ratio for 2 h at room temperature. Following the formation of complexes, GDP hydrolysis was catalyzed via the addition of 2 units of apyrase (NEB) and a further incubation for 2 h. Uncomplexed G protein was separated by an additional purification step with M2 anti-FLAG resin (Sigma Aldrich). The resulting eluate was subjected to size exclusion chromatography using a Superdex 200 10/300 column in 40 mM HEPES pH 7.5, 100 mM NaCl, 3 mM MgCl2, 0.1 mM TCEP, 5 μM AM-3, 0.00075% LMNG, 0.000075% CHS and 0.00025% GDN buffer. Peak fractions were concentrated and immediately deployed for downstream cryo-EM studies. MOR:Gi-wt complexes were concentrated to ∼1 mg/mL using a 50-kDa cutoff concentrator (Amicon). Immediately after, 3 µL of the purified complex was applied to glow-discharged UltrAuFoil 1.2/1.3 300-mesh grids. Grids were blotted for 1-2 seconds in 95% relative humidity at 4°C, then rapidly vitrified in liquid ethane using a Vitrobot Mark IV (Thermo Fisher). Cryo-EM data collection was performed on a Titan Krios (Thermo Fisher) at 300 keV, using an aberration-free image shift (AFIS) data collection scheme, and a K3 direct-electron detector coupled with a BioQuantum energy filter (Gatan), set to a 20 eV slit width. Four images per hole were acquired with EPU data acquisition software (version 2.0). Each image was captured with a total exposure time of 1.8 seconds, a cumulative dose of 51 e−/Å² (cis-AM-3) and 55 e−/Å² (trans-AM-3), and a defocus range of −1 µm to −3 µm.

### Single particle cryo-EM image processing

All data processing was performed using the software package cryoSPARC^67^ (v4.5.3; Structura Biotechnology) (**Extended Data Fig. 8-9, Extended Data Table 1**). A total of 13,418 and 28,180 micrographs were collected for *cis*-AM-3 and *trans*-AM-3 respectively. Motion correction was performed on raw micrographs using Patch Motion Correction, followed by CTF estimation. Micrographs with CTF estimates worse than 3.5 Å were excluded from further processing. Automated particle picking was performed using a reference-free blob picker, and particles were extracted with a box size of 128 pixels (bin=4, 512 pixels uncropped box size), followed by three rounds of 2D classification. Ab initio reconstruction was performed using a “clean” stack of 1,142,073 particles for *cis*-, and 1,329,432 particles for *trans*-AM-3, resulting in a 3D volume with clearly visible densities for all components of the MOR-Gi complex (Extended Fig. 8-9). Particles were then re-extracted using a box size of 256 pixels (bin=2, 512 pixels uncropped box size) for *cis-* and 384 pixels (bin=1.33, 512 pixels uncropped box size) for *trans-*, followed by additional 2D classification. Final particle sets comprising 580,551 particles for *cis*- and 628,564 particles for *trans*-AM-3 were deployed for further downstream processing.

For the *cis*-AM-3 MOR:Gi complex, the initial ab initio reconstruction was subjected to multiple rounds of non-uniform refinement (low pass filter = 10 Å), interspersed with local refinements resulting in a density map with resolution estimates of 3 Å. Next, focused refinements were performed using manually created masks around the TMD, masking the micelle (‘Mask 1’) and heterotrimeric G protein (‘Mask 2’), yielding focused maps with resolutions between 2.9 Å −3.1 Å. These were combined in Chimera, for subsequent model building and refinement. Finally, 3D Variability Analysis^33^ (**Fig. 3g**; **Extended Data Fig. 8**) was performed on the final particle set (filter resolution = 3.3 Å), and the resulting principal components of motion were visualized in chimera.

Processing of the *trans*-AM-3 MOR-Gi density map followed initial non-uniform and local refinement steps. While these refinements produced maps with strong and unambiguous density for the naltrexone scaffold, the *trans*-azobenzene group appeared highly dynamic. To address this, the particle stack underwent two rounds of 3D variability analysis^33^ (filter resolution = 3.3 Å), focusing on particles contributing to frames displaying prominent density for the trans-azobenzene group. The first round of 3DVA was performed on the stack of 628,564 particles.

Frames 5–14 from principal component (PC) 2 were selected, resulting in a subset of 511,627 particles. *Ab initio* reconstruction was performed on this subset, followed by non-uniform and local refinements. These steps resulted in the appearance of the *trans*-azobenzene group, however it remained insufficiently resolved. A second round of 3DVA^33^ was subsequently conducted, and frames 9–13 from PC2 were selected, reducing the particle set to 284,756 particles (**Fig. 3h**, **Extended Data Fig. 9**). *Ab initio* reconstruction, followed by non-uniform and local refinements, yielded maps with unambiguous density for the trans-azobenzene group.

Finally, focused refinements were performed using manually created masks. Mask 1 targeted the transmembrane domain (TMD) while excluding the micelle, and Mask 2 did the same for the heterotrimeric G protein (**Fig. 3h**, **Extended Data Fig. 9)**. The focused maps were finally combined in Chimera for subsequent model building and refinement.

### Radioligand binding

Radioligand competition assays were performed using membrane fractions prepared from HEK293F cells transiently expressing wild-type MOR. To prepare membranes, cells transfected with 1 µg/mL plasmid were first harvested and resuspended in hypotonic buffer (10 mM HEPES pH 7.5, 10 mM MgCl_2_, 20 mM KCl supplemented with 2 mM AEBSF, 14 μM E-64, 1 μM leupeptin and 0.3 μM aprotinin). Resuspended cells were then dounce homogenized and centrifuged at 175,000 g in two rounds to clear soluble fractions. Protein concentration of the membrane pellets was typically determined to be 4 mg/mL using a Bradford assay (Pierce), and aliquots were flash frozen in liquid nitrogen, and stored at −80°C until further use.

Competition binding assays were setup in 96-well plates containing membrane fractions diluted to 0.15 mg/mL, along with [^3^H]-Naltrexone at 2 nM, and an AM-3 dose (25 µM - 6.25 pM), all prepared in binding buffer (10 mM HEPES, 10 mM MgCl2, 20 mM KCl, 0.1% BSA and 100 µM Bacitracin). Competition reactions were incubated for 2 h in the dark, and terminated by vacuum filtration onto cold 0.3 % PEI soaked GF/A filters, followed by three rounds of washing with cold 50 mM HEPES (pH 7.50). Counts were read using a Microbeta2 plate reader (PerkinElmer) for one minute per well. Results were analyzed in GraphPad Prism 10.1.1, and the inhibitor constant (Ki) was determined using the one site - Fit Ki model equation: logEC50=log(10^logKi*(1+RadioligandNM/HotKdNM)), Y=Bottom + (Top-Bottom)/(1+10^(X-LogEC50)).

### Mice

All experiments used male C57BL/6J (Jackson Laboratory; 20–30g) or CD-1 (Charles River, 25– 40 g) mice, between 8 and 13 weeks old, housed 4 or 5 per cage under a 12-hour light–dark cycle (lights off at 19:00 and on at 7:00) with ad libitum access to food and water. Mice were kept at a constant temperature (20–24°C) and relative humidity (40–50%). All experiments involving animals were approved and performed according to Weill Cornell Medicine Institutional Animal Care & Use Committee (IACUC) guidelines under protocol 2017-0023 and followed ethical guidelines of IASP for investigation of experimental pain in conscious animals. C57BL/6J mice were used for tests following local intraplantar administrations, whereas CD-1 mice were used for tests following systemic intravenous (tail vein) administrations.

### *In vivo* AM-3 and light exposure protocols

AM-3 was dissolved in a vehicle consisting of 0.36, 1.44, or 2.88% DMSO in sterile 0.9% NaCl for intraplantar (i.pl.) injections. Morphine sulfate (Sigma-Aldrich, Inc., St. Louis, MO, US) was dissolved in the vehicle and injected i.pl. at 5 µg/5 µL or intravenously (i.v., tail vein) at 5 mg/kg. Naloxone hydrochloride (20 µg/5 µL, i.pl.; Tocris, Minneapolis, MN, US) was dissolved in sterile 0.9% NaCl. All drugs were i.pl. injected in a final volume of 5 µL. For intravenous (i.v.) tail vein injections, AM-3, loperamide (1 mg/kg; Tocris, Minneapolis, MN, US), or morphine (5 mg/kg) was dissolved in 10% DMSO in 0.9% NaCl. The injected volume was 5 ml/kg as a bolus injection.

For pre-illumination experiments, AM-3 or vehicle was exposed to 365 nm UV light (EA-140, 4 Watt, 120 V, 60 Hz, 0.2 mA; Spectro-UV, Farmingdale, NY, US) for 10 minutes prior to injections, under dark conditions using a far-red lamp (BlockBlueLight, North Charleston, SC, US). *In vivo* photoswitching experiments involved post-injection exposure of mice to 365 nm UV light for a total of 5 minutes (single or repeated exposure) or to blue 465 nm light (10 Hz, 10 ms pulses, 68 pulses per sequence, 90 sequences; Doric Lenses, Quebec, Canada) for a total of 3 minutes or as otherwise specified. All injections and light exposures were conducted under dark conditions using a far-red lamp. For *in vivo* photoswitching with alternating UV and blue light illumination (Fig. 5b), mice were exposed to 460 nm light for 2 minutes, tested 4 minutes after switching the light off, then re-exposed to 365 nm light for 3 minutes, retested after 4 minutes, and the cycle was repeated.

For structural studies, purification of MOR in the presence of cis-AM-3 or trans-AM-3 was performed in the dark or in ambient light, respectively. In each case, AM-3 was exposed to either 360 nm UV or blue 465 nm light for 30 minutes prior to addition, followed by periodic exposure of the sample, to the respective light.

### Measurement of analgesia and pain models

#### Hargreaves thermal plantar test

Mice were habituated to a Hargreaves apparatus (Ugo Basile, Varese, Italy) on tempered glass maintained at ∼28°C for 90–120 minutes per day over 2 days before behavioral testing. On test days, thermal nociceptive thresholds were recorded following habituation by measuring paw withdrawal latencies (PWL) to an infrared light source (800–1200 nm) focused on the plantar surface of each hind paw. After determining baseline PWLs, mice received vehicle or drug administrations. For opioid blockade, mice were pre-injected with naloxone or saline 10 minutes prior to AM-3 or vehicle administration. Recorded withdrawal behaviors included paw withdrawal, licking, biting, or shaking of the targeted hind paw. Measurements were repeated 1– 3 times per hind paw, with at least 2-minute intervals, and averaged for each paw. The intensity of the light source was adjusted to produce baseline responses averaging ∼6 seconds under both acute and post-SNI (spared nerve injury) surgery conditions, with a cut-off of 20 seconds to avoid tissue damage. To reduce experimental variability due to potential increases in paw skin temperature during photoswitching experiments ^68,69^, glass temperature was monitored to prevent increases exceeding +1–1.5°C.

#### Tail flick test

Analgesia was assessed using the Ugo Basile 37360 tail flick apparatus. For baseline measurements, mice were gently wrapped, and the lower one-third of the tail was positioned over the sensor emitting radiant heat via an infrared light source (800–1200 nm). The response latency was measured by the tail flick, with a maximal latency of 20 seconds set to avoid tissue damage. Following baseline recording, mice received an i.v. injection of vehicle or cis-AM-3 at doses of 2.5, 5, or 10 mg/kg (pre-illuminated). Analgesic response was tested every 5 minutes for 60 minutes, followed by assessments at 90 and 95 minutes post-administration.

#### Spared nerve injury model

Spared nerve injury was performed according to the method of Decosterd and Woolf ^70^. Under isoflurane anesthesia, the sciatic nerve was exposed, identifying the 3 peripheral branches (sural, common peroneal, and tibial nerves), and both tibial and common peroneal nerves were ligated and transected together. Animals recovered for 3 weeks after surgery. Thermal hyperalgesia was absent in healthy (pre-surgery) animals, and the paw withdrawal threshold in mice before SNI (pre-surgery) was very close to the set cut-offs.

### Locomotor activity

Activity was measured in a 50 cm x 50 cm black paved open field arena with 35 cm high walls. Locomotor activity was measured at 5-minute intervals, with cumulative counts taken for data analysis; counts represented the total number of beam breaks per 5-minute increment, capturing all movements, including running and turning behaviors ^71,72^. To evaluate the effects of AM-3 or morphine on locomotor behavior, mice were habituated in the activity monitor for 120 minutes. Immediately after the habituation period, mice were injected with drug or vehicle and returned to the monitor, where locomotor activity was recorded for the following 90 minutes. All analyses were conducted using EthoVision software (v.17.5, Noldus, Leesburg, VA, US).

### Gastrointestinal motility

The constipative effect of AM-3 and loperamide was assessed by measuring total accumulated fecal boli, as previously described^73^. Briefly, SNI mice were injected with drugs or vehicle and placed in a Plexiglass chamber (5 cm × 8 cm × 8 cm) positioned on a mesh screen. Mice had ad libitum access to food and water before testing. Fecal boli were collected and weighed 95 minutes post-administration.

### P-glycoprotein transporter substrate analysis

The permeability test was performed as previously described^74,75^. MDCKII-MDR1 cells were seeded at 1.56 × 10⁶ cells/mL in 96-well HTS Transwell plates (50 µL per well) and cultured at 37 °C, 5% CO₂, and 95% relative humidity for 3–8 days, with medium replaced every other day. Before experiments, plates were washed twice with pre-warmed HBSS (10 mM HEPES, pH 7.4) and incubated at 37 °C for 30 minutes. Test compounds (10 µM) were prepared by diluting 2 mM DMSO stock solutions in HBSS (final DMSO concentration: 0.5%). For apical-to-basolateral (A→B) transport, 125 µL of the compound solution was added to the apical compartment, with 50 µL immediately transferred to acetonitrile containing internal standards (IS: 100 nM ketoprofen, 200 nM labetalol, and 100 nM tolbutamide). For basolateral-to-apical (B→A) transport, 285 µL of the compound solution was added to the basolateral compartment, and 50 µL was sampled similarly. Receiver compartments were filled with transport buffer (1 μM for control compound or 5 μM for test compound diluted in 0.5% DMSO in 10 mM HEPES, pH 7.4). Plates were incubated at 37 °C for 2 hours. Post-incubation, 50 µL samples were collected from donor and receiver compartments, mixed with acetonitrile containing IS, vortexed (10 min), and centrifuged (3,220 g, 40 min). Supernatants (100 µL) were diluted with ultra-pure water for LC-MS/MS analysis.

### Data Analyses and Statistics

All results are presented as mean ± s.e.m. *In vivo* data were tested for normality and sphericity and a Greenhouse Geisser correction was applied where appropriate. Assumption of Normality was tested either with the Shapiro–Wilk test or with a Q-Q plot. Two-way ANOVAs were used with treatment and time as variables. Tuckey’s was used for *post hoc* comparisons of groups. GraphPad Prism 9.5 was used for data treatment and statistical analysis.

## Supporting information

SupplementaryMovie1_cisAM3_3DVA

SupplementaryMovie2_cisAM3_3DVA

## Data availability

The final cryo-EM maps for MOR:Gαi:*cis*-AM-3 and MOR:Gαi:*trans*-AM-3 have been deposited in the Electron Microscopy Data Bank under accession code: EMD-XXXXX and EMD-XXXXX. Corresponding atomic coordinates have been deposited in the PDB under accession code: PDB ID XXXX and PDB ID XXXX. Source data are provided with this paper.

## Acknowledgments

We thank Francis Lee (Weill Cornell) for access to the tail flick apparatus, Chris Mason (Weill Cornell) for plate reader access, and Nigel Liverton (Tri-Institutional Therapeutic Discovery Institute) for assistance with *in vitro* permeability measurements. We acknowledge the Center of Excellence for NanoImaging (CNI) at the University of Southern California for cryo-electron microscopy time. We thank H. Khant for assistance with cryo-EM data collection. We acknowledge the Center for Advanced Research Computing (CARC) for support with high performance computing resources. This research was supported by US National Institutes of Health grants R66/R33DA051529 (J.L., D.T., V.K.), R01AT012075 (C.G.), R01GM144965 (C.G.), R35GM153437 (V.K.) and T32GM132081 (G.R.), as well as the Rohr Family Research Scholar Award (J.L.) and the Monique Weill-Caulier Award (J.L). L.P., G.R., X.J., and S.K. contributed equally and all have the right to list themself first in bibliographic documents.

---

**Extended Data Figure 1.**
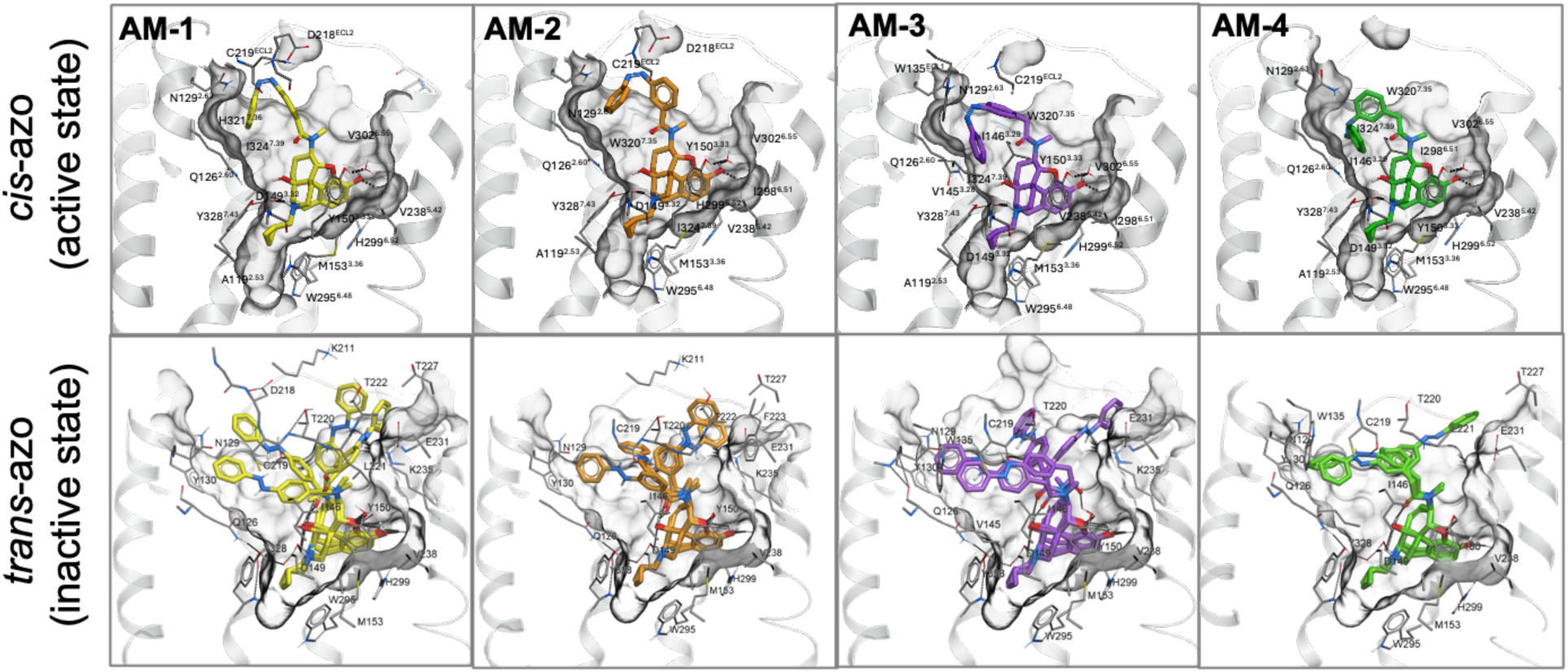
Molecular docking analysis of azo-morphines. Docking results of **AM1-4** analogs in *cis*- and *trans* isoforms to the MOR. The *cis*-azo isomer is shown by one conformation, consistently reproduced in 15 parallel docking simulations. The *trans*-azo isomer is shown by several examples of top-scoring conformations with similar scores, observed in docking simulations. The receptor models are shown in grey, with residues of the pocket in stick and surface presentations. The ligands are shown as sticks with carbons colored yellow, orange, magenta or green for each ligand, and other atoms colored by atom type (red: oxygen, blue: nitrogen). See Methods for details of docking simulations.

**Extended Data Figure 2.**
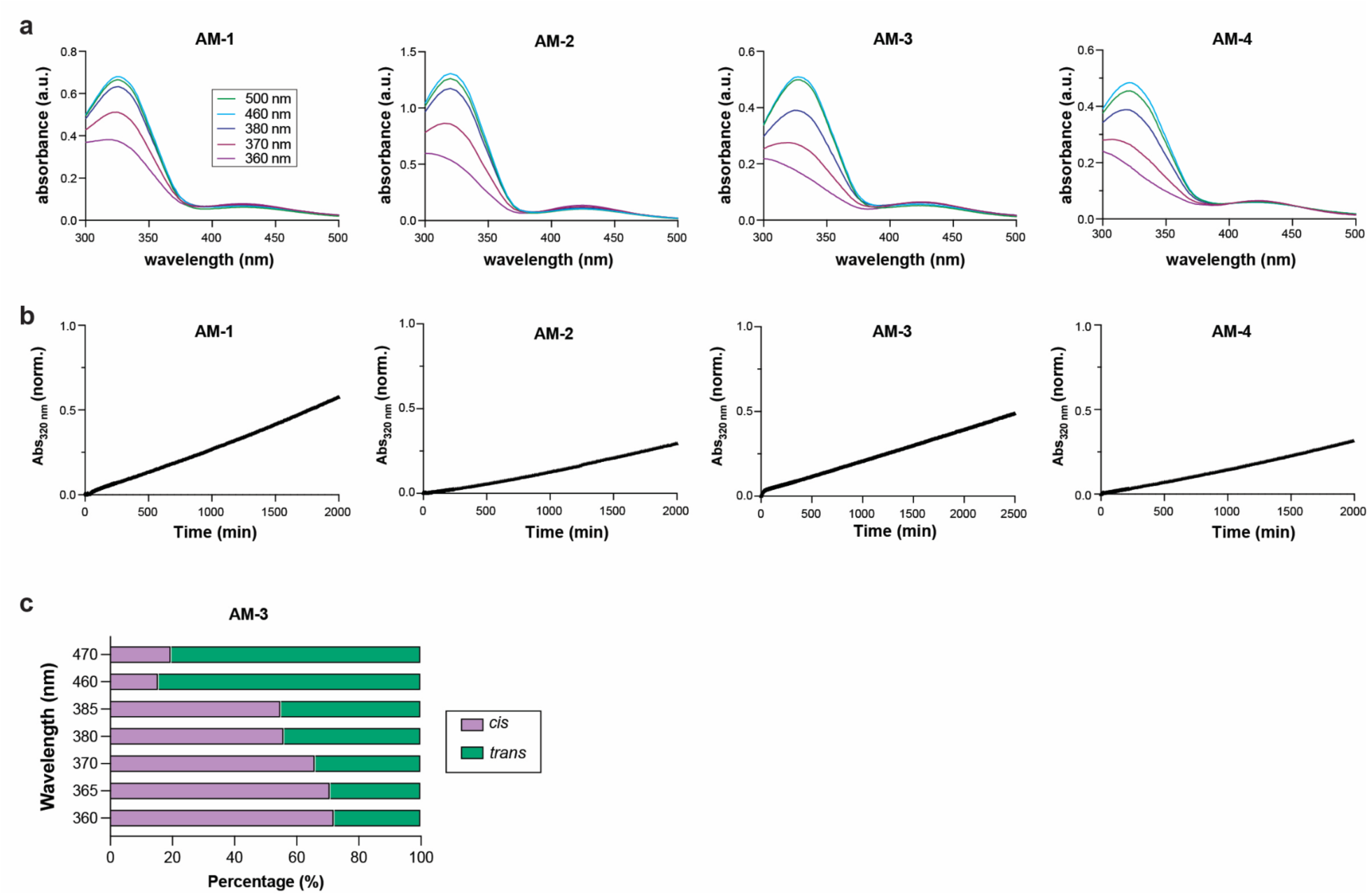
Photophysical characterization of azo-morphines. **a,** UV/vis spectra showing similar light sensitivity across azo-morphine variants with maximal *cis* content following 360 nm illumination and maximal *trans* content following >460 nm illumination. **b,** Time course of thermal relaxation to the *trans* state following 365 nm illumination to drive *cis* enrichment. **c,** Photostationary state measurements of relative *cis* and *trans* proportions of AM-3 following illumination with different wavelengths. Note: data in a and b were collected in PBS:DMSO = 9:1; data in c was collected in acetonitrile/H_2_O mixture.

**Extended Data Figure 3.**
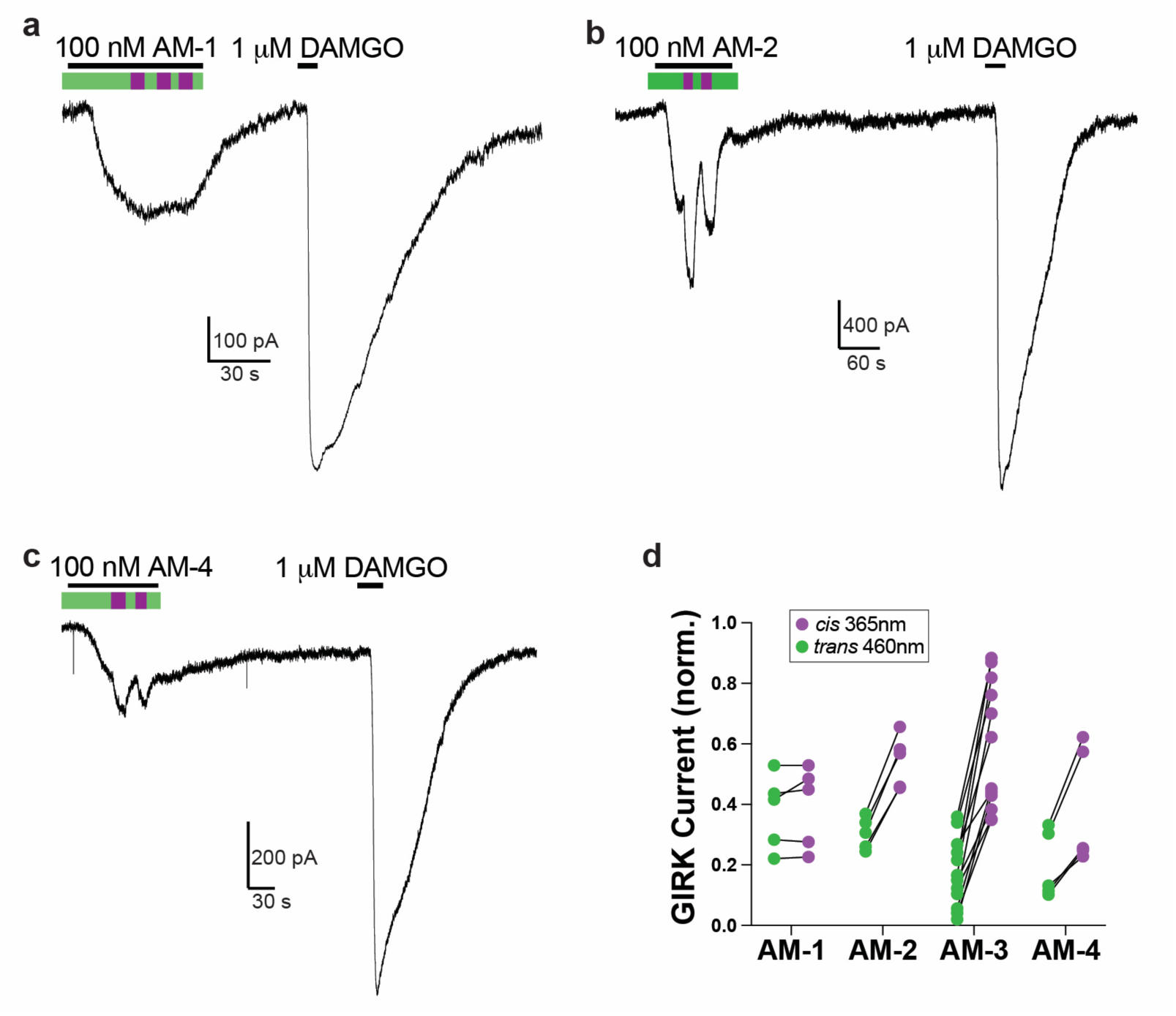
Further comparative electrophysiological analysis of azo-morphines. **a-d,** Representative GIRK current traces (a-c) and summary graph (d) showing different light sensitivities of azo-morphine variants on the rat MOR. In (d), each pair of points represents an individual cell and 100 nM azo-morphine was used.

**Extended Data Figure 4.**
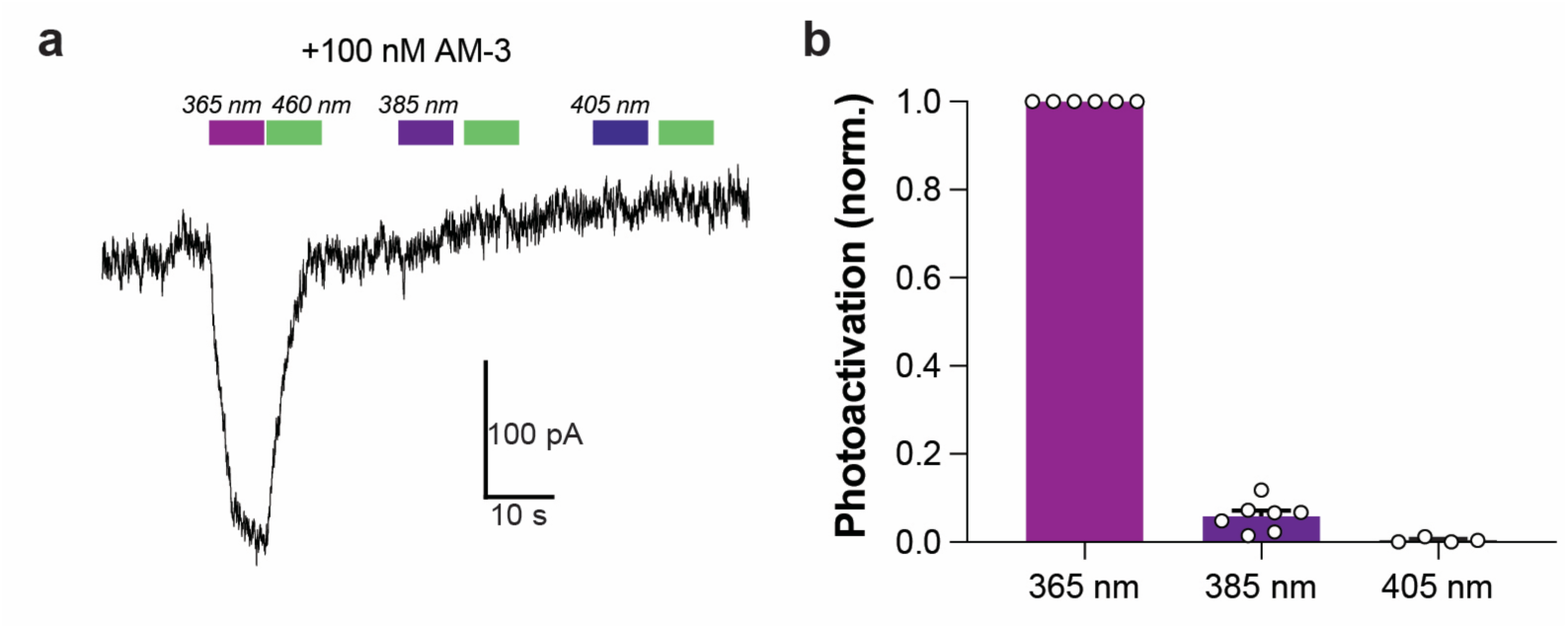
Analysis of wavelength-dependence of AM-3 photo-activation. **a-b,** Representative GIRK current trace (a) and summary bar graph (b) showing wavelength-dependence of AM-3 photoactivation. Each point represents an individual cell.

**Extended Data Figure 5.**
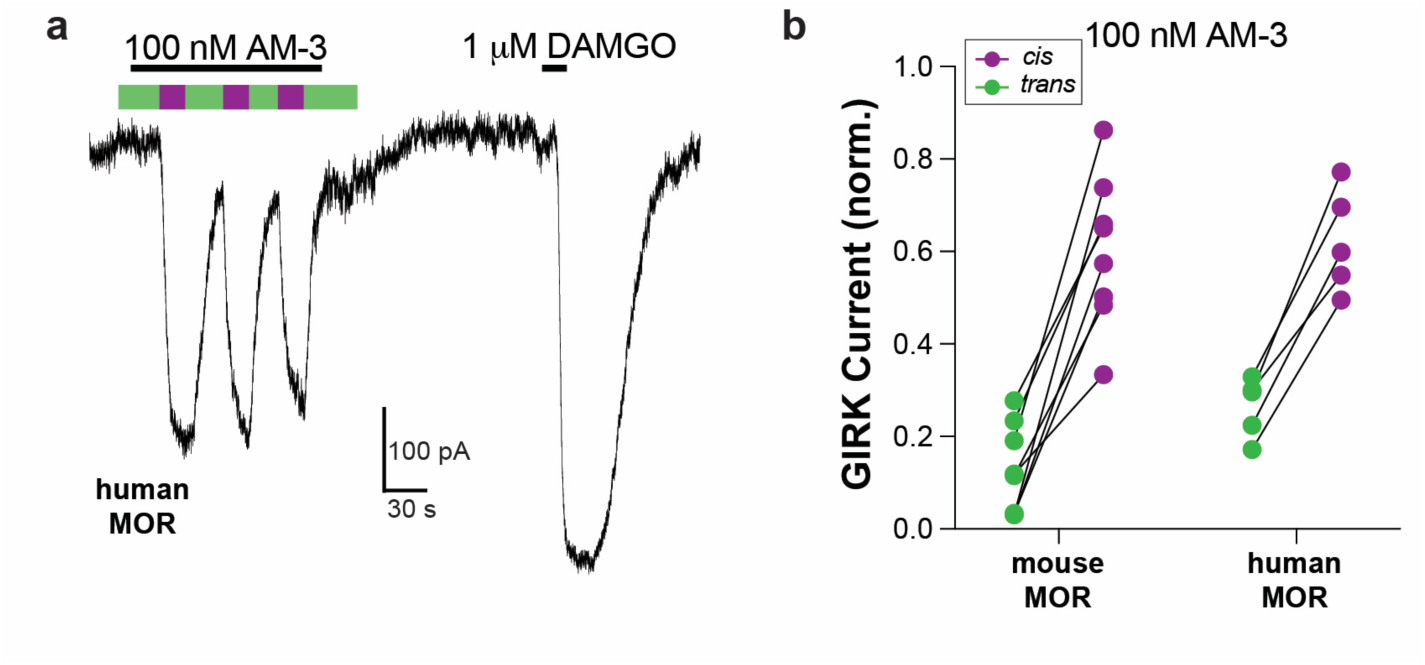
AM-3 photoactivation of mouse and human mu-opioid receptor. **a,** Representative GIRK current trace showing light dependent AM-3 activation of the human MOR. **b,** Summary graph showing stronger activation of mouse and human MOR via *cis*-AM-3. In (d), each pair of points represents an individual cell and current values are normalized to the response to 1 µM DAMGO.

**Extended Data Figure 6.**
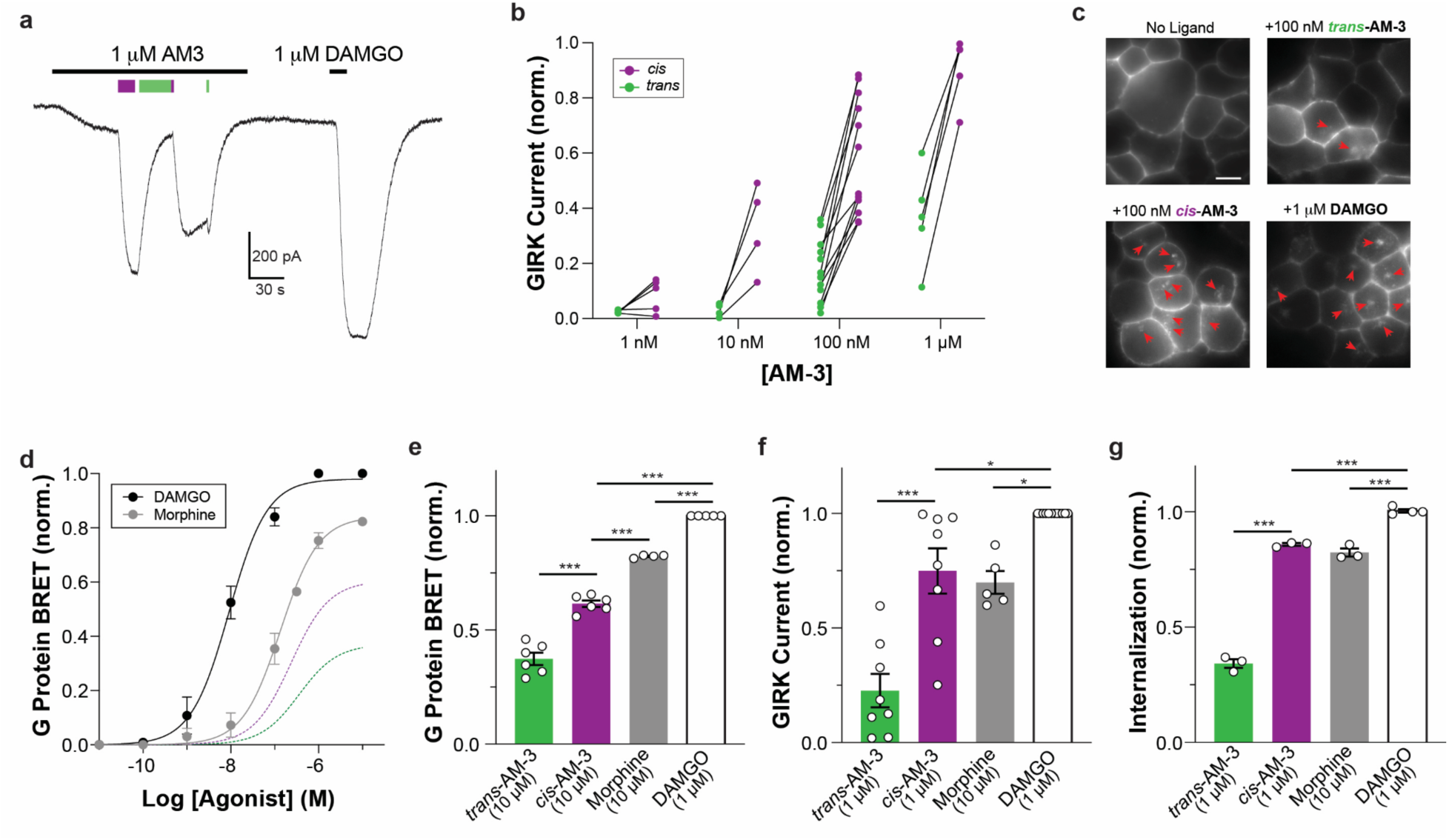
Further analysis of dose-dependence of AM-3 MOR activation. **a,** Representative GIRK current trace showing photoactivation of MOR via 1 uM AM-3. **b,** Summary graph showing relative activation of MOR by *cis* and *trans*-AM-3 across a range of concentrations. **c,** Representative images of cells showing that *cis*-AM-3 and DAMGO produce substantially more intracellular puncta (red arrowheads) compared to *trans*-AM-3 or no ligand treatment. Cells were labeled with BG-Alexa-546 to visualize SNAP-tagged MOR and then treated with ligands for 30 minutes prior to imaging. **d,** ONE-GO G protein BRET Dose response curves for DAMGO and morphine showing relative potency and efficacy compared to *cis* (purple dotted line) and *trans*-AM-3 (green dotted line). **e-g,** Summary bar graphs showing relative to maximal response to ligands across assays. All data are shown as mean±s.e.m. Points represent independent experimental days for (e) and (g) and individual cells for (b, f). ∗p<0.05; ∗∗∗p<0.001 vs vehicle. 1-way ANOVA was used.

**Extended Data Figure 7.**
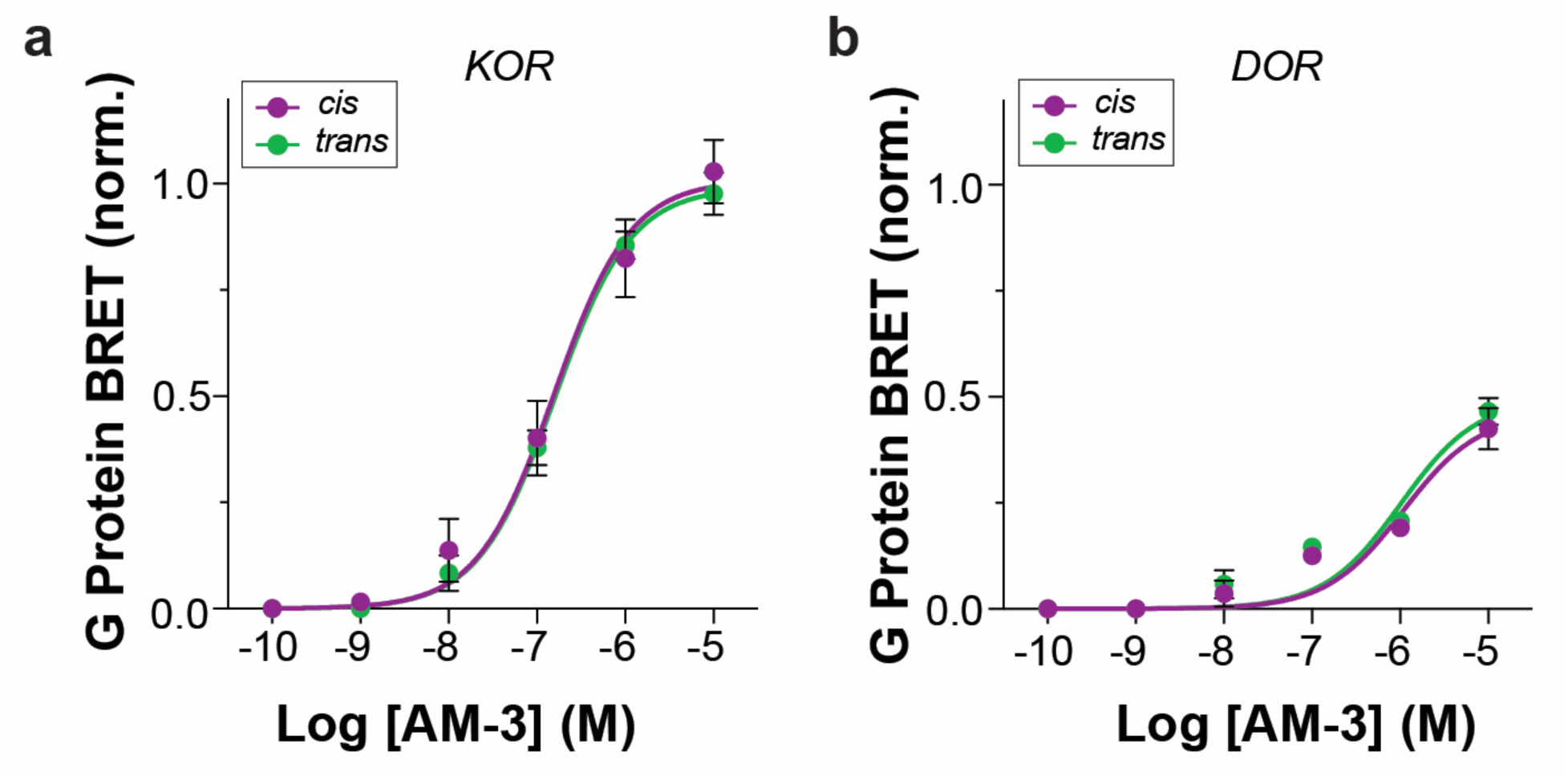
Lack of light-dependence of activation of KOR and DOR by AM3. **a-b,** ONE-GO G protein BRET dose response curves showing similar activation of KOR and DOR by *cis* versus *trans*-AM-3. All data are shown as mean±s.e.m.

**Extended Data Table 1.** Cryo-EM statistics.

|  |  |  |
| --- | --- | --- |
| **Structure** | MOR-*cis*-AM-3:Gi | MOR-*trans*-AM-3:Gi |
| Accession Codes | EMD-48249<br>PDB: 9MGC | EMD-48250<br>PDB:9MGD |
| **Data collection and processing** |  |  |
| Magnification | 130,000 | 130,000 |
| Voltage (kV) | 300 | 300 |
| Electron exposure (e<sup>-</sup>/Å<sup>2</sup>) | 51 | 55 |
| Defocus range (μm) | -1.0 - -3.0 | -1.0 - -3.0 |
| Pixel size (Å) | 0.647 | 0.647 |
| Symmetry | C1 | C1 |
| Micrographs (No.) | 13,148 | 28,180 |
| Initial particles (No.) | 5,179,391 | 8,335,494 |
| Final particles (No.) | 580,551 | 284,756 |
| Map resolution (Å) | 3.2 | 3.1 |
| FSC threshold | 0.143 | 0.143 |
| **Refinement** |  |  |
| Initial model | 8EF5 | 8EF5 |
| Model resolution (Å) | 3.2 | 3.2 |
| FSC threshold | 0.5 | 0.5 |
| Map sharpening B factor (Å<sup>2</sup>) | 110.9 | 94.4 |
| **Model composition** |  |  |
| Non-hydrogen atoms | 7,104 | 7,236 |
| Protein atoms | 900 | 899 |
| Ligands | 1 | 1 |
| Other | 3 | 4 |
| **B-factors (Å<sup>2</sup>)** |  |  |
| Protein | 39.6 | 52.8 |
| Ligand | 50.9 | 23.5 |
| Other | 51.4 | 67.2 |
| **R.m.s deviations** |  |  |
| Bond length (Å) | 0.004 | 0.004 |
| Bond angle (°) | 0.699 | 0.728 |
| **Validation** |  |  |
| MolProbity Score | 1.44 | 1.40 |
| Clash Score | 4.94 | 3.38 |
| Poor rotamers | 0.0 | 0.0 |
| **Ramachandran plot** |  |  |
| Favored (%) | 96.97 | 96.06 |
| Allowed (%) | 3.03 | 3.94 |
| Disallowed (%) | 0.0 | 0.0 |

**Extended Data Figure 8.**
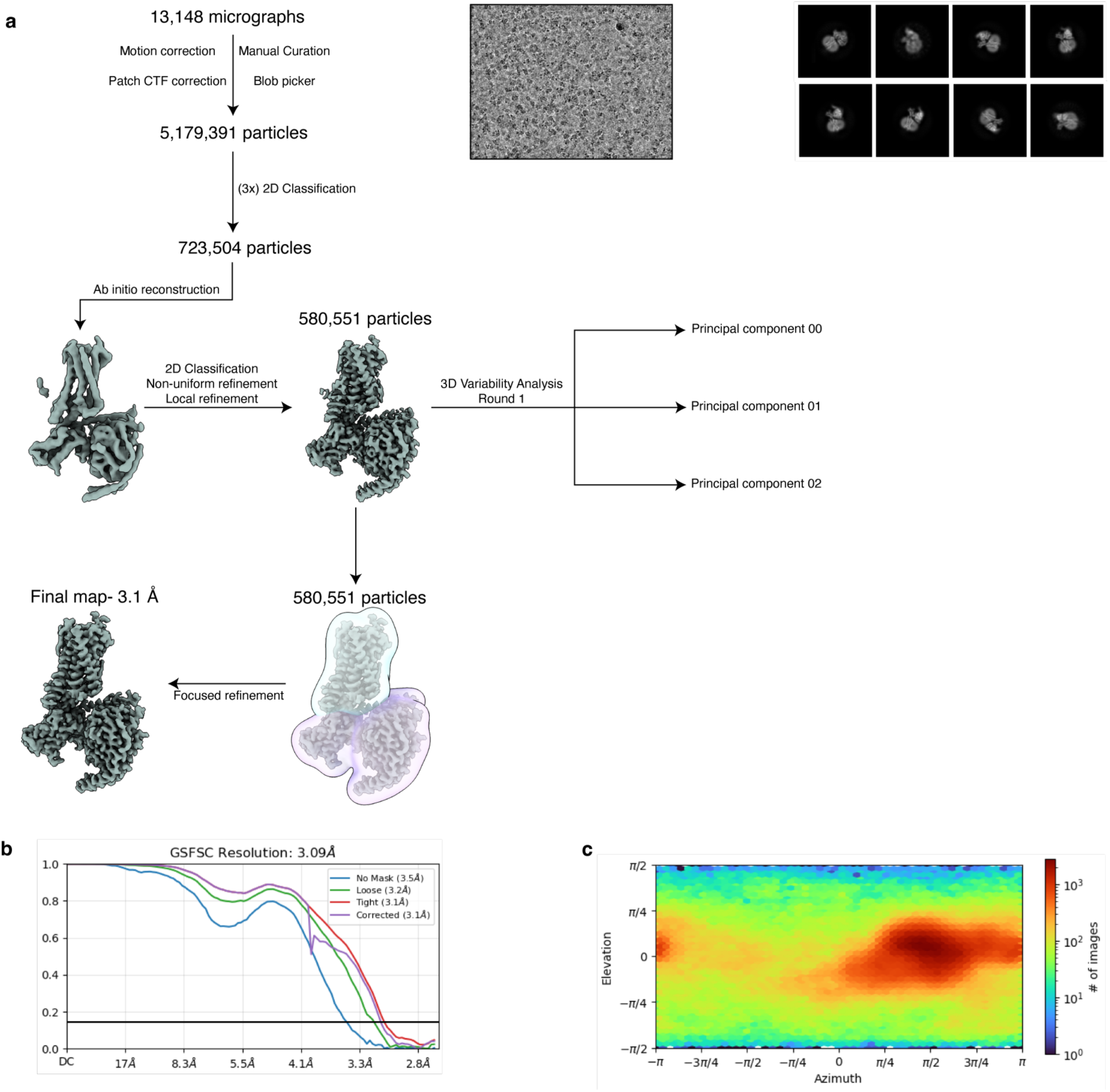
Data processing pipeline for the *cis*-AM-3 MOR:Gi complex. **a,** Workflow depicting cryo-EM processing the *cis*-AM-3 MOR:Gi complex with representative micrograph and 2D classes. Data processing was performed in cryoSPARC^67^ using established workflows. Following motion correction, CTF estimation, and particle picking, the dataset was cleaned using 2D classification. Ab initio reconstruction was performed to obtain a 3D volume, which was subjected to non-uniform and local refinement resulting in a density map with unambiguous density for *cis*-AM-3. 3DVA was performed on the final particle set to determine conformational dynamics within the MOR:Gi complex. **b-c,** Data quality of the final reconstruction, shown as a (b) gold-standard Fourier shell correlation plot (masked and unmasked) and (c) angular sampling distribution.

**Extended Data Figure 9.**
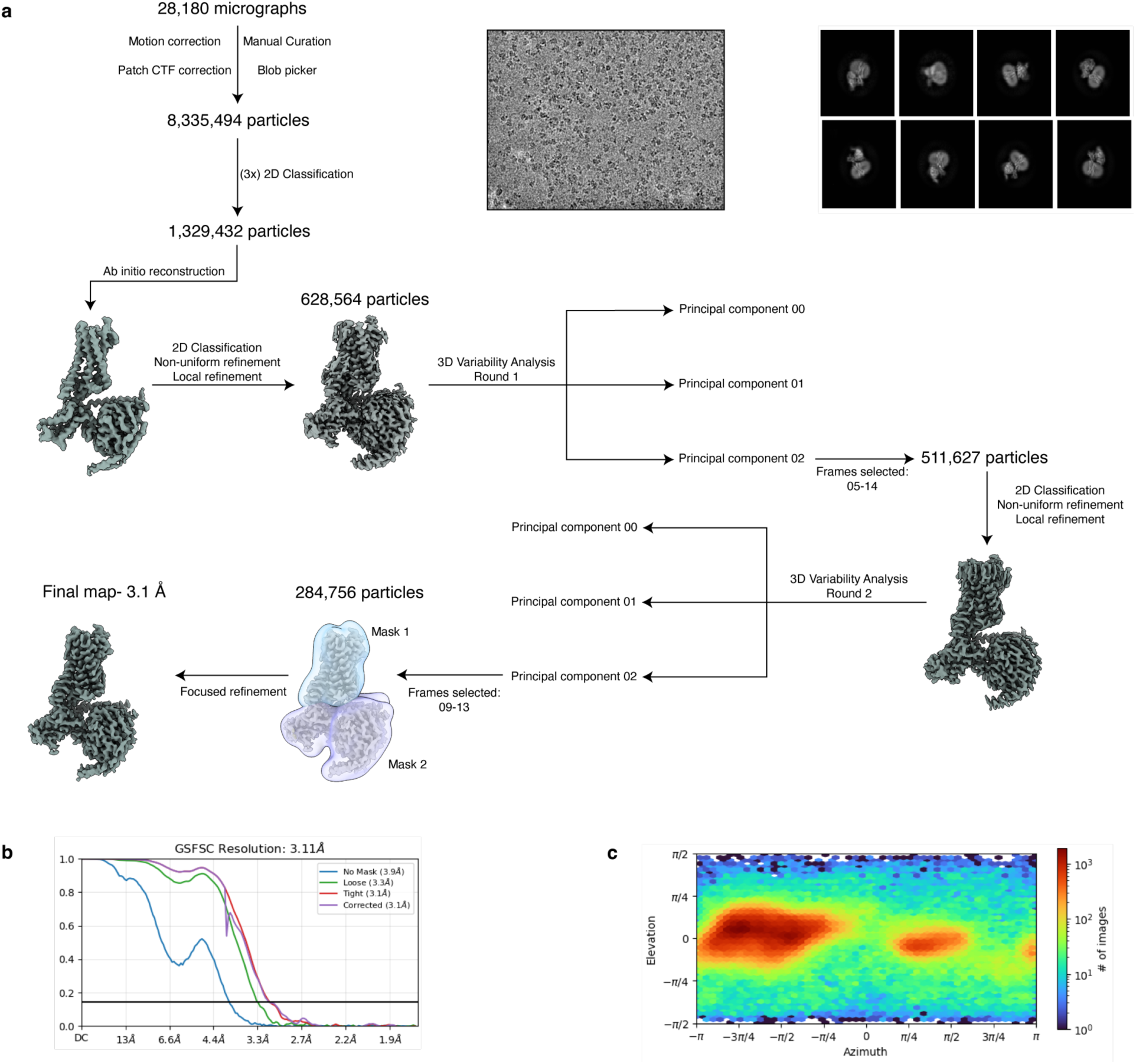
Data processing pipeline for the *trans*-AM-3 MOR:Gi complex. **a,** Workflow depicting cryo-EM processing the *trans*-AM-3 MOR:Gi complex with representative micrograph and 2D classes. Data processing was performed in cryoSPARC^67^. Following motion correction, CTF estimation, and particle picking, the dataset was cleaned using 2D classification and 3D ab initio reconstruction. Initial refinements revealed strong density for the naltrexone scaffold but dynamic density for the trans-azobenzene group. Two rounds of 3DVA were performed to select particles contributing to frames with improved *trans*-azobenzene density, followed by focused refinements. **b-c,** Data quality of the final reconstruction, shown as a (b) gold-standard Fourier shell correlation plot (masked and unmasked) and (c) angular sampling distribution.

**Extended Data Figure 10.**
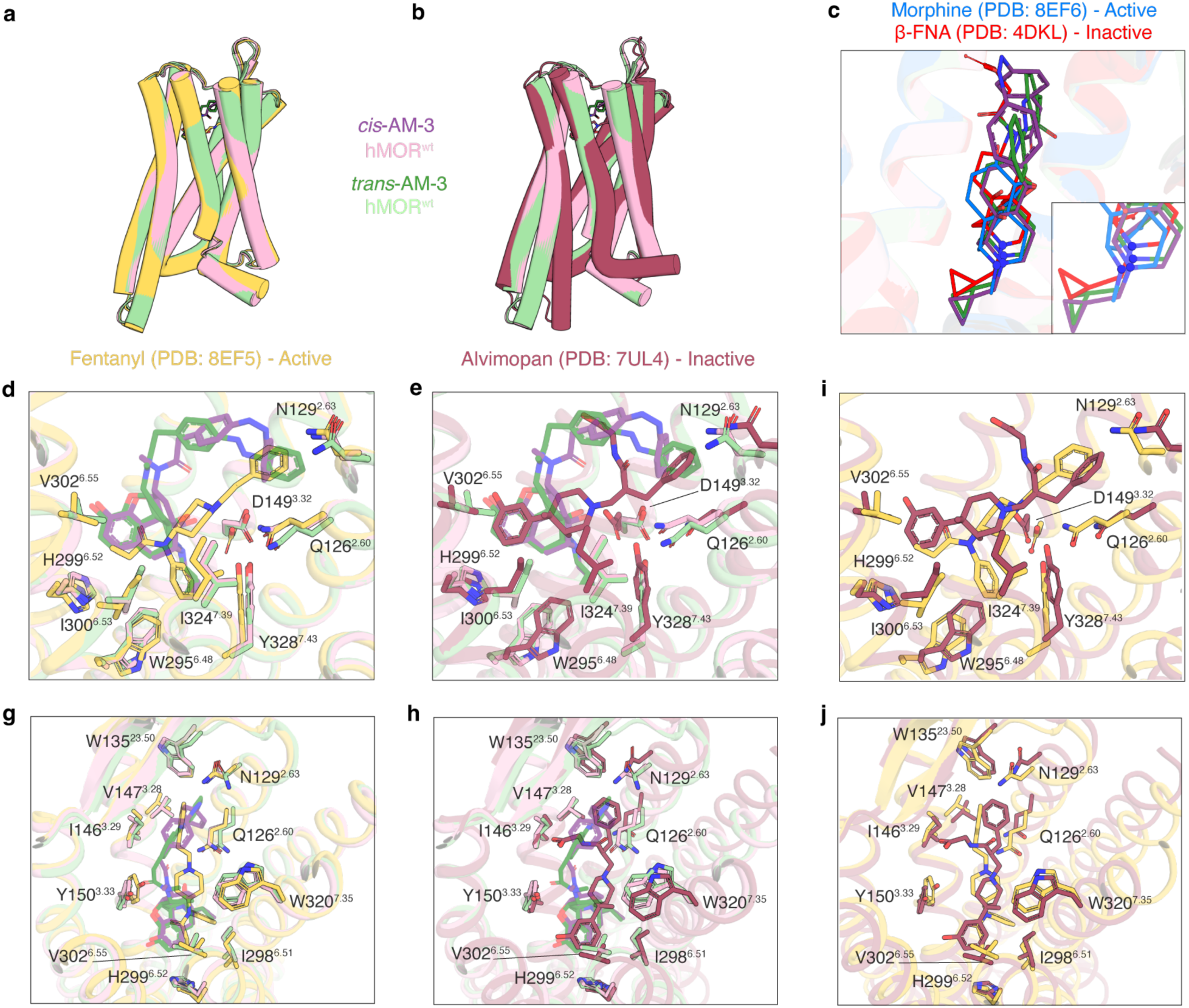
Structural comparison of MOR bound to AM-3, fentanyl, and alvimopan. **a-b,** Model overlays of *cis-* (light green) and *trans-*AM-3 (pink) bound MOR to the (a) active state fentanyl-bound (PDB:8EF5, yellow), and (b) inactive state alvimopan-bound MOR (PDB:7UL4, maroon). **c,** Alignments of *cis*- and *trans*-AM-3 to morphine (PDB:8EF6, blue) and β-FNA (PDB:4DKL) highlighting the position of the morphinan tertiary amine. Deeper penetration into the orthosteric site correlates with ligand efficacy. **d-j,** Detailed comparisons of the *cis-* and *trans-*AM-3 bound orthosteric site of MOR to fentanyl (d,i,g,j) and alvimopan (e,h,i,j) showing overlapping occupation of the subpocket through the terminal azo-benzene of AM-3, and the phenyl and benzyl groups of fentanyl and alvimopan, respectively.

**Extended Data Figure 11.**
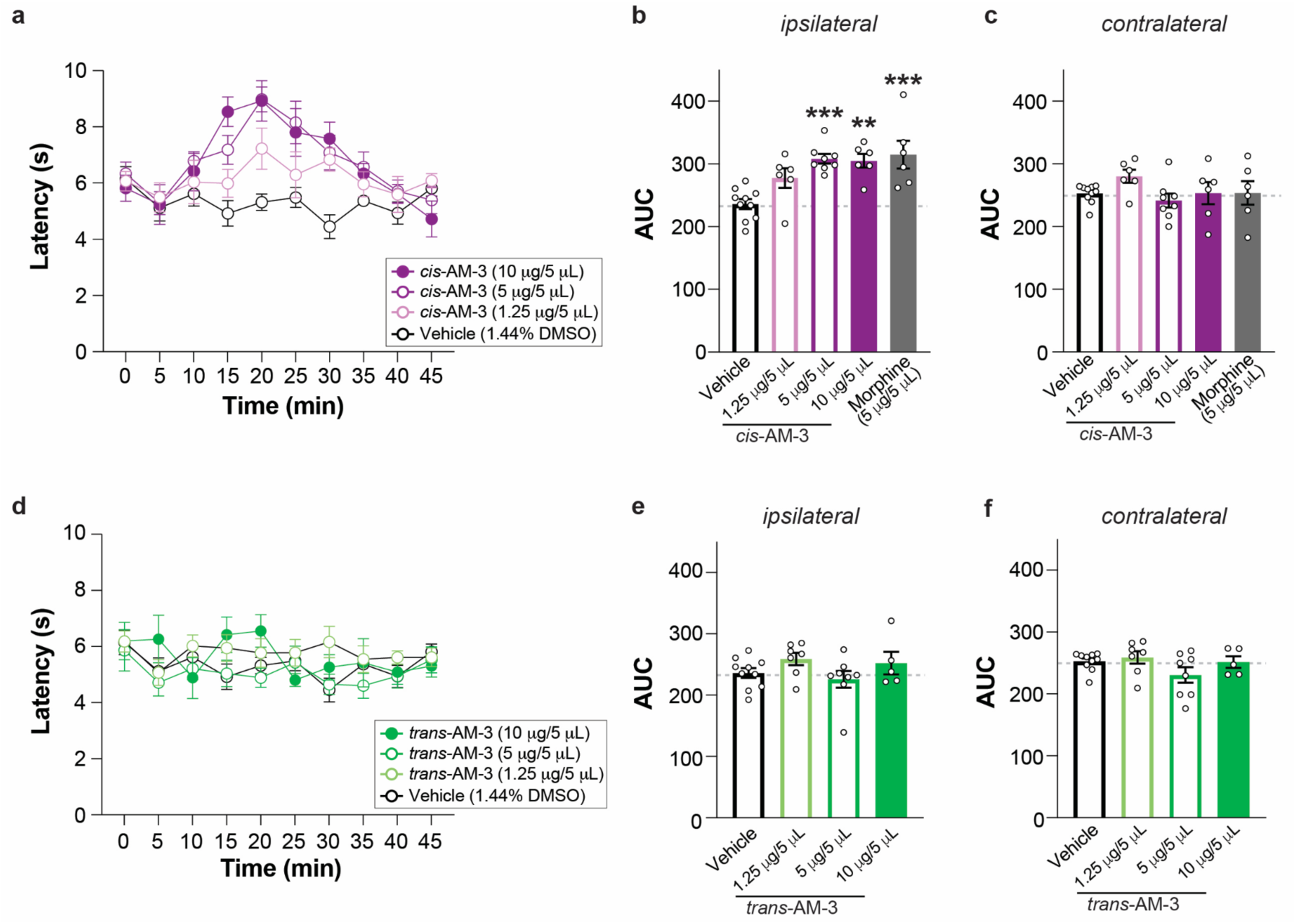
Dose-dependence of antinociceptive effects of AM-3. **a,** Time course of PWL of cis-AM-3 across doses. **b-c,** Summary bar graphs of area under curve showing analgesic effect of cis-AM-3 and morphine in the ipsilateral, injected paw (b), but not the contralateral paw (c). **d-f,** Same as a-c but for trans-AM-3. All data are shown as mean±s.e.m. with n>6 mice/condition (a-c) and n>5 (d-f); for (b, c, e, f) points represent individual mice; ∗∗p<0.01; ∗∗∗p<0.001 vs vehicle. 1-way ANOVA was used for (b, c, e, f).

**Extended Data Figure 12.**
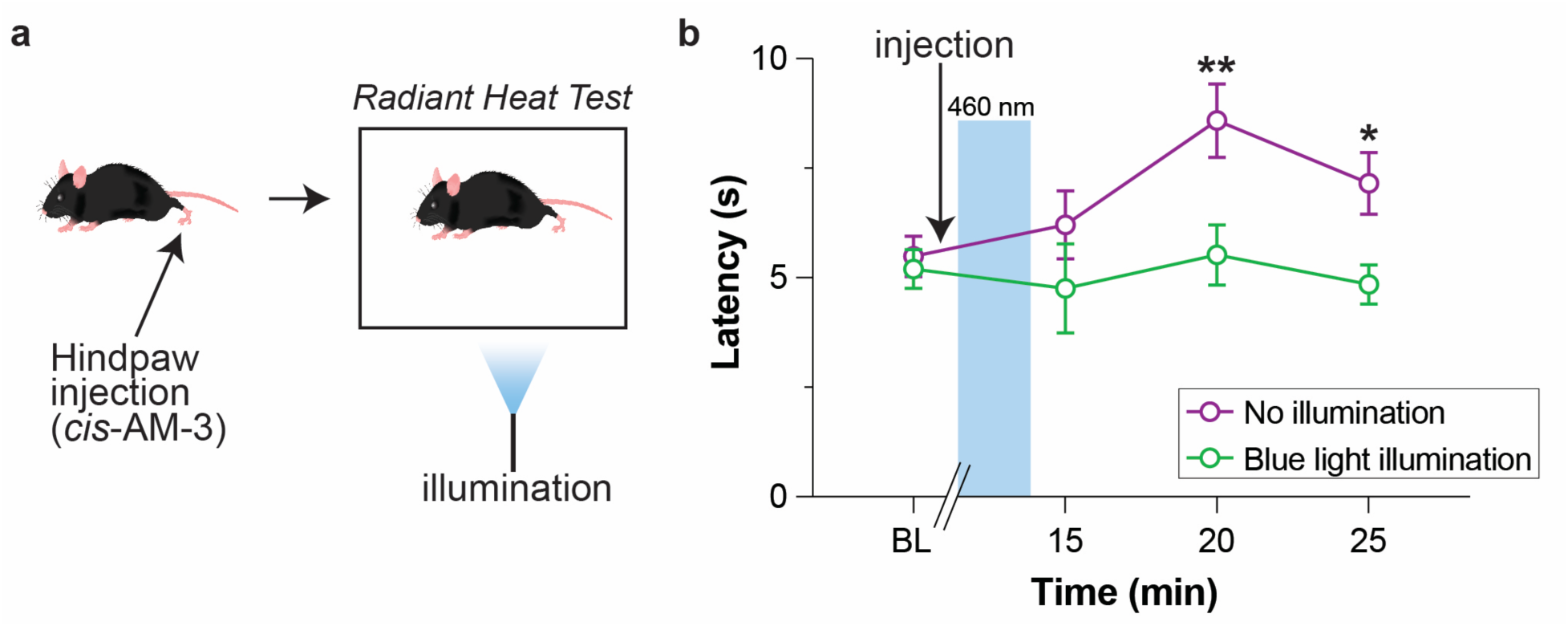
Photo-deactivation of AM-3 *in vivo*. **a-b,** Schematic (a) and time course (b) showing reversal of the PWL effect of cis-AM-3 via 460 nm illumination to drive conversion to trans-AM-3. All data are shown as mean±s.e.m. with n>5 mice/condition; ∗p<0.05; ∗∗p<0.01; vs blue light illumination. 2-way ANOVA was used for (b).

**Extended Data Figure 13.**
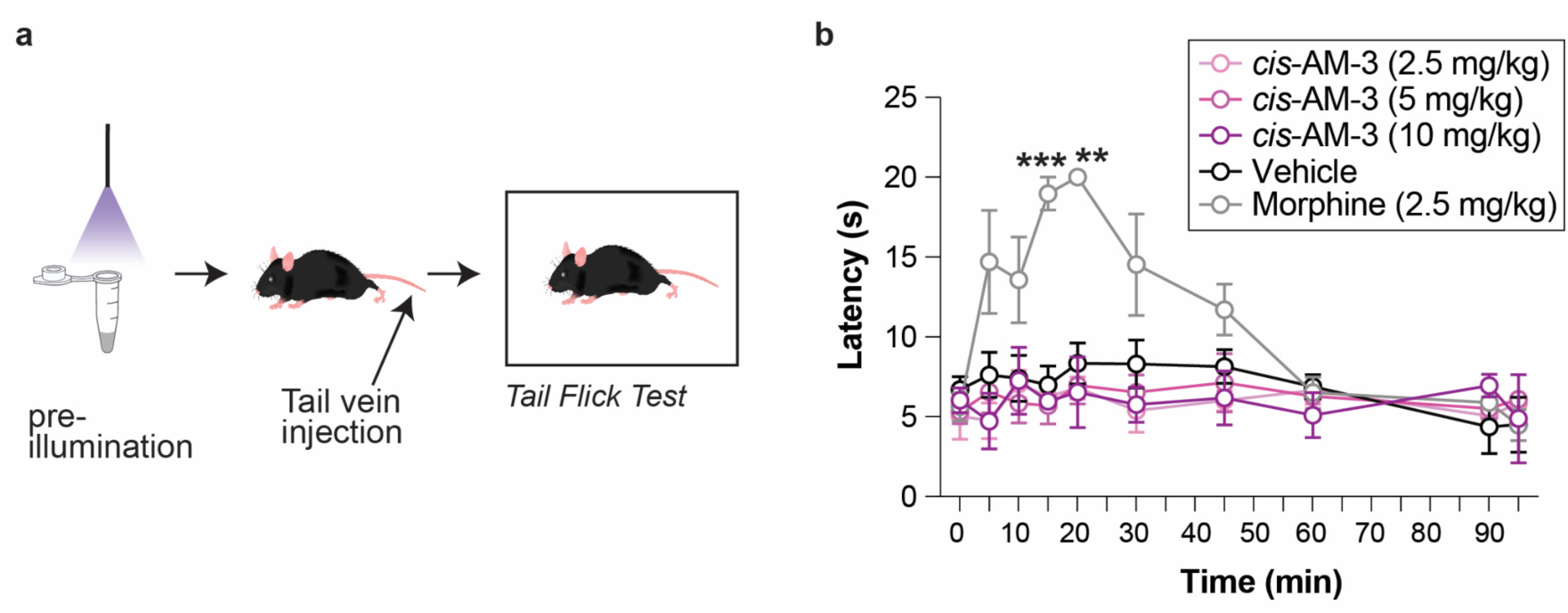
Lack of effect of intra-venous AM-3 in the tail flick test. , Schematic (a) and time course (b) showing lack of spinal analgesic effect of 2.5, 5, and 10 mg/kg i.v. cis-AM-3 in the tail flick test. All data are shown as mean±s.e.m. with n>4 mice/condition; ∗∗p<0.01; ∗∗∗p<0.001 vs vehicle. 2-way ANOVA was used for (b).

**Extended Data Table 2.** Permeability analysis of AM-3 in a MDCKII-MDR1 cell line assay.

| Compound | P<sub>app</sub> <sub>(A-B)</sub> (10<sup>-6</sup>, cm/s) | P<sub>app</sub> <sub>(B-A)</sub> (10<sup>-6</sup>, cm/s) | Efflux Ratio | Recovery (%) A-B | Recovery (%) B-A |
| --- | --- | --- | --- | --- | --- |
| Metoprolol | 24.24 | 27.42 | 1.14 | 102.95 | 99.01 |
| Digoxin | 0.57 | 11.80 | 20.68 | 106.08 | 118.28 |
| *cis*-AM-3 | 2.28 | 16.43 | 7.32 | 59.44 | 75.47 |
| *trans*-AM-3 | 2.58 | 16.96 | 6.67 | 62.83 | 79.25 |

The efflux ratios for cis- and trans-AM-3 were compared to standard compound Metoprolol (non-substrate for efflux transporters) and Digoxin (substrate for efflux transporters) were used as controls to validate the assay.

Apparent permeability (Papp) was calculated for drug transport assays using the equation:

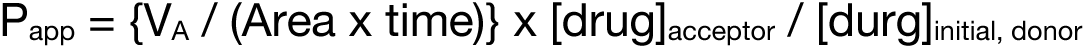

where P_app_ is apparent permeability (cm/s x 10^−6^), V_A_ is the volume (in mL) in the acceptor well, Area is the surface area of the membrane (0.143 cm^2^ for Transwell-96 Well Permeable Supports) time is the total transport time in seconds.

Efflux ratio was determined using the following equation: Efflux Ratio = P_app_(B-A)/P_app_(A-B), where P_app_(B-A) indicates the apparent permeability coefficient in basolateral to apical direction, and P_app_(A-B) indicates the apparent permeability coefficient in apical to basolateral direction.

Mass balance (% recovery) was determined using the following equation:

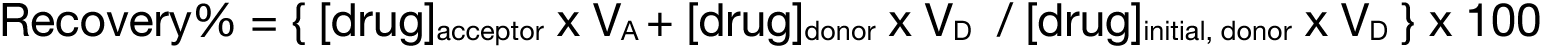

where V_A_ and V_D_ are the acceptor and donor well volumes, respectively (0.235 mL for A→B flux, 0.075 mL for B→A flux).

**Extended Data Figure 14.**
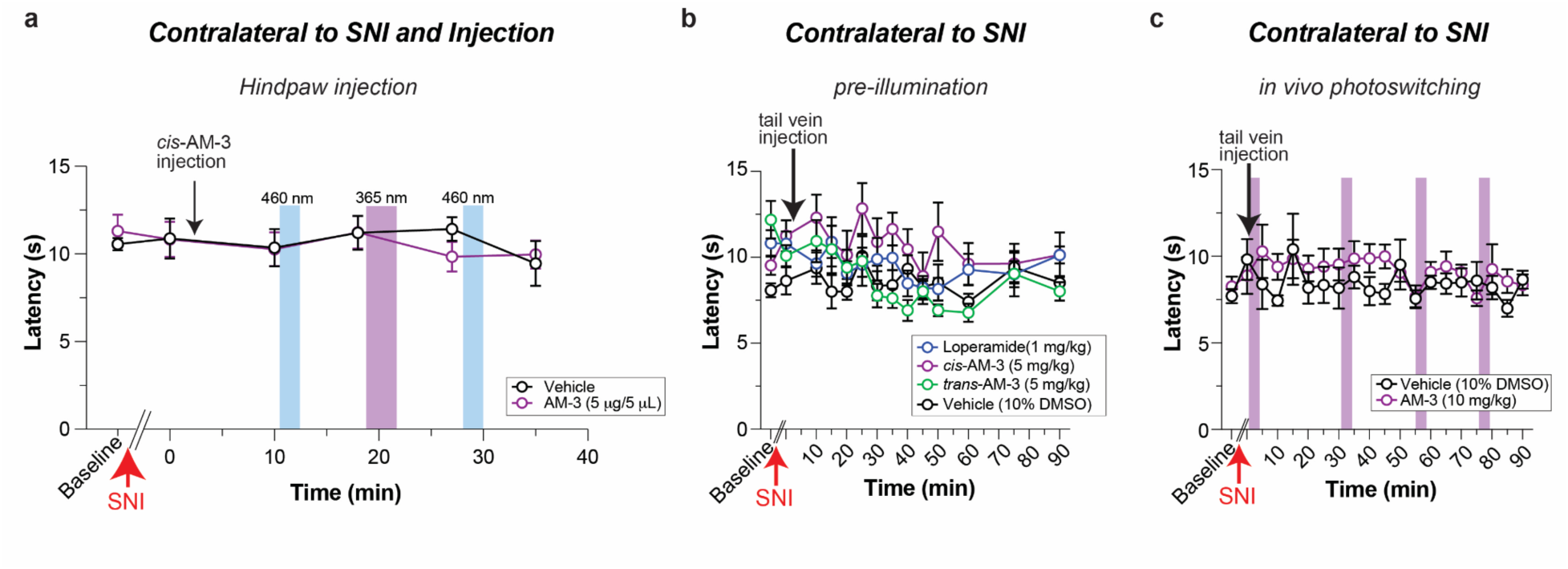
Lack of effect of AM-3 in the contralateral paw in the SNI neuropathic pain model. **a-c,** Time courses showing lack of antihyperalgesic effect of AM-3 in the contralateral paw in SNI mice following ipsilateral hindpaw (a) or tail vein injection (b, c). All data are shown as mean±s.e.m. with n≥5 mice/condition.

