## Supplementary Information for "An opioid efficacy switch for reversible optical control of peripheral analgesia"

### Synthesis

**Methods and Chemicals**: Unless otherwise stated, all reactions were performed with magnetic stirring under a positive pressure of nitrogen or argon gas. Solvents and reagents were used as received from commercial sources (Sigma-Aldrich, Alfa Aesar, Acros Organics, Strem Chemicals, Combi-Blocks, Oakwood, Toronto Research Chemicals). Solvents were purchased from Fisher and Acros Organics. The anhydrous solvents were either purified using a Pure Process Technologies solvent purification system or purchased from Acros Organics and used as received.

**Thin Layer Chromatography (TLC) and Column Chromatography**: Reactions were monitored by thin-layer chromatography (TLC) using silica gel $F_{254}$ pre-coated glass plates (Merck) and visualized by exposure to ultraviolet light (λ = 254 nm) or by staining with aqueous potassium permanganate ($KMnO_4$) solution (7.5 g $KMnO_4$, 50 g $K_2CO_3$, 6.25 mL aqueous 10% NaOH, 1000 mL distilled $H_2O$) or aqueous acidic ceric ammonium molybdate (IV) (CAM) solution (2.0 g $Ce(NH_4)_4(SO_4)_4{\cdot}2\ H_2O$, 48 g $(NH_4)_6Mo_7O_{24}{\cdot}4\ H_2O$, 60 mL concentrated sulfuric acid, 940 mL distilled $H_2O$) followed by heating with a heat gun (150–600 °C). Column chromatography was performed using silica gel (pore size, 60 Å; 40 to 63 µm; Merck KGaA) using a Teledyne ISCO CombiFlash EZ Prep flash purification system.

**Nuclear Magnetic Resonance (NMR) Spectroscopy**: Proton ($^1H$) and carbon ($^{13}C$) nuclear magnetic resonance spectra were recorded on a Bruker NEO 400 MHz and Bruker NEO 600 MHz. Proton chemical shifts are expressed in parts per million (ppm, δ scale) and referenced to residual undeuterated solvent signals. Carbon chemical shifts are expressed in parts per million (ppm, δ scale) and referenced to the central carbon resonance of the solvent. The reported data is represented as follows: chemical shift in parts per million (ppm, δ scale) (multiplicity, coupling constants J in Hz, integration intensity). Abbreviations used for analysis of multiplets are as follows: s (singlet), br s (broad singlet), d (doublet), t (triplet), q (quartet), p (pentet), h (hextet), and m (multiplet) or combinations thereof. NMR spectroscopy was performed on a Bruker AVIII-500 MHz spectrometer equipped with a BBO probe and using a BCUII for variable temperature. NMR spectra were acquired at 25 °C unless stated otherwise. Spectra analysis was conducted with the software MestReNova. The compounds that contain a photoswitchable moiety were thermally relaxed to the dark-adapted state by wrapping the NMR tubes with aluminum foil and placing them in a hot water bath overnight before taking an NMR.

**Mass Spectrometry (MS)**: Low-resolution mass spectrometry was performed on Waters UPLC-SQD or Agilent 1260 Infinity II/Infinity Lab LC/MSD systems. Elution was performed using a gradient from 5:95% to 100:0% MeCN: $H_2O$ with 0.1% formic acid over 5 min. High-resolution mass spectrometry was performed using Waters GCT Premier or Bruker scimaX instruments that provided mass accuracy up to > 10 million resolving power. All systems employ electrospray ionization (ESI).

### Determination of Photophysical Properties

**UV-Vis Spectroscopy**: Ultraviolet-visible (UV-vis) spectra were recorded on the Varian Cary 60 UV-Visible spectrophotometer using Brand disposable UV cuvettes (850 µL, 10 mm light path) by Brandtech Scientific Inc. Samples were stored and prepared under red light to avoid the formation of the (*Z*) isomers. Stock solutions (10 mM) were prepared in the dark and diluted with DMSO and PBS to a final concentration of 50 µM for measurement.

**Wavelength Scan**: Light at different wavelengths was provided by an Optoscan Monochromator with an Optosource (75 mW lamp), which was controlled through a program written in Matlab. UV-vis spectra of **Azo-morphines** were recorded following irradiation with different wavelengths for 5 min using a monochromator. Measurement was started from the dark-adapted state followed by 550 nm irradiation and incrementally decreased the wavelength. The wavelength scan of **Azo-morphines** were measured in phosphate-buffered saline (PBS) with 10% DMSO.

**Thermal Relaxation, Photocycling and PSS**: Thermal relaxation was measured by preirradiating **Azo-morphines** with 365-nm light and observing the absorption at 320 nm over 36-72 hours at 37°C in PBS with 10% DMSO in tightly sealed cuvettes. The reversible switching and photochemical stability of **Azo-morphines** were demonstrated in PBS with 10% DMSO, cycling the irradiation of the monochromator between 365 nm and 460 nm. PSSs (photostationary state) were obtained from the internal UV-vis detector of the liquid chromatography–mass spectrometry (LC-MS) by irradiating the sample in 100 µM $H_2O$ with 10% MeCN with different wavelengths for 10 mins before injection. Samples were eluted using a 5 min gradient from 95:5 to 0:100 (water:acetonitrile w/ 0.1% v/v formic acid), and 1 min hold. The absorption peaks at isosbestic point (280 nm) for the *trans* and *cis* isomers were integrated to determine their ratios on LCMS. We infer any discrepancies between wavelength scan and PSS to be from solvatochromism.

### Synthetic Procedures and Characterization

**(4*R*,4a*S*,7*R*,7a*R*,12b*S*)-7-(benzyl(methyl)amino)-3-(cyclopropylmethyl)-1,2,3,4,5,6,7,7a-octahydro-4aH-4,12-methanobenzofuro[3,2-e]isoquinoline-4a,9-diol (S-1)**

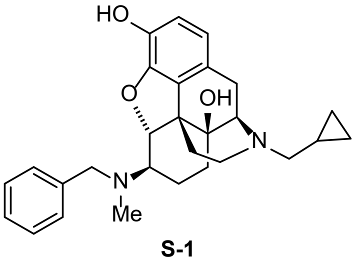


**S-1**

**Procedure:** In a flame-dried 10 mL round bottom flask, naltrexone hydrochloride (113.4 mg, 0.3 mmol, 1.0 eq.), *N*-methylbenzylamine (155 µL, 1.2 mmol, 4.0 eq.), benzoic acid (55 mg, 0.45 mmol, 1.5 eq.) and *p*-TsOH·$H_2O$ (11.4 mg, 0.06 mmol, 0.2 eq.) were dissolved in anhydrous benzene (6 mL, 0.05 M). The solution was refluxed in the presence of a Dean-Stark apparatus for 48 h at 140 °C. Upon completion of the reaction, the reaction mixture was concentrated *in vacuo*. The residue was dissolved in anhydrous MeOH (3 mL) and cooled to −20 °C. A freshly prepared solution of $NaBH_3CN$ (56.6 mg, 0.9 mmol, 3.0 eq.) in anhydrous MeOH (3 mL) was then added. The reaction mixture was allowed to slowly warm up to rt for 4 h and stir at rt for another 14 h. Upon completion of the reaction, the reaction was diluted with EtOAc (30 mL), transferred to a separatory funnel and the organic phase was washed with sat. $NaHCO_{3(aq)}$ (2x 20 mL), dried over $Na_2SO_4$, filtered and concentrated *in vacuo*. The crude product was purified on silica gel (gradient from 0% to 5% MeOH in $CH_2Cl_2$ with 1% ammonia, 7 M in methanol) to afford the desired tertiary amine (102 mg, 0.23 mmol, 76% yield) as a white foamy solid.

**Characterization**:

**$^1$H NMR** (400 MHz, $CDCl_3$) δ 7.38 – 7.26 (m, 4H), 7.25 – 7.17 (m, 1H), 6.64 (d, *J* = 8.1 Hz, 1H), 6.51 (d, *J* = 8.1 Hz, 1H), 4.69 (d, *J* = 8.0 Hz, 1H), 3.80 (d, *J* = 13.5 Hz, 1H), 3.59 (d, *J* = 13.6 Hz, 1H), 3.15 – 2.87 (m, 2H), 2.70 – 2.49 (m, 3H), 2.42 – 2.31 (m, 5H), 2.23 (td, *J* = 12.3, 4.9 Hz, 1H), 2.10 (td, *J* = 11.9, 3.5 Hz, 1H), 2.03 – 1.88 (m, 1H), 1.61 (dd, *J* = 10.4, 3.6 Hz, 2H), 1.50 – 1.41 (m, 1H), 1.36 – 1.23 (m, 2H), 0.90 – 0.76 (m, 1H), 0.58 – 0.46 (m, 2H), 0.14 – 0.06 (m, 2H).
**$^{13}$C NMR** (101 MHz, $CDCl_3$) δ 142.4, 139.8, 139.6, 131.9, 128.8, 128.3, 127.0, 124.8, 118.7, 116.6, 91.4, 70.5, 63.5, 62.6, 59.3, 58.9, 48.0, 44.2, 38.2, 30.9, 30.7, 22.8, 17.9, 9.6, 4.0, 4.0.
The characterizations match the data that has been reported previously.[1]

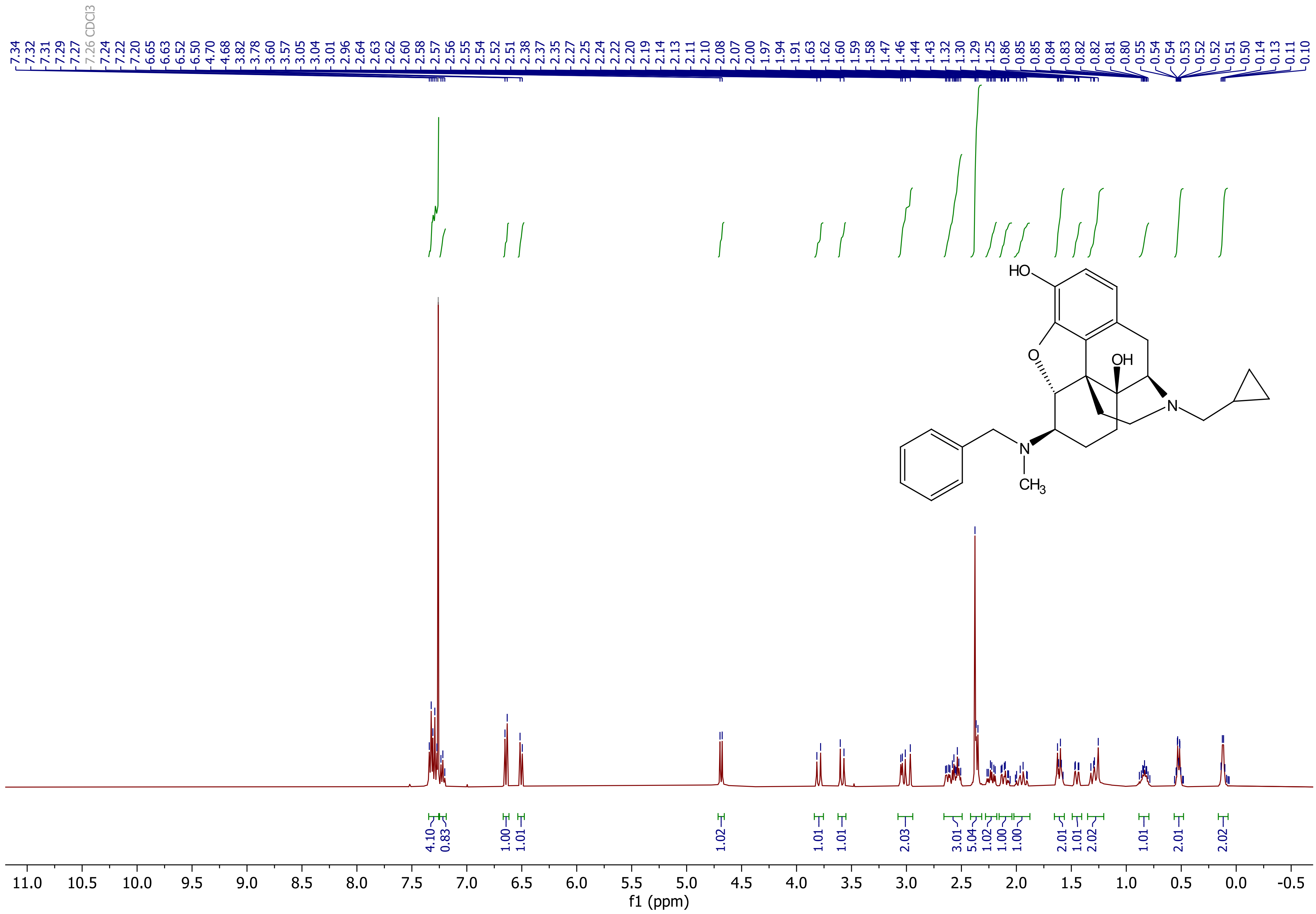

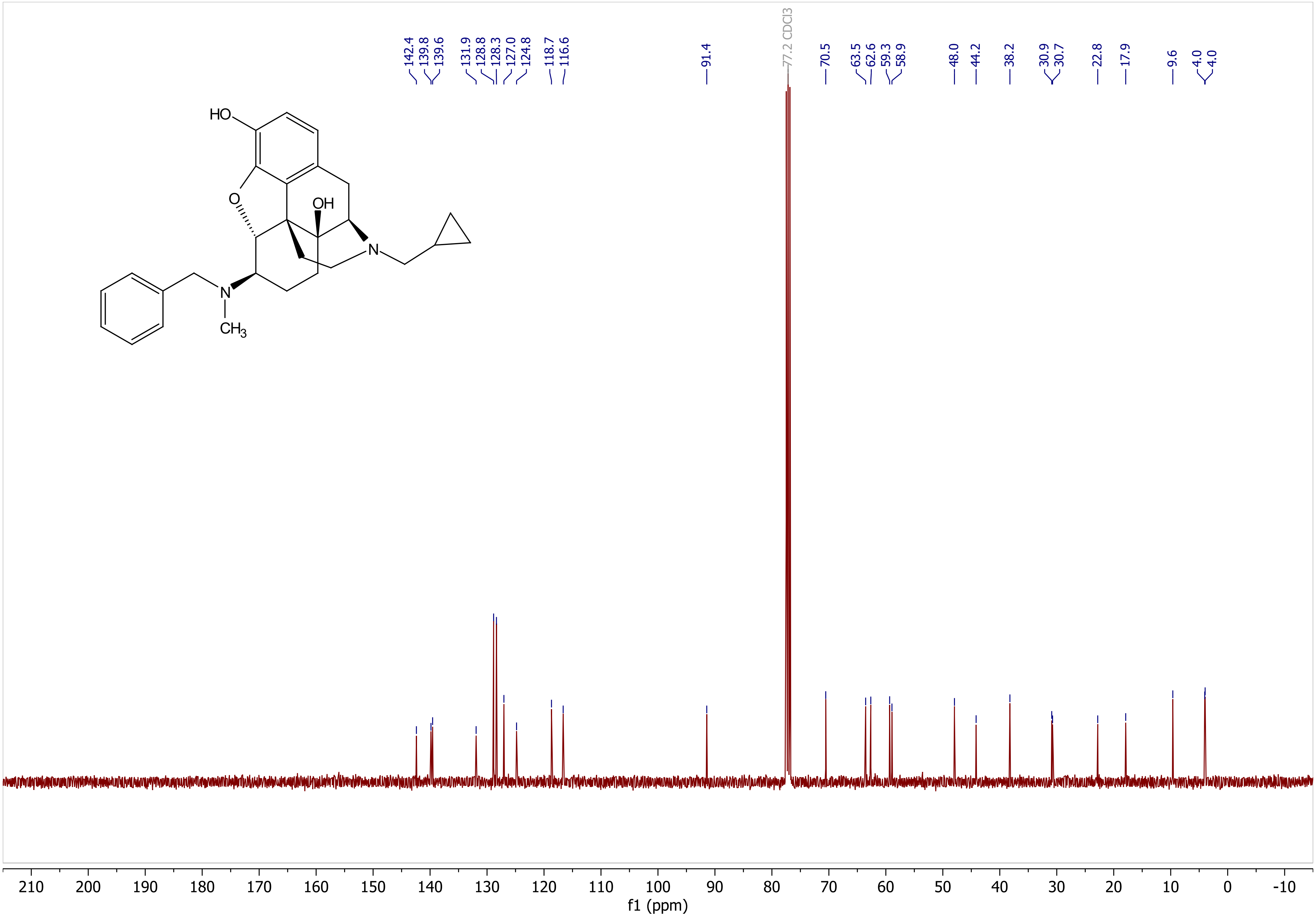

142.4
139.8
139.6
131.9
128.8
128.3
127.0
124.8
118.7
116.6
91.4
77.2 CDCl3
70.5
63.5
62.6
59.3
58.9
48.0
44.2
38.2
30.9
30.7
22.8
17.9
9.6
4.0
4.0
HO
O
OH
N
N
CH3
210 200 190 180 170 160 150 140 130 120 110 100 90 80 70 60 50 40 30 20 10 0 -10
f1 (ppm)

**(4*R*,4a*S*,7*R*,7a*R*,12b*S*)-3-(cyclopropylmethyl)-7-(methylamino)-1,2,3,4,5,6,7,7a-octahydro-4aH-4,12-methanobenzofuro[3,2-e]isoquinoline-4a,9-diol (1)**

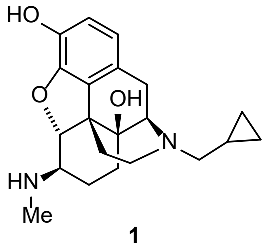


**1**

**Procedure**: To a flame-dried 5 mL round bottom flask, **S-1** (78 mg, 0.175 mmol, 1.0 eq.) and Pd/C 10% (37 mg, 0.035 mmol, 0.2 eq.) was added. The atmosphere was exchanged to argon, and a 1:1 ratio of *i*-PrOH:AcOH was added (1.75 mL respectively, 3.5 mL total, 0.05 M). The solution was then purged three times with $H_2$ (1 atm), and two hydrogen balloons were used for the reaction. The reaction mixture was stirred at room temperature until complete conversion and the progress of the reaction was monitored with LCMS. Upon the completion of the reaction, the atmosphere was changed back to argon. The reaction mixture was vacuum-filtered over celite and rinsed with *i*-PrOH and $CH_2Cl_2$. The filtrate was concentrated *in vacuo*. The crude product was purified on silica gel (gradient from 5% to 15% MeOH in $CH_2Cl_2$ with 1% ammonia, 7 M in methanol) to afford the desired secondary amine (60 mg, 0.17 mmol, 97% yield) as a pale yellow solid.

**Characterization**:

**$^1$H NMR** (400 MHz, $CDCl_3$) δ 6.65 (d, *J* = 8.1 Hz, 1H), 6.54 (d, *J* = 8.1 Hz, 1H), 4.51 (d, *J* = 7.5 Hz, 1H), 3.10 – 2.88 (m, 2H), 2.59 (ddt, *J* = 18.5, 14.1, 5.1 Hz, 3H), 2.47 (s, 3H), 2.35 (d, *J* = 6.5 Hz, 2H), 2.27 – 2.07 (m, 2H), 1.97 – 1.82 (m, 1H), 1.71 – 1.57 (m, 2H), 1.45 – 1.33 (m, 2H), 0.89 – 0.76 (m, 1H), 0.51 (dt, *J* = 8.2, 3.1 Hz, 2H), 0.17 – 0.06 (m, 2H).
**$^{13}$C NMR** (101 MHz, $CDCl_3$) δ 142.2, 140.9, 131.5, 123.5, 119.4, 118.3, 91.3, 70.5, 62.6, 59.3, 59.3, 47.5, 44.2, 30.9, 30.5, 30.5, 22.8, 21.7, 9.6, 4.0, 3.9.
The characterizations match the data that has been reported previously. [1]

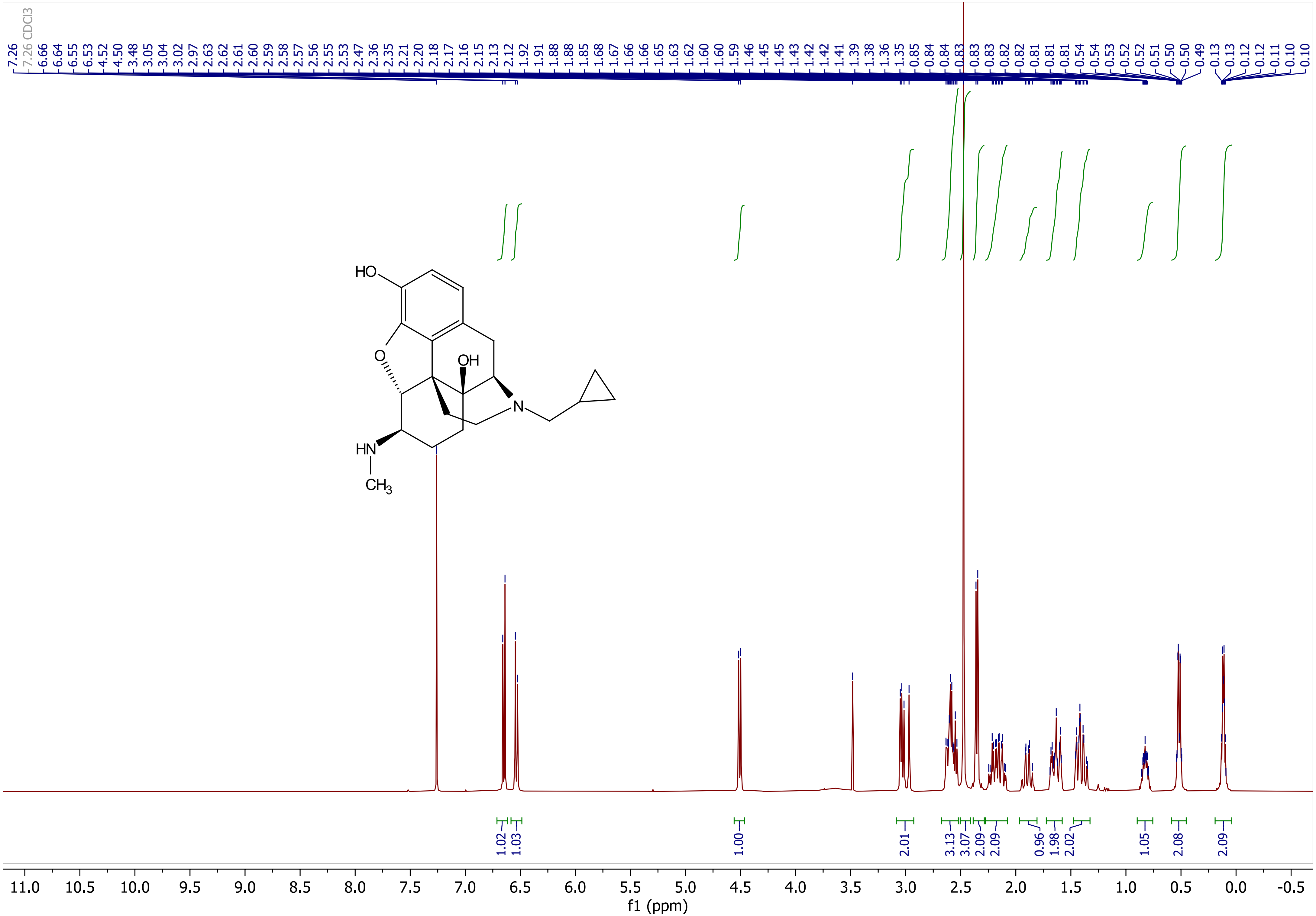
7.26
7.26 CDCl3
6.66
6.64
6.55
6.53
4.52
4.50
3.48
3.05
3.04
3.02
2.97
2.63
2.62
2.61
2.60
2.59
2.58
2.57
2.56
2.55
2.53
2.47
2.36
2.35
2.21
2.20
2.18
2.17
2.16
2.15
2.13
2.12
1.92
1.91
1.88
1.88
1.85
1.68
1.67
1.66
1.66
1.65
1.63
1.62
1.60
1.60
1.59
1.46
1.45
1.45
1.43
1.42
1.42
1.41
1.39
1.38
1.36
1.35
0.85
0.84
0.84
0.83
0.83
0.83
0.82
0.82
0.81
0.81
0.81
0.54
0.54
0.53
0.52
0.52
0.51
0.50
0.49
0.13
0.13
0.12
0.12
0.11
0.10
0.10
HO
OH
O
N
HN
CH3
1.02
1.03
1.00
2.01
3.13
3.07
2.09
2.09
0.96
1.98
2.02
1.05
2.08
2.09
11.0 10.5 10.0 9.5 9.0 8.5 8.0 7.5 7.0 6.5 6.0 5.5 5.0 4.5 4.0 3.5 3.0 2.5 2.0 1.5 1.0 0.5 0.0 -0.5
f1 (ppm)

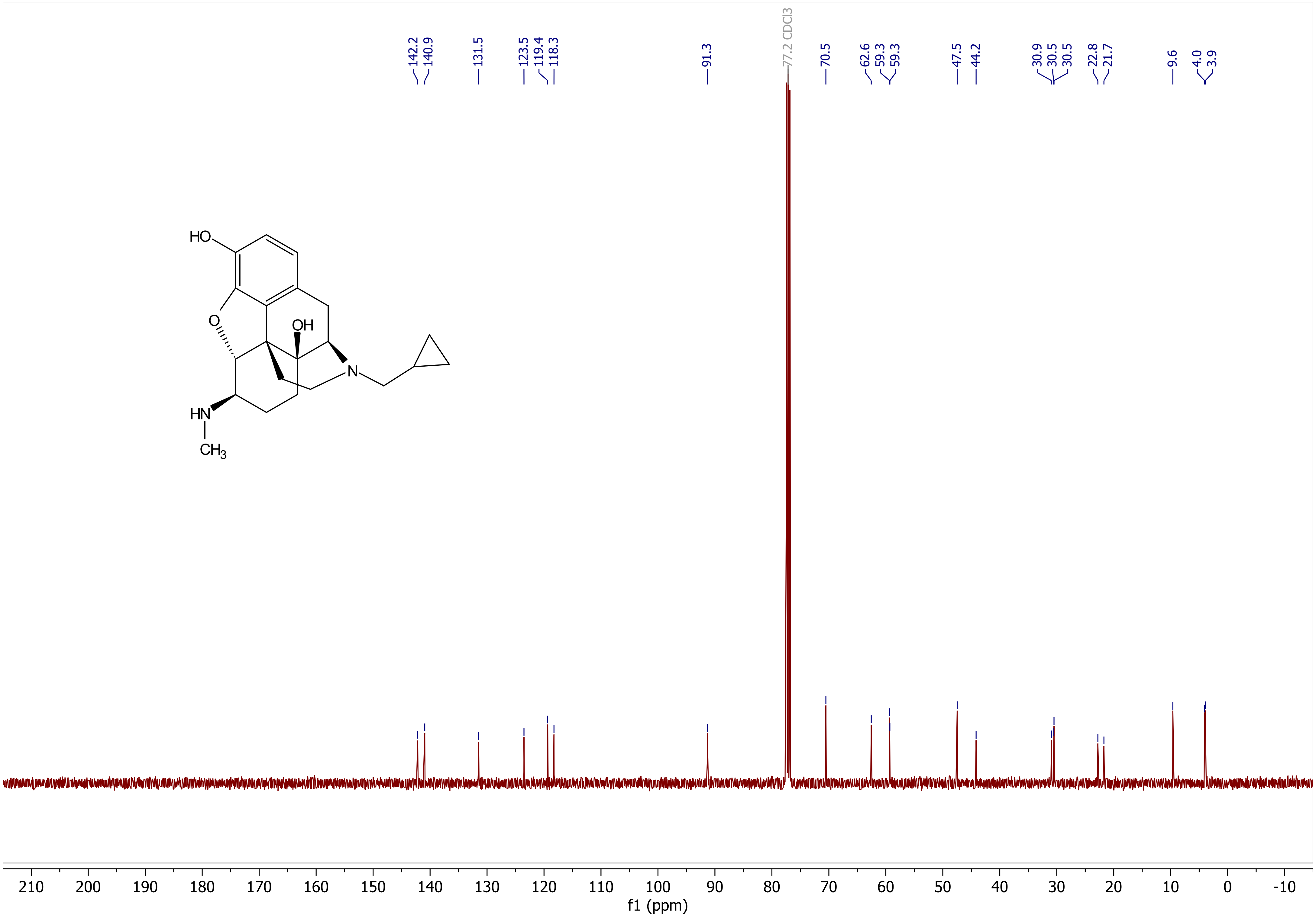

142.2
140.9
131.5
123.5
119.4
118.3
91.3
77.2 CDCl3
70.5
62.6
59.3
59.3
47.5
44.2
30.9
30.5
30.5
22.8
21.7
9.6
4.0
3.9
HO
O
OH
N
HN
CH3
210 200 190 180 170 160 150 140 130 120 110 100 90 80 70 60 50 40 30 20 10 0 -10
f1 (ppm)

**3-(phenyldiazenyl)benzoic acid (S-2)**

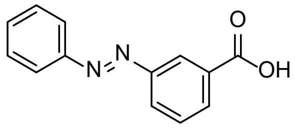

**S-2**

**Procedure:** In a 50 mL round bottom flask, 3-aminobenzoic acid (411 mg, 3.0 mmol, 1.0 eq.) and nitrosobenzene (418 mg, 3.9 mmol, 1.3 eq.) were suspended in AcOH (12 mL, 0.25 M). The reaction mixture was stirred vigorously (800 rpm) for 96 h at room temperature. After the completion of the reaction, celite was added to the flask and the solution was concentrated *in vacuo*. The crude mixture was then loaded on silica gel and purified by flash chromatography (0-5% MeOH in EtOAc) to afford the desired azobenzene (594 mg, 2.63 mmol, 88% yield) as an orange solid.

**Characterization**:

**$^1$H NMR** (600 MHz, $CDCl_3$) δ 8.67 (d, *J* = 1.9 Hz, 1H), 8.24 (dt, *J* = 7.7, 1.4 Hz, 1H), 8.17 (dt, *J* = 7.9, 1.5 Hz, 1H), 7.99 – 7.94 (m, 2H), 7.65 (t, *J* = 7.8 Hz, 1H), 7.58 – 7.48 (m, 3H).
**$^{13}$C NMR** (151 MHz, $CDCl_3$) δ 171.2, 152.8, 152.6, 132.3, 131.7, 130.5, 129.5, 129.3, 127.8, 124.9, 123.2.
The characterizations match the data that has been reported previously.[2]

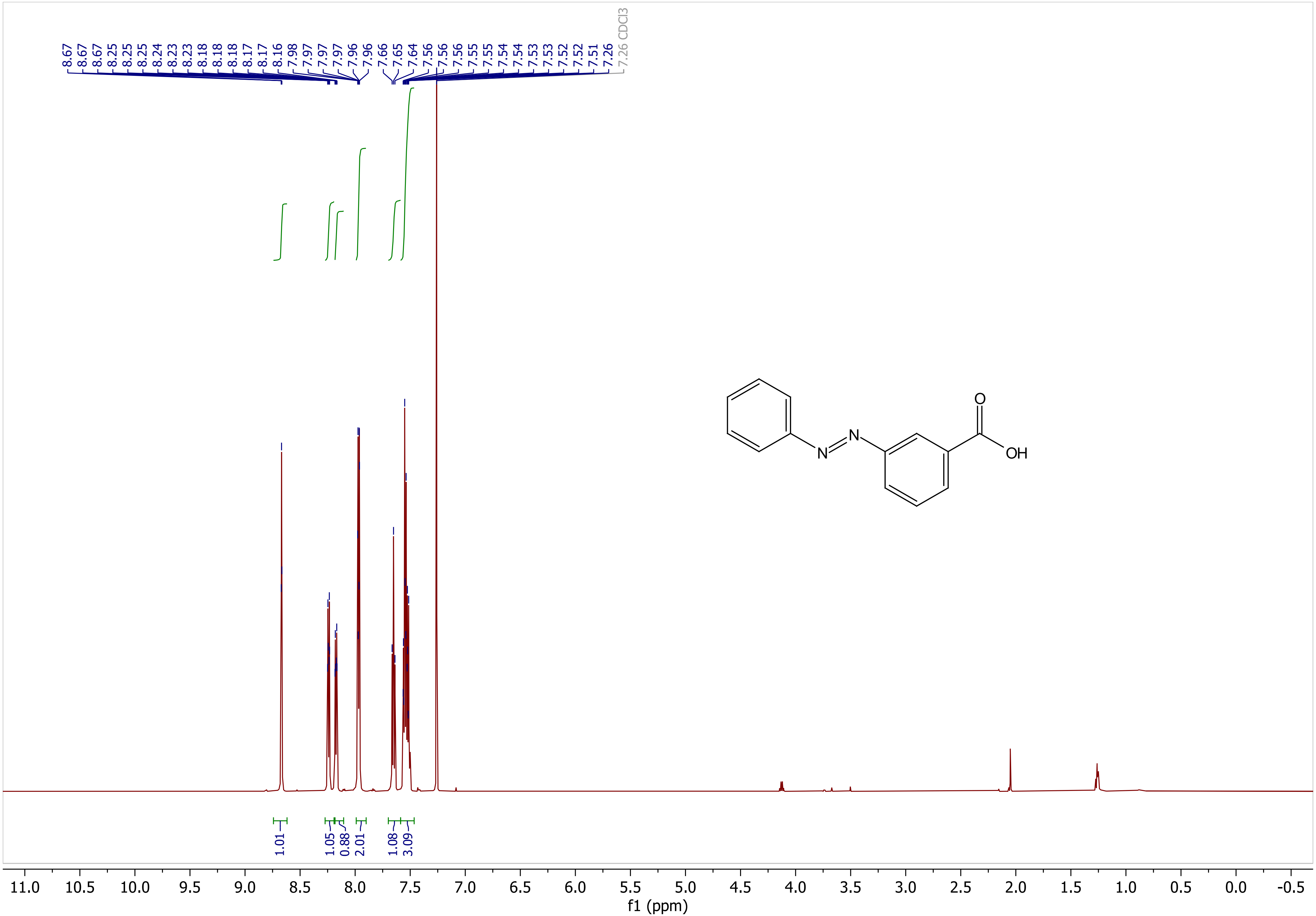

8.67
8.67
8.67
8.25
8.25
8.24
8.23
8.23
8.18
8.18
8.18
8.17
8.17
8.16
7.98
7.97
7.97
7.97
7.96
7.96
7.66
7.65
7.64
7.56
7.56
7.56
7.55
7.55
7.54
7.54
7.53
7.53
7.52
7.52
7.51
7.26
7.26 CDCl3
N
N
O
OH
1.01
1.05
0.88
2.01
1.08
3.09
11.0
10.5
10.0
9.5
9.0
8.5
8.0
7.5
7.0
6.5
6.0
5.5
5.0
4.5
4.0
3.5
3.0
2.5
2.0
1.5
1.0
0.5
0.0
-0.5
f1 (ppm)

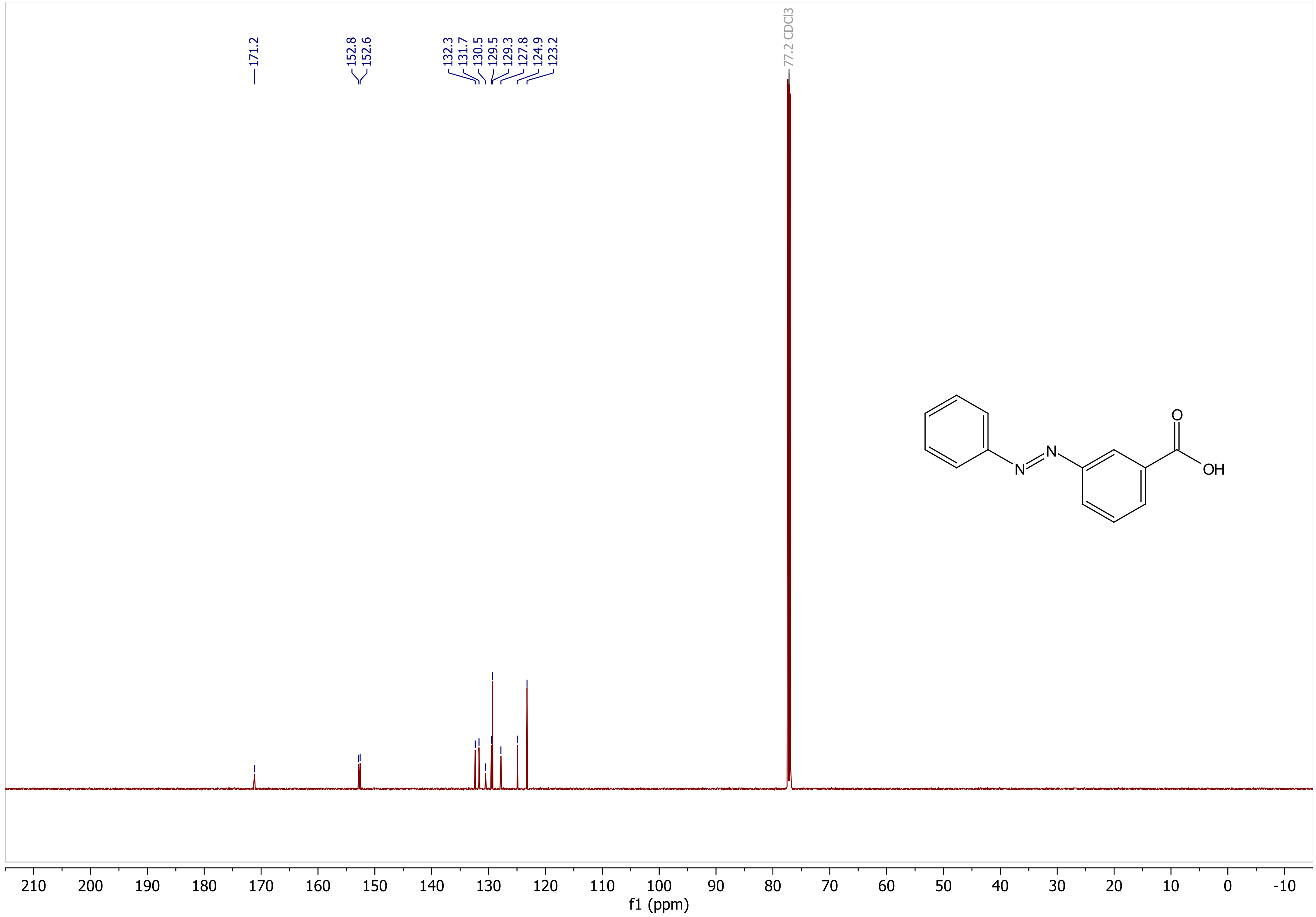

171.2
152.8
152.6
132.3
131.7
130.5
129.5
129.3
127.8
124.9
123.2
77.2 CDCl3
N
N
O
OH
210
200
190
180
170
160
150
140
130
120
110
100
90
80
70
60
50
40
30
20
10
0
-10
f1 (ppm)

**2-(4-(phenyldiazenyl)phenyl)acetic acid (2)**

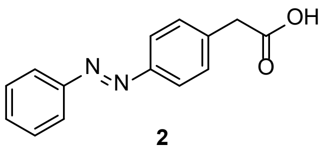

**2**

**Procedure:** In a 50 mL round bottom flask, 4-aminophenylacetic acid (454 mg, 3.0 mmol, 1.0 eq.) and nitrosobenzene (418 mg, 3.9 mmol, 1.3 eq.) were suspended in AcOH (12 mL, 0.25 M). The reaction mixture was stirred vigorously (800 rpm) for 96 h at room temperature. After the completion of the reaction, celite was added to the flask and the solution was concentrated *in vacuo*. The crude mixture was then loaded on silica gel and purified by flash chromatography (0-5% MeOH in EtOAc) to afford the desired azobenzene (334 mg, 1.39 mmol, 46% yield) as an orange solid.

**Characterization**:

**$^1$H NMR** (600 MHz, $CDCl_3$) δ 7.91 (td, $J$ = 5.9, 3.1 Hz, 4H), 7.55 – 7.43 (m, 5H), 3.76 (s, 2H).
**$^{13}$C NMR** (151 MHz, $CDCl_3$) δ 174.8, 152.8, 152.0, 136.4, 131.2, 130.3, 129.2, 123.3, 123.0, 40.6.
The characterizations match the data that has been reported previously.[3]

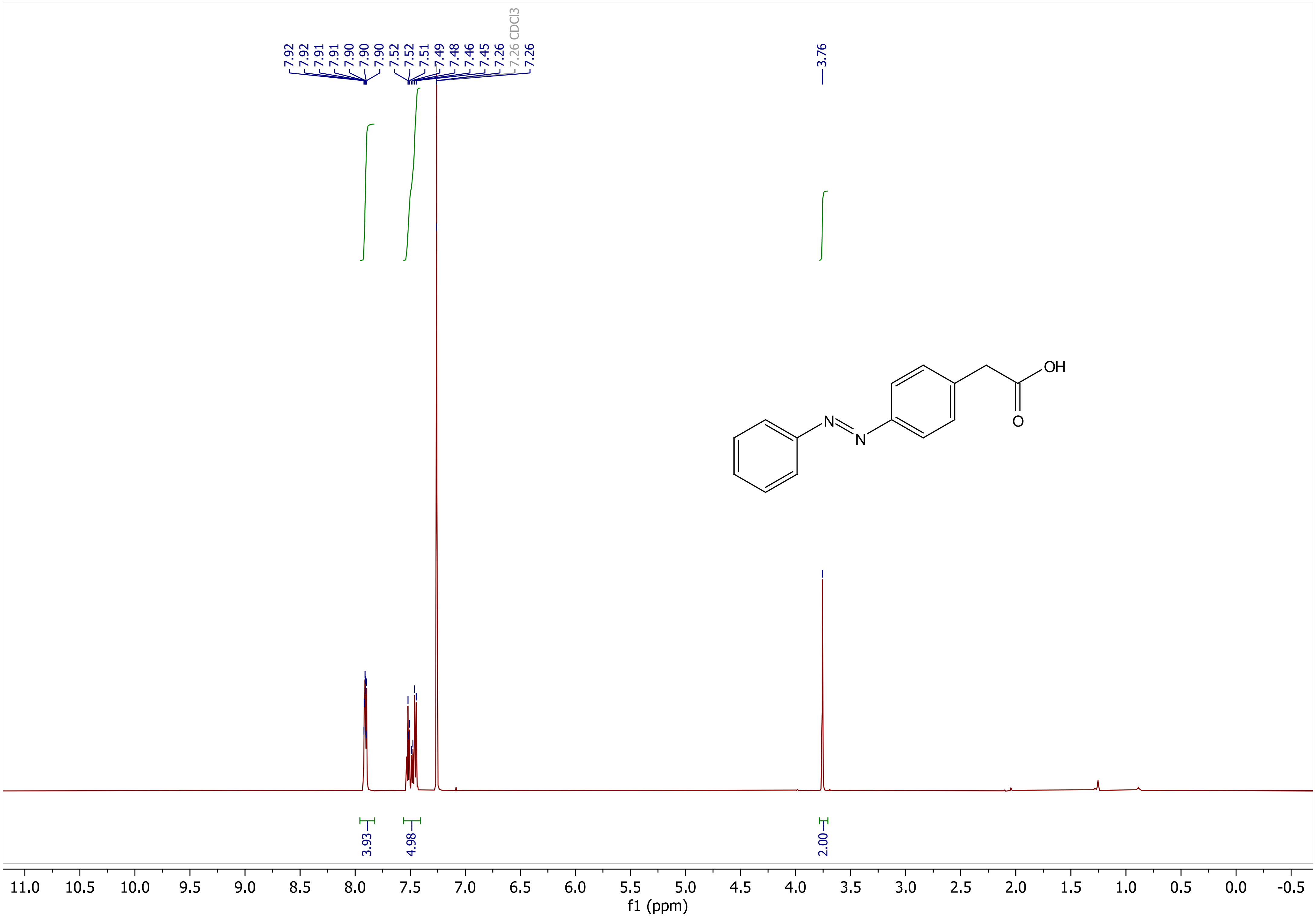

7.92
7.92
7.91
7.91
7.90
7.90
7.90
7.52
7.52
7.51
7.49
7.48
7.46
7.45
7.26
7.26 CDCl3
7.26
3.76
OH
O
N
N
3.93
4.98
2.00
11.0 10.5 10.0 9.5 9.0 8.5 8.0 7.5 7.0 6.5 6.0 5.5 5.0 4.5 4.0 3.5 3.0 2.5 2.0 1.5 1.0 0.5 0.0 -0.5
f1 (ppm)

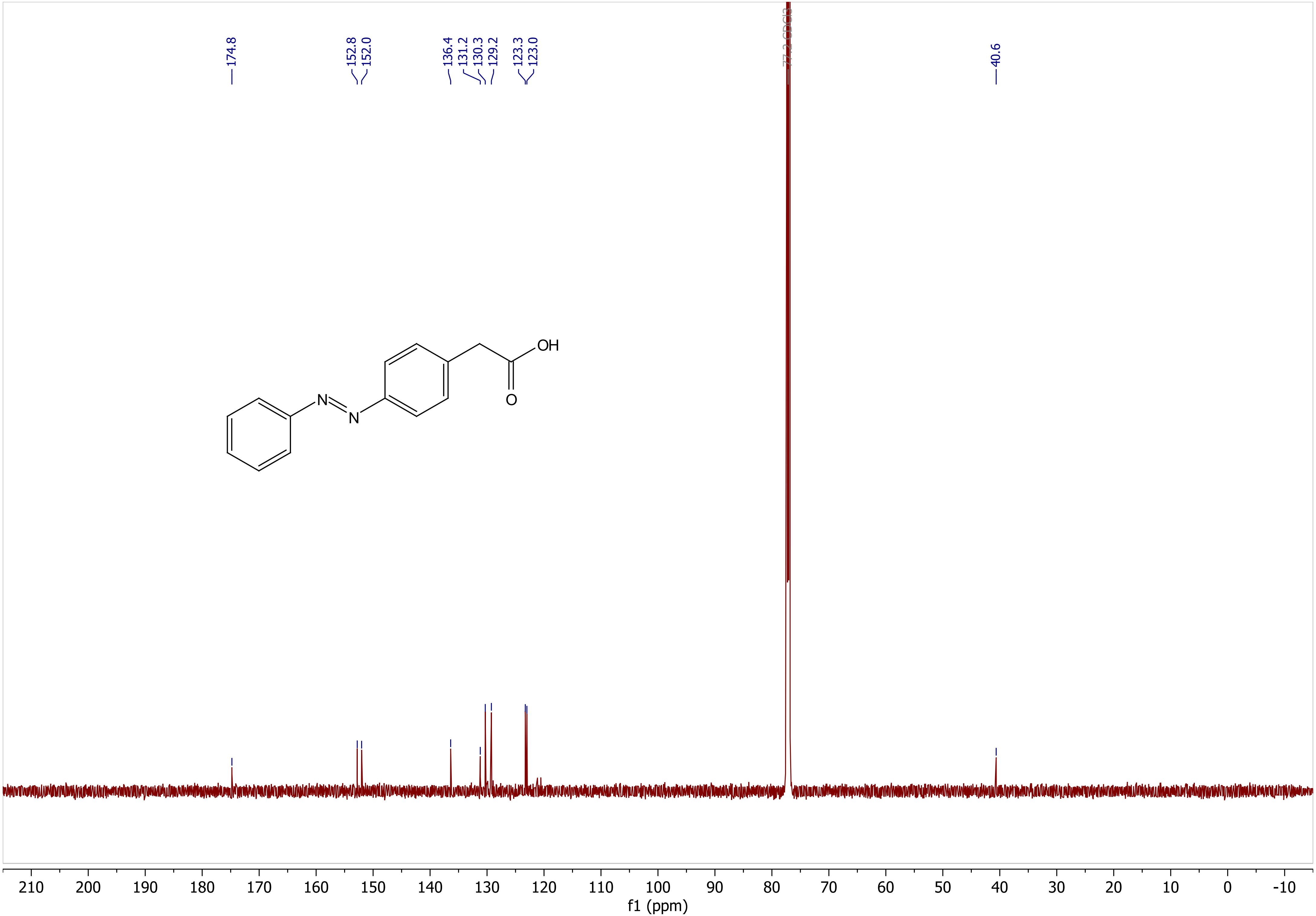
174.8
152.8
152.0
136.4
131.2
130.3
129.2
123.3
123.0
77.2 CDCl3
40.6
OH
O
N
N
210
200
190
180
170
160
150
140
130
120
110
100
90
80
70
60
50
40
30
20
10
0
-10
f1 (ppm)

**2-(4-(phenyldiazenyl)phenyl)acetic acid (S-3)**

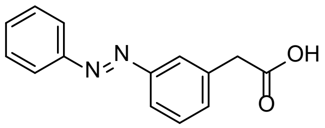

**S-3**

**Procedure:** In a 50 mL round bottom flask, 3-aminophenylacetic acid (454 mg, 3.0 mmol, 1.0 eq.) and nitrosobenzene (418 mg, 3.9 mmol, 1.3 eq.) were suspended in AcOH (12 mL, 0.25 M). The reaction mixture was stirred vigorously (800 rpm) for 96 h at room temperature. After the completion of the reaction, celite was added to the flask and the solution was concentrated *in vacuo*. The crude mixture was then loaded on silica gel and purified by flash chromatography (0-5% MeOH in EtOAc) to afford the desired azobenzene (625 mg, 2.60 mmol, 87% yield) as an orange solid.

**Characterization**:

**$^1$H NMR** (600 MHz, $CDCl_3$) δ 7.94 – 7.89 (m, 2H), 7.88 – 7.84 (m, 2H), 7.55 – 7.45 (m, 4H), 7.41 (dt, *J* = 7.6, 1.5 Hz, 1H), 3.78 (s, 2H).
**$^{13}$C NMR** (151 MHz, $CDCl_3$) δ 176.3, 153.0, 152.7, 134.5, 132.0, 131.3, 129.5, 129.2, 123.6, 123.0, 122.5, 40.8.
**HRMS (ESI)**: m/z $[M-H]^-$ calcd for $C_{14}H_{11}N_2O_2$: 239.082601, found: 239.082519 (Δ = 0.3 ppm)

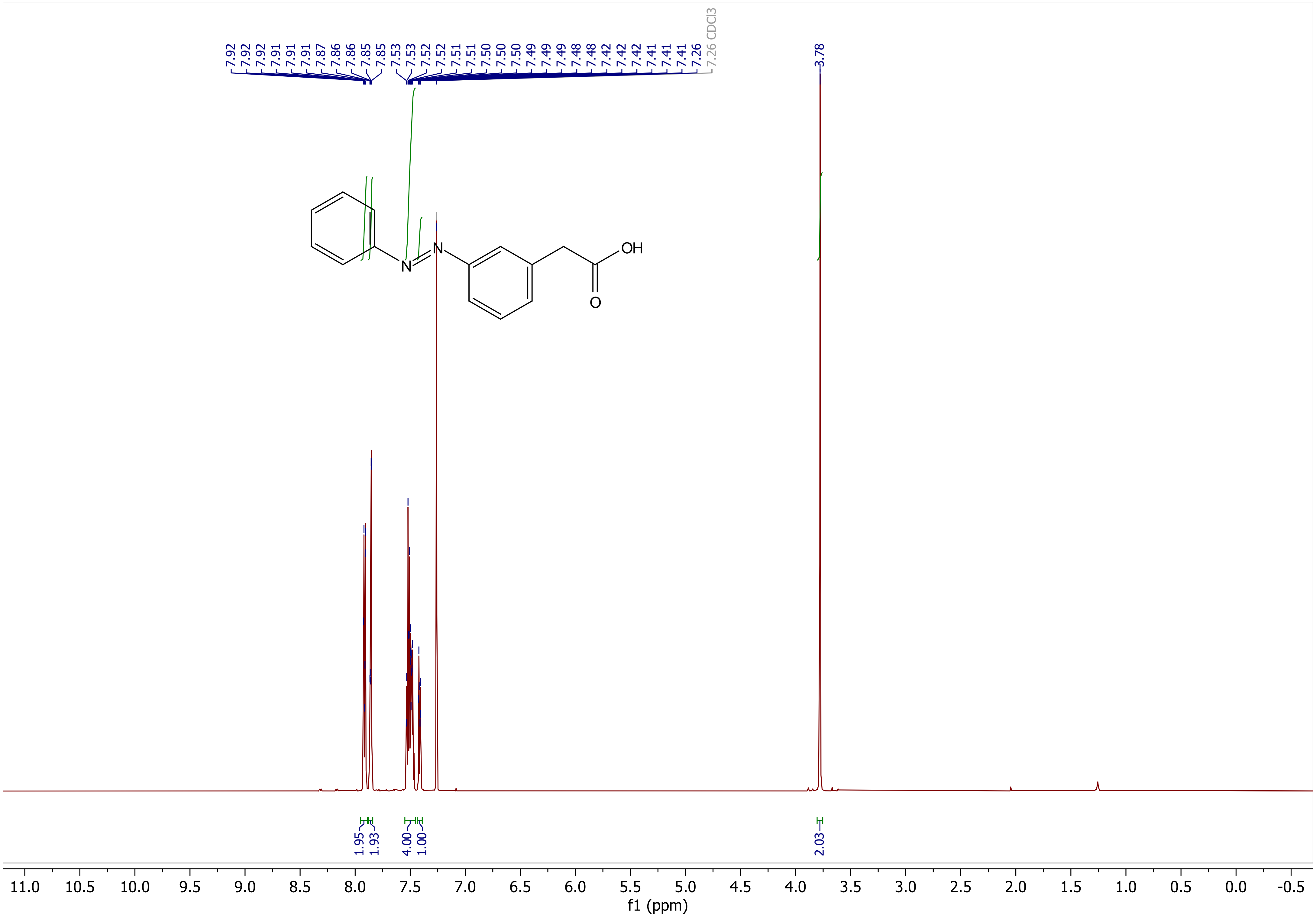

7.92
7.92
7.92
7.91
7.91
7.87
7.86
7.86
7.85
7.85
7.53
7.53
7.52
7.52
7.51
7.51
7.50
7.50
7.50
7.49
7.49
7.49
7.48
7.48
7.42
7.42
7.42
7.41
7.41
7.41
7.26
7.26 CDCl3
3.78
N
N
OH
O
1.95
1.93
4.00
1.00
2.03
11.0 10.5 10.0 9.5 9.0 8.5 8.0 7.5 7.0 6.5 6.0 5.5 5.0 4.5 4.0 3.5 3.0 2.5 2.0 1.5 1.0 0.5 0.0 -0.5
f1 (ppm)

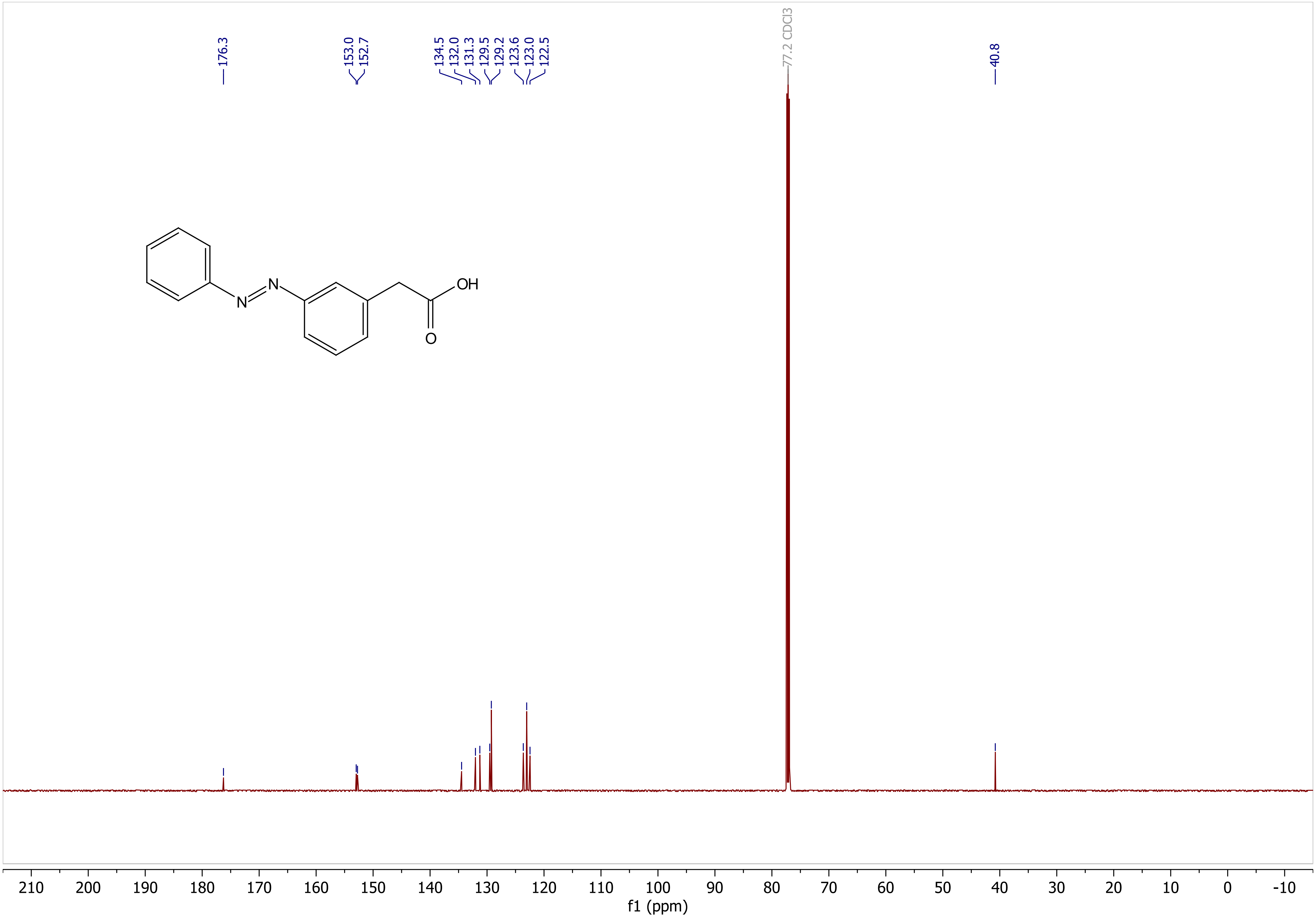
176.3
153.0
152.7
134.5
132.0
131.3
129.5
129.2
123.6
123.0
122.5
77.2 CDCl3
40.8
OH
O
N
N
210
200
190
180
170
160
150
140
130
120
110
100
90
80
70
60
50
40
30
20
10
0
-10
f1 (ppm)

**<u>*N*-((4*R*,4a*S*,7*R*,7a*R*,12b*S*)-3-(cyclopropylmethyl)-4a,9-dihydroxy-2,3,4,4a,5,6,7,7a-octahydro-1H-4,12-methanobenzofuro[3,2-*e*]isoquinolin-7-yl)-*N*-methyl-4-((*E*)-phenyldiazenyl)benzamide (AM-1)</u>**

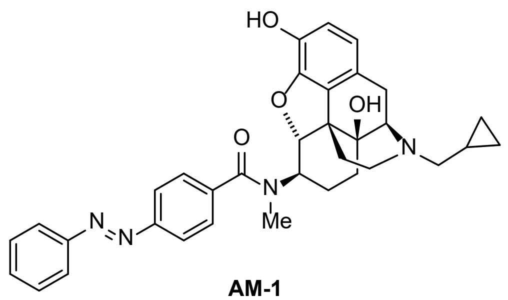

**AM-1**

**<u>Procedure:</u>** In a flame-dried 5 mL round bottom flask, naltrexone-derived amine (**1**) (14 mg, 0.040 mmol, 1.0 eq.), 4-(phenylazo)benzoic acid (11 mg, 0.048 mmol, 1.2 eq.) and COMU coupling reagent (19 mg, 0.044 mmol, 1.1 eq.) were dissolved in anhydrous DMF (0.40 mL, 0.1 M). The reaction was stirred under argon at 0 °C, and anhydrous DIPEA (15 µL, 0.088 mmol, 2.2 eq.) was added. The reaction mixture was stirred at 0 °C for 1 h before the cooling bath was removed, and the solution was allowed to warm up and stirred at room temperature for another 16 h. Upon the completion of the reaction, the mixture was diluted with EtOAc:$Et_2O$ 1:1 (20 mL) and transferred to a separatory funnel. 0.1 M $NaHCO_{3(aq)}$ (10 mL) was added to the separatory funnel, and layers were separated. The aqueous layer was extracted with EtOAc (2x20 mL). The combined organic layer was extracted with 10% $LiCl_{(aq)}$ (2x20 mL) and brine (1x20 mL). The organic layer was dried over $Na_2SO_4$, filtered, and concentrated *in vacuo*. The crude product was purified on silica gel (gradient from 0% to 10% MeOH in $CH_2Cl_2$ with 1% ammonia, 7 M in methanol) to afford the desired **AM-1** (24.6 mg, 0.033 mmol, 82% yield) as an orange oil. For long-term storage, the compound was kept as a trifluoroacetic acid or a formic acid salt. Trifluoroacetic acid or formic acid (1.3 eq) in acetonitrile (99:1 MeCN:TFA/formic acid) was added to the compound and concentrated *in vacuo* to afford the salt in order to avoid the formation of *N*-oxides. The dilution of the acid was necessary to avoid the decomposition of the product.

**<u>Characterization</u>**:

**$^1$H NMR** (500 MHz, $CD_3OD$, exists as ~8:2 rotamers, only peaks of the major rotamer shown) δ 7.92 – 7.88 (m, 2H), 7.83 (d, *J* = 8.1 Hz, 2H), 7.63 – 7.50 (m, 5H), 6.66 (d, *J* = 8.2 Hz, 1H), 6.56 (d, *J* = 8.2 Hz, 1H), 4.92 (d, *J* = 8.1 Hz, 1H), 3.81 (d, *J* = 5.7 Hz, 1H), 3.53 (ddd, *J* = 12.8, 8.1, 4.4 Hz, 1H), 3.24 (q, *J* = 6.8 Hz, 1H), 3.17 (s, 3H), 3.11 – 3.03 (m, 2H), 2.98 (dd, *J* = 19.6, 6.0 Hz, 1H), 2.88 – 2.80 (m, 1H), 2.66 – 2.53 (m, 2H), 2.35 – 2.22 (m, 1H), 1.74 – 1.66 (m, 1H), 1.54 – 1.45 (m, 1H), 1.34 – 1.20 (m, 2H), 1.07 – 1.00 (m, 1H), 0.80 – 0.67 (m, 2H), 0.51 – 0.41 (m, 2H).
**$^{13}$C NMR** (151 MHz, $CD_3OD$, exists as ~8:2 rotamers, only resonance of major rotamer shown) δ 174.2, 154.0, 153.9, 142.7, 142.5, 139.7, 132.7, 130.3, 129.1, 128.8, 124.0, 123.9, 123.9, 120.8, 118.9, 88.7, 71.3, 67.6, 66.9, 63.8, 61.8, 59.3, 38.6, 31.4, 29.0, 23.8, 23.5, 15.4, 5.4, 3.7.
* note: the NMR of this compound at room temperature is a rotameric mixture.

**HRMS (ESI)**: m/z $[M+H]^+$ calcd for $C_{34}H_{36}N_4O_4$: 565.280932, found: 565.283392 (Δ = 4.4 ppm)
**$R_f$** (4% MeOH in $CH_2Cl_2$ with 1% ammonia, 7 M in methanol): 0.47; stains yellow with $KMnO_4$; detected by UV

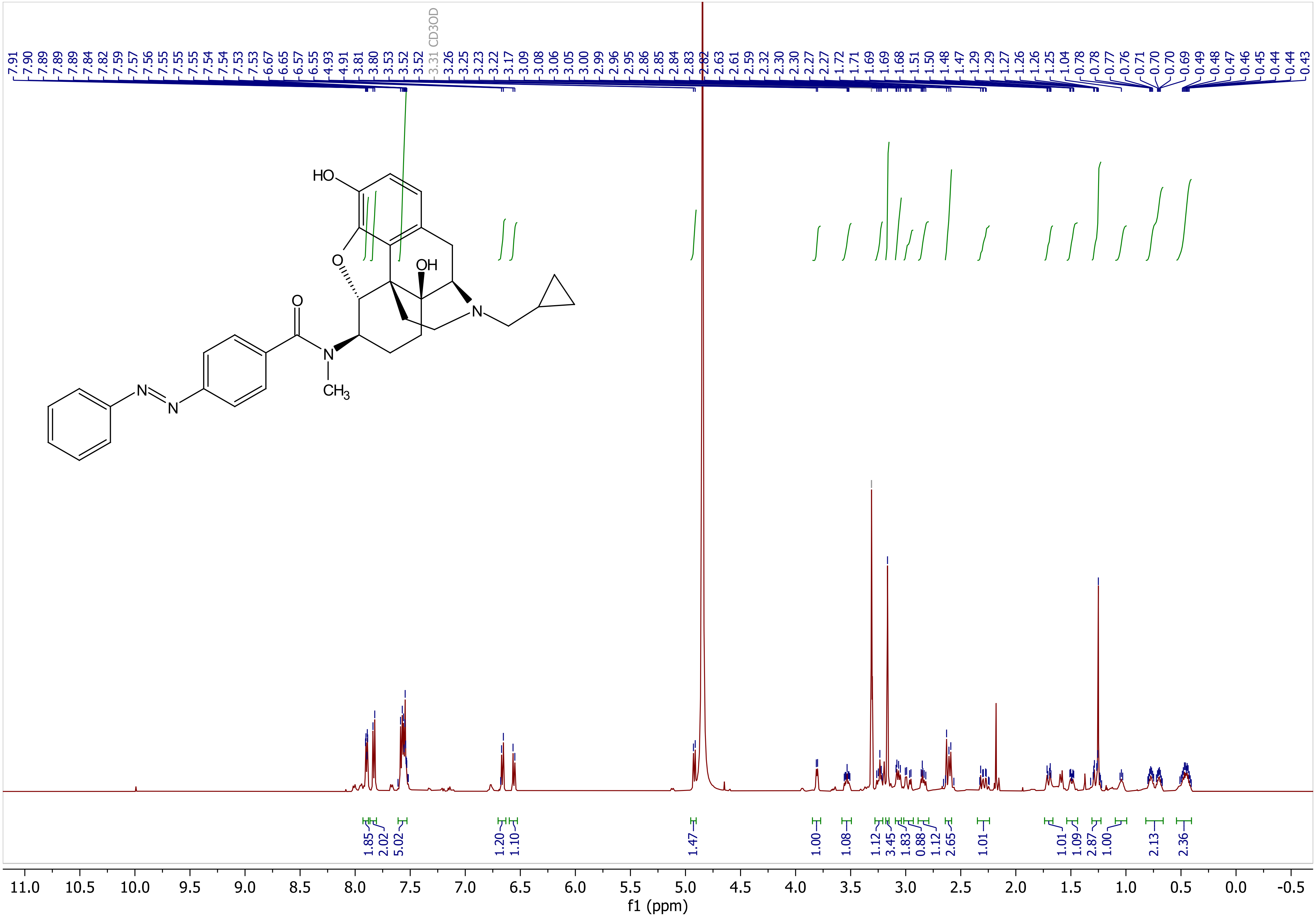

7.91 7.90 7.89 7.89 7.84 7.82 7.59 7.57 7.56 7.55 7.55 7.55 7.54 7.54 7.53 7.53 6.67 6.65 6.57 6.55 4.93 4.91 3.81 3.80 3.53 3.52 3.52 3.31 CD3OD 3.26 3.25 3.23 3.22 3.17 3.09 3.08 3.06 3.05 3.00 2.99 2.96 2.95 2.86 2.85 2.84 2.83 2.82 2.63 2.61 2.59 2.32 2.30 2.30 2.27 2.27 1.72 1.71 1.69 1.69 1.68 1.51 1.50 1.48 1.47 1.29 1.29 1.27 1.26 1.26 1.25 1.04 0.78 0.78 0.77 0.76 0.71 0.70 0.70 0.69 0.49 0.48 0.47 0.46 0.45 0.44 0.44 0.43
HO
O
OH
N
O
N
CH3
N
N
1.85 2.02 5.02 1.20 1.10 1.47 1.00 1.08 1.12 3.45 1.83 0.88 1.12 2.65 1.01 1.01 1.09 2.87 1.00 2.13 2.36
11.0 10.5 10.0 9.5 9.0 8.5 8.0 7.5 7.0 6.5 6.0 5.5 5.0 4.5 4.0 3.5 3.0 2.5 2.0 1.5 1.0 0.5 0.0 -0.5
f1 (ppm)

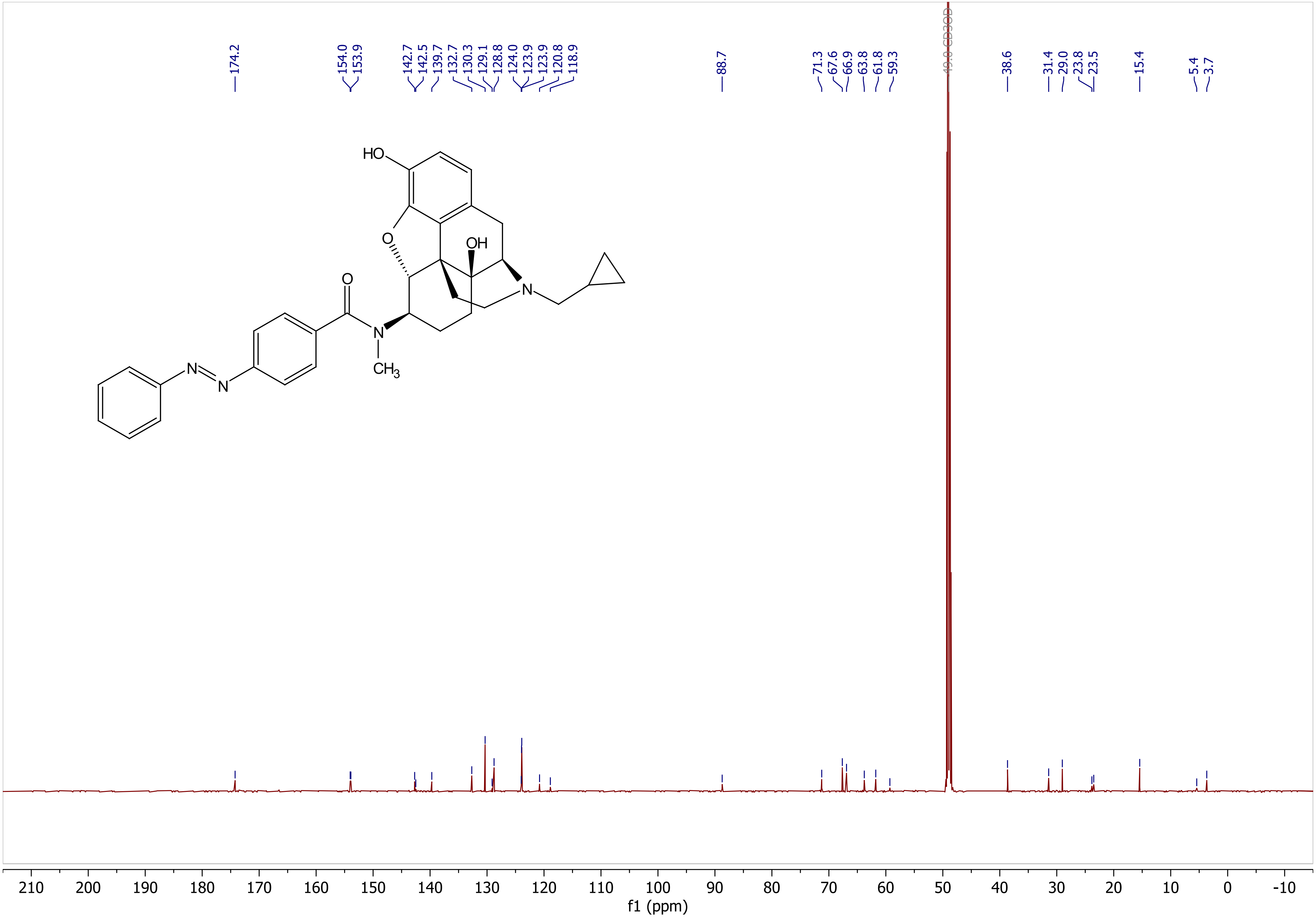
174.2
154.0
153.9
142.7
142.5
139.7
132.7
130.3
129.1
128.8
124.0
123.9
123.9
120.8
118.9
88.7
71.3
67.6
66.9
63.8
61.8
59.3
49.0 CD3OD
38.6
31.4
29.0
23.8
23.5
15.4
5.4
3.7
HO
O
OH
N
O
N
CH3
N
N
210 200 190 180 170 160 150 140 130 120 110 100 90 80 70 60 50 40 30 20 10 0 -10
f1 (ppm)

**<u>*N*-((4*R*,4a*S*,7*R*,7a*R*,12b*S*)-3-(cyclopropylmethyl)-4a,9-dihydroxy-2,3,4,4a,5,6,7,7a-octahydro-1H-4,12-methanobenzofuro[3,2-*e*]isoquinolin-7-yl)-*N*-methyl-3-((*E*)-phenyldiazenyl)benzamide (AM-2)</u>**

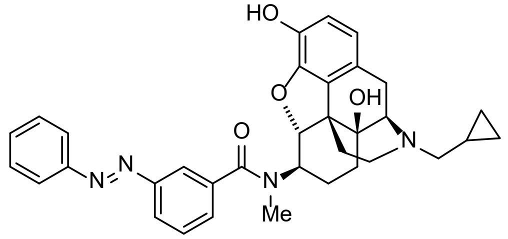


**AM-2**

**<u>Procedure:</u>** In a flame-dried 5 mL round bottom flask, naltrexone-derived amine (**1**) (14 mg, 0.040 mmol, 1.0 eq.), 3-(phenylazo)benzoic acid (**S-2**) (11 mg, 0.048 mmol, 1.2 eq.) and COMU coupling reagent (19 mg, 0.044 mmol, 1.1 eq.) were dissolved in anhydrous DMF (0.40 mL, 0.1 M). The reaction was stirred under argon at 0 °C, and anhydrous DIPEA (15 µL, 0.088 mmol, 2.2 eq.) was added. The reaction mixture was stirred at 0 °C for 1 h before the cooling bath was removed, and the solution was allowed to warm up and stirred at room temperature for another 16 h. Upon the completion of the reaction, the mixture was diluted with EtOAc:$Et_2O$ 1:1 (20 mL) and transferred to a separatory funnel. 0.1 M $NaHCO_{3(aq)}$ (10 mL) was added to the separatory funnel, and layers were separated. The aqueous layer was extracted with EtOAc (2x20 mL). The combined organic layer was extracted with 10% $LiCl_{(aq)}$(2x20 mL) and brine (1x20 mL). The organic layer was dried over $Na_2SO_4$, filtered, and concentrated *in vacuo*. The crude product was purified on silica gel (gradient from 0% to 10% MeOH in $CH_2Cl_2$ with 1% ammonia, 7 M in MeOH) to afford the desired **AM-2** (17.4 mg, 0.031 mmol, 77% yield) as an orange oil. For long-term storage, the compound was kept as a trifluoroacetic acid or a formic acid salt. Trifluoroacetic acid or formic acid (1.3 eq) in acetonitrile (99:1 MeCN:TFA/formic acid) was added to the compound and concentrated *in vacuo* to afford the salt in order to avoid the formation of *N*-oxides. The dilution of the acid was necessary to avoid the decomposition of the product.

**<u>Characterization</u>**:

**$^1$H NMR** (500 MHz, $CD_3OD$, exists as a mixture of rotamers, only peaks of the major rotamer shown) δ 8.51 (br s, 0H, formate salt peak), 7.90 (dt, *J* = 6.4, 1.7 Hz, 2H), 7.83 – 7.79 (m, 2H), 7.53 (tt, *J* = 21.6, 7.9 Hz, 5H), 6.60 (d, *J* = 8.2 Hz, 1H), 6.56 (d, *J* = 8.2 Hz, 1H), 4.91 (d, *J* = 8.1 Hz, 1H), 3.78 (d, *J* = 5.7 Hz, 1H), 3.55 (ddd, *J* = 12.7, 8.0, 4.4 Hz, 1H), 3.24 – 3.17 (m, 4H), 3.12 – 2.91 (m, 2H), 2.82 (dd, *J* = 13.4, 7.5 Hz, 1H), 2.58 (d, *J* = 8.1 Hz, 2H), 2.35 – 2.23 (m, 1H), 1.68 (t, *J* = 14.8 Hz, 2H), 1.55 (dd, *J* = 23.0, 11.2 Hz, 1H), 1.35 – 1.14 (m, 2H), 1.07 (d, *J* = 46.8 Hz, 1H), 0.72 (d, *J* = 34.8 Hz, 2H), 0.57 – 0.32 (m, 2H).
**$^{13}$C NMR** (126 MHz, $CD_3OD$, exists as a mixture of rotamers, only peaks of the major rotamer shown) δ 174.1, 153.8, 153.7, 142.8, 138.4, 132.7, 130.7, 130.5, 130.3, 129.9, 129.2, 126.3, 124.4, 124.0, 121.5, 121.0, 119.4, 88.4, 71.2, 69.5, 64.0, 61.6, 58.9, 56.2, 47.8, 31.3, 29.1, 24.1, 23.2, 7.0, 6.0, 3.4.
* note: the NMR of this compound at room temperature is a rotameric mixture.

**HRMS (ESI)**: m/z $[M+H]^+$ calcd for $C_{34}H_{36}N_4O_4$: 565.280932, found: 565.283586 (Δ = 4.7 ppm)
**$R_f$** (4% MeOH in $CH_2Cl_2$ with 1% ammonia, 7 M in methanol): 0.41; stains yellow with $KMnO_4$; detected by UV

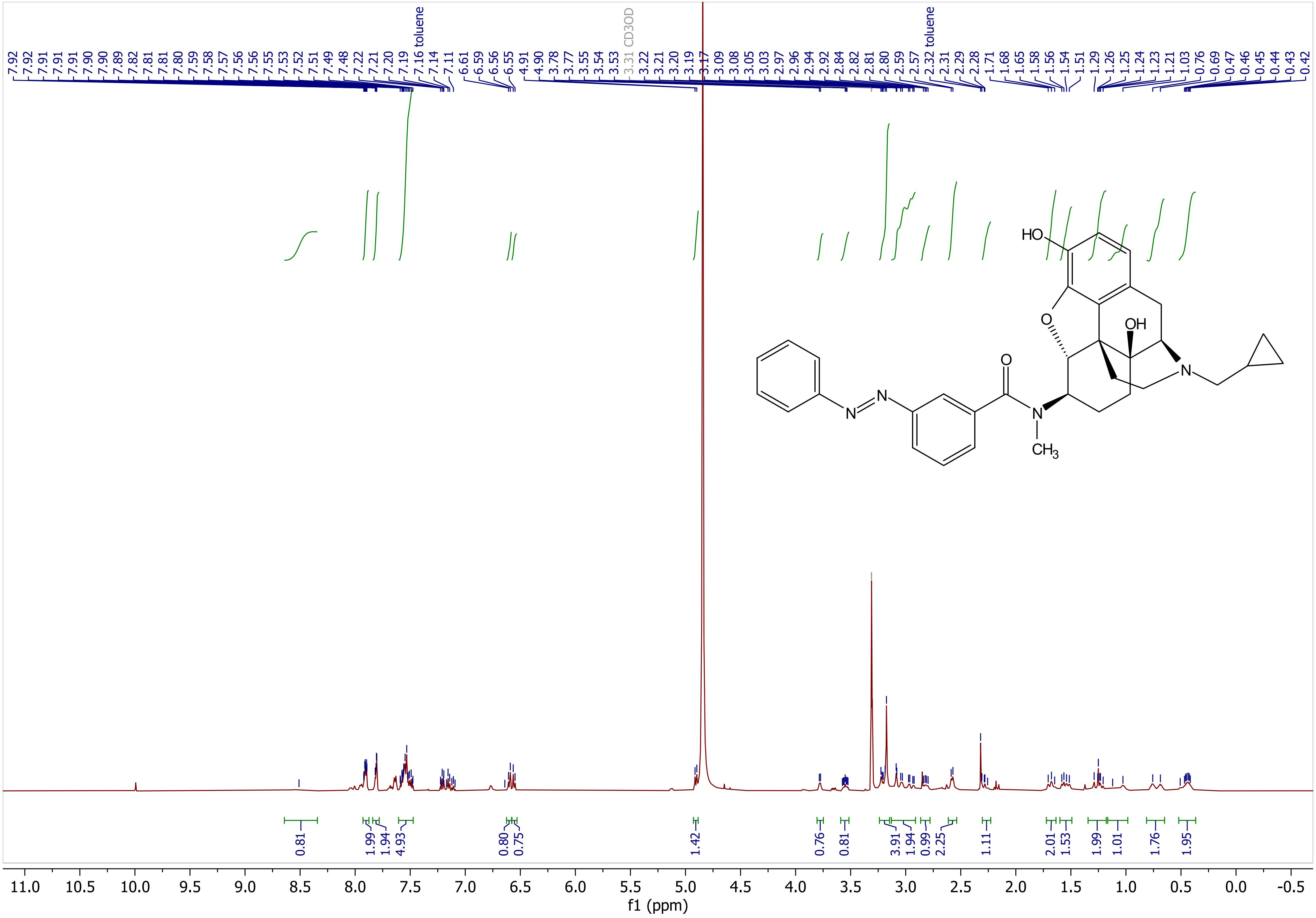

7.92 7.92 7.91 7.91 7.91 7.90 7.90 7.89 7.82 7.81 7.81 7.80 7.59 7.58 7.57 7.56 7.56 7.55 7.53 7.52 7.51 7.49 7.48 7.22 7.21 7.20 7.19 7.16 toluene 7.14 7.11 6.61 6.59 6.56 6.55 4.91 4.90 3.78 3.77 3.55 3.54 3.53 3.31 CD3OD 3.22 3.21 3.20 3.19 3.17 3.09 3.08 3.05 3.03 2.97 2.96 2.94 2.92 2.84 2.82 2.81 2.80 2.59 2.57 2.32 toluene 2.31 2.29 2.28 1.71 1.68 1.65 1.58 1.56 1.54 1.51 1.29 1.26 1.25 1.24 1.23 1.21 1.03 0.76 0.69 0.47 0.46 0.45 0.44 0.43 0.42
HO
O
OH
O
N
N
N
N
$CH_3$
0.81 1.99 1.94 4.93 0.80 0.75 1.42 0.76 0.81 3.91 1.94 0.99 2.25 1.11 2.01 1.53 1.99 1.01 1.76 1.95
11.0 10.5 10.0 9.5 9.0 8.5 8.0 7.5 7.0 6.5 6.0 5.5 5.0 4.5 4.0 3.5 3.0 2.5 2.0 1.5 1.0 0.5 0.0 -0.5
f1 (ppm)

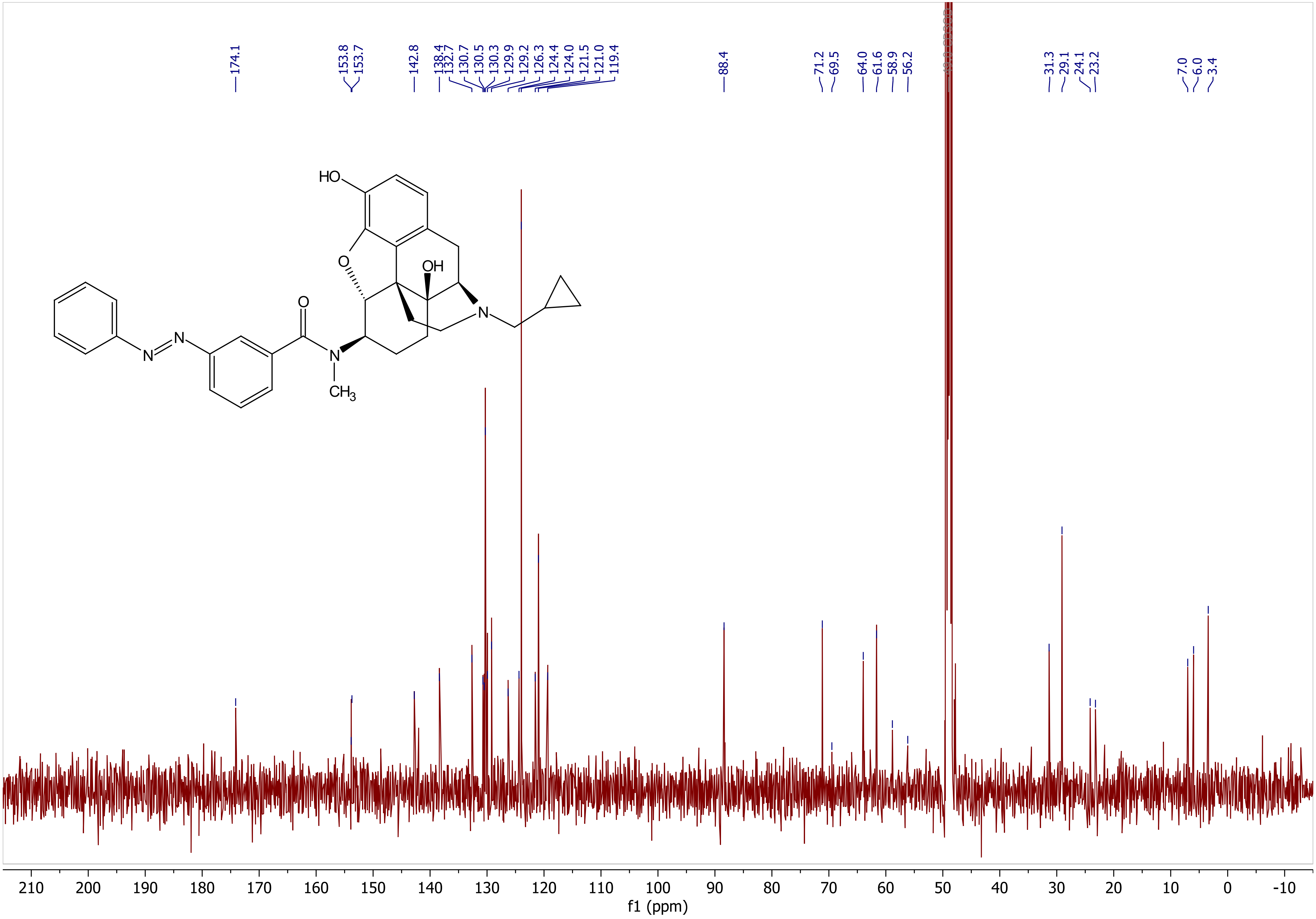

174.1
153.8
153.7
142.8
138.4
132.7
130.7
130.5
130.3
129.9
129.2
126.3
124.4
124.0
121.5
121.0
119.4
88.4
71.2
69.5
64.0
61.6
58.9
56.2
31.3
29.1
24.1
23.2
7.0
6.0
3.4
HO
O
OH
O
N
N
N
N
CH3
f1 (ppm)
210 200 190 180 170 160 150 140 130 120 110 100 90 80 70 60 50 40 30 20 10 0 -10

**_N_-((4_R_,4a_S_,7_R_,7a_R_,12b_S_)-3-(cyclopropylmethyl)-4a,9-dihydroxy-2,3,4,4a,5,6,7,7a-octahydro-1H-4,12-methanobenzofuro[3,2-_e_]isoquinolin-7-yl)-_N_-methyl-2-(4-((_E_)-phenyldiazenyl)phenyl)acetamide (AM-3)**

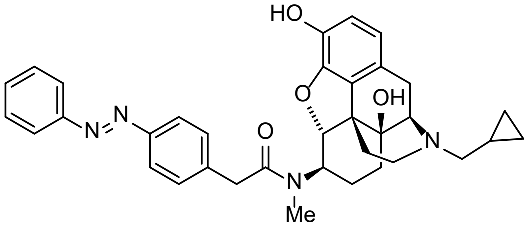


**AM-3**

**Procedure:** In a flame-dried 5 mL round bottom flask, naltrexone-derived amine (**1**) (14 mg, 0.040 mmol, 1.0 eq.), azobenzene (**2**) (11 mg, 0.048 mmol, 1.2 eq.) and COMU coupling reagent (19 mg, 0.044 mmol, 1.1 eq.) were dissolved in anhydrous DMF (0.40 mL, 0.1 M). The reaction was stirred under argon at 0 °C, and anhydrous DIPEA (15 µL, 0.088 mmol, 2.2 eq.) was added. The reaction mixture was stirred at 0 °C for 1 h before the cooling bath was removed, and the solution was allowed to warm up and stirred at room temperature for another 16 h. Upon the completion of the reaction, the mixture was diluted with EtOAc:$Et_2O$ 1:1 (20 mL) and transferred to a separatory funnel. 0.1 M $NaHCO_3$ (10 mL) was added to the separatory funnel, and layers were separated. The aqueous layer was extracted with EtOAc (2x20 mL). The combined organic layer was extracted with 10% $LiCl_{(aq)}$(2x20 mL) and brine (1x20 mL). The organic layer was dried over $Na_2SO_4$, filtered, and concentrated *in vacuo*. The crude product was purified on silica gel (gradient from 0% to 10% MeOH in $CH_2Cl_2$ with 1% ammonia, 7 M in MeOH) to afford the desired opto-opioid (18.0 mg, 0.031 mmol, 78% yield) as an orange oil. For long-term storage, the compound was kept as a trifluoroacetic acid or a formic acid salt. Trifluoroacetic acid or formic acid (1.3 eq) in acetonitrile (99:1 MeCN:TFA/formic acid) was added to the compound and concentrated *in vacuo* to afford the salt in order to avoid the formation of *N*-oxides. The dilution of the acid was necessary to avoid the decomposition of the product. It has been confirmed that neither TFA nor formate salt has influence on biological activity.

**Characterization**:

**$^1$H NMR** (500 MHz, DMSO-d6, exists as 8:2 rotamers, * denotes minor rotamer) δ 8.22 (br s, 0H, formate salt peak), 7.91 – 7.82 (m, 2H+3H*), 7.77 – 7.70 (m, 2H), 7.63 – 7.51 (m, 3H+3H*), 7.45 (d, *J* = 8.1 Hz, 2H*), 7.10 (d, *J* = 8.3 Hz, 2H), 6.72 (d, *J* = 8.1 Hz, 1H), 6.64 (d, *J* = 8.1 Hz, 1H), 6.58 (d, *J* = 8.1 Hz, 1H*), 6.52 (d, *J* = 8.1 Hz, 1H*), 4.71 (d, *J* = 8.2 Hz, 1H*), 4.62 (d, *J* = 8.0 Hz, 1H), 3.90 – 3.78 (m, 2H*), 3.70 (q, *J* = 15.7 Hz, 2H), 3.57 (ddd, *J* = 12.2, 7.7, 3.3 Hz, 1H+1H*), 3.16 – 3.05 (m, 2H*), 3.02 (h, *J* = 5.7 Hz, 1H), 2.98 (d, *J* = 3.3 Hz, 1H), 2.81 (s, 3H), 2.74 (s, 3H*), 2.69 – 2.54 (m, 1H+2H*), 2.54 – 2.46 (m, 1H), 2.36 (qt, *J* = 12.7, 7.0 Hz, 2H+2H*), 2.20 (qd, *J* = 12.6, 5.1 Hz, 1H+1H*), 2.00 (qd, *J* = 11.9, 3.4 Hz, 2H), 1.50 (d, *J* = 13.2 Hz, 1H*), 1.40 (dt, *J* = 13.2, 3.2 Hz, 1H), 1.35 – 1.28 (m, 1H), 1.27 – 1.21 (m, 2H*), 1.18 – 1.09 (m, 1H+1H*), 1.03 (dt, *J* = 12.6, 3.6 Hz, 1H), 0.82 (dd, *J* = 9.5, 4.3 Hz, 1H+1H*), 0.50 – 0.42 (m, 2H+2H*), 0.17 – 0.09 (m, 2H+2H*).
**$^{13}$C NMR** (126 MHz, DMSO-d6, exists as 8:2 rotamers, * denotes minor rotamer) δ 170.0, 152.0*, 151.9, 150.5*, 150.4, 142.1*, 141.5, 140.9, 140.7*, 140.1, 139.8*, 131.8, 131.4, 131.3*, 130.2*, 130.0, 129.4, 123.7, 122.5, 122.3, 119.2, 117.1, 88.5, 87.7*, 69.5*, 69.4, 65.9, 61.7*, 61.5, 58.3, 57.6, 47.2, 47.0*, 43.7, 37.9, 30.6, 29.7, 29.6*, 27.8, 22.4*, 22.2, 21.4*, 9.0, 3.7, 3.5.
* note: the NMR of this compound at room temperature is a rotameric mixture; a variable temperature NMR was conducted to confirm the existence of rotamers.

**HRMS (ESI)**: m/z $[M+H]^+$ calcd for $C_{35}H_{38}N_4O_4$: 579.2971, found: 579.2982 (Δ = 1.9 ppm)
**$R_f$** (4% MeOH in $CH_2Cl_2$ with 1% ammonia, 7 M in methanol): 0.29; stains yellow with $KMnO_4$; detected by UV

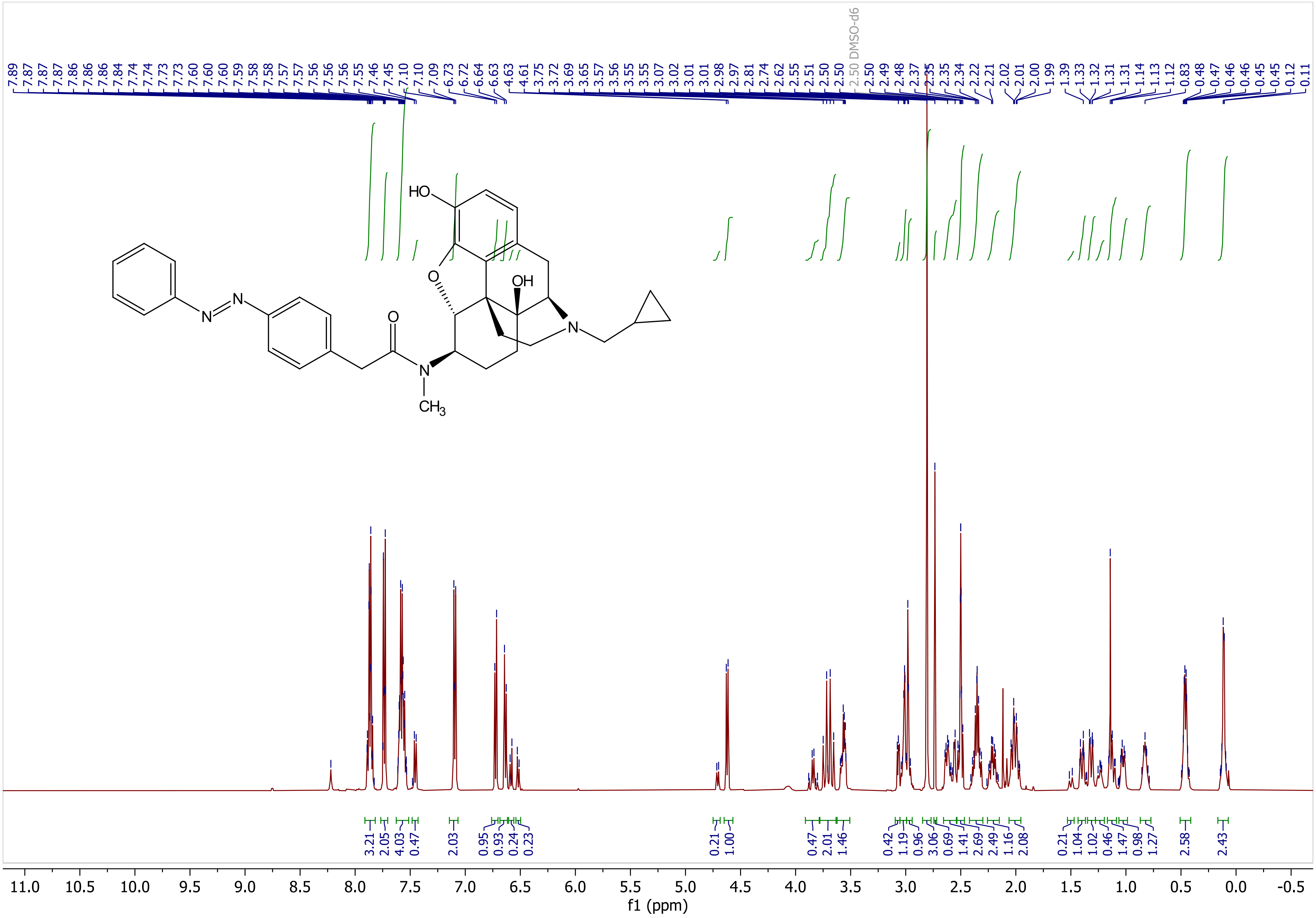

7.89 7.87 7.87 7.87 7.86 7.86 7.86 7.84 7.74 7.74 7.73 7.73 7.60 7.60 7.60 7.59 7.58 7.58 7.57 7.57 7.56 7.56 7.56 7.55 7.46 7.45 7.10 7.10 7.09 6.73 6.72 6.64 6.63 4.63 4.61 3.75 3.72 3.69 3.65 3.57 3.56 3.55 3.55 3.07 3.02 3.01 2.98 2.97 2.81 2.74 2.62 2.55 2.51 2.50 2.50 2.50 DMSO-d6 2.50 2.49 2.48 2.37 2.35 2.35 2.34 2.22 2.21 2.02 2.01 2.00 1.99 1.39 1.33 1.32 1.31 1.31 1.14 1.13 1.12 0.83 0.48 0.47 0.46 0.46 0.45 0.45 0.12 0.11
HO
O
OH
N
N
N
O
N
CH3
3.21 2.05 4.03 0.47 2.03 0.95 0.93 0.24 0.23 0.21 1.00 0.47 2.01 1.46 0.42 1.19 0.96 3.06 0.69 1.41 2.69 2.49 1.16 2.08 0.21 1.04 1.02 0.46 1.47 0.98 1.27 2.58 2.43
11.0 10.5 10.0 9.5 9.0 8.5 8.0 7.5 7.0 6.5 6.0 5.5 5.0 4.5 4.0 3.5 3.0 2.5 2.0 1.5 1.0 0.5 0.0 -0.5
f1 (ppm)

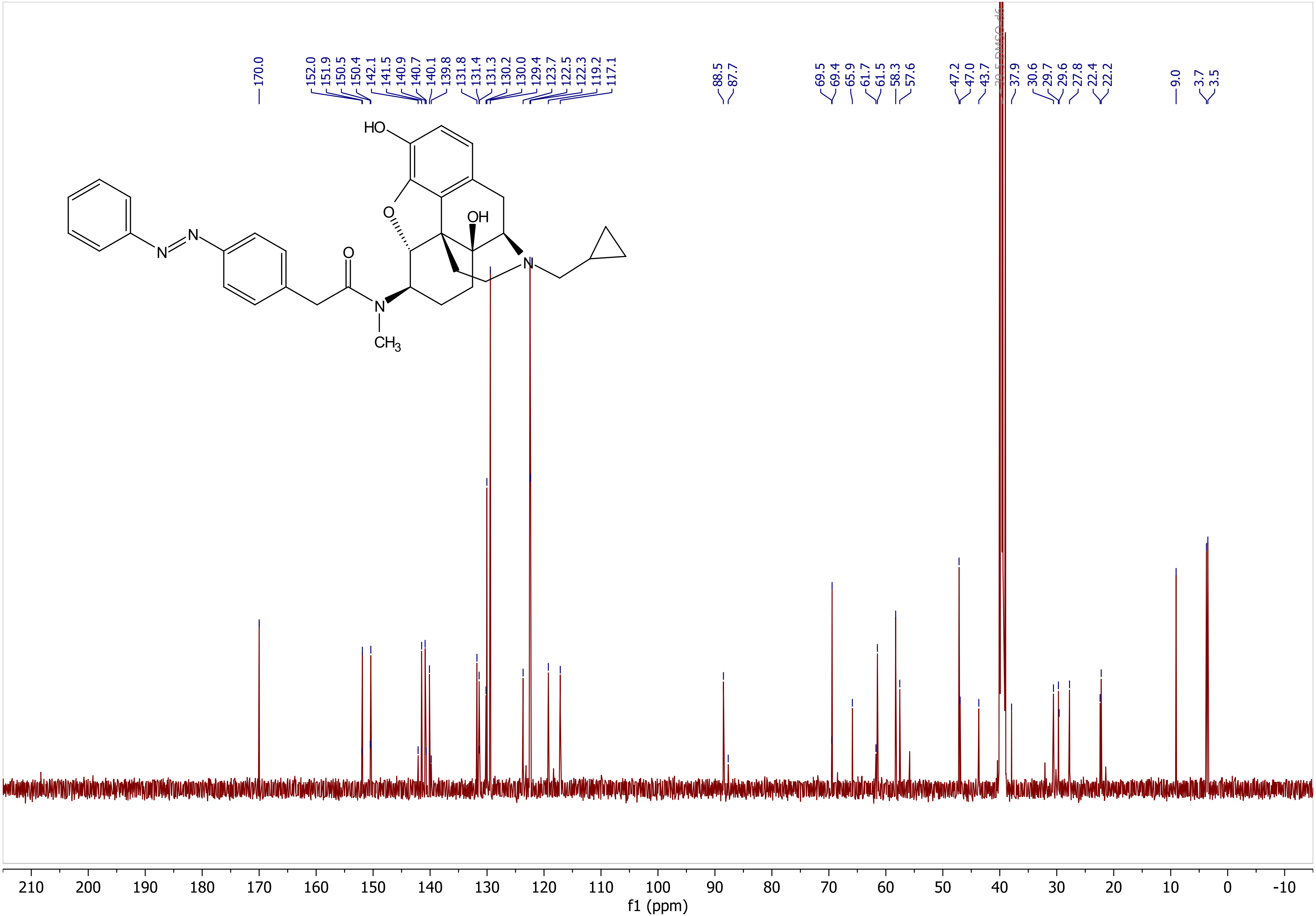
170.0
152.0
151.9
150.5
150.4
142.1
141.5
140.9
140.7
140.1
139.8
131.8
131.4
131.3
130.2
130.0
129.4
123.7
122.5
122.3
119.2
117.1
88.5
87.7
69.5
69.4
65.9
61.7
61.5
58.3
57.6
47.2
47.0
43.7
39.5 DMSO-d6
37.9
30.6
29.7
29.6
27.8
22.4
22.2
9.0
3.7
3.5
HO
O
OH
N
N
N
O
N
CH3
210
200
190
180
170
160
150
140
130
120
110
100
90
80
70
60
50
40
30
20
10
0
-10
f1 (ppm)

AM-3: Variable Temperature (VT) NMR Experiment

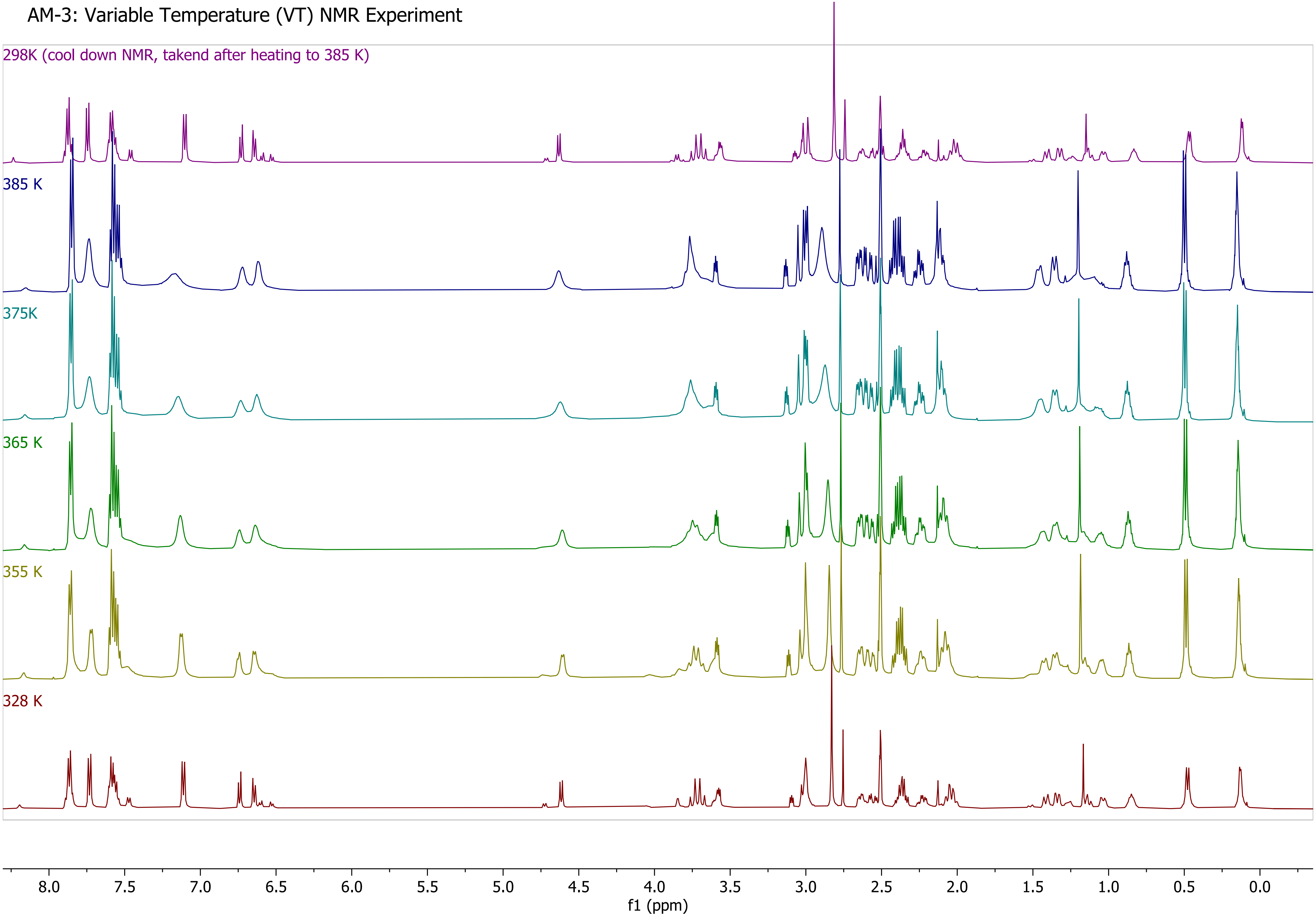

***N*-((4*R*,4a*S*,7*R*,7a*R*,12b*S*)-3-(cyclopropylmethyl)-4a,9-dihydroxy-2,3,4,4a,5,6,7,7a-octahydro-1H-4,12-methanobenzofuro[3,2-*e*]isoquinolin-7-yl)-*N*-methyl-2-(3-((*E*)-phenyldiazenyl)phenyl)acetamide (AM-4)**

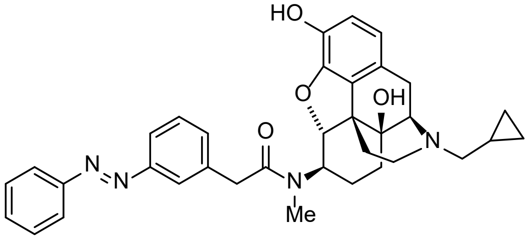


**AM-4**

**Procedure:** In a flame-dried 5 mL round bottom flask, naltrexone-derived amine (**1**) (9.0 mg, 0.025 mmol, 1.0 eq.), azobenzene (**S-4**) (7.9 mg, 0.033 mmol, 1.3 eq.) and COMU coupling reagent (13 mg, 0.030 mmol, 1.2 eq.) were dissolved in anhydrous DMF (0.25 mL, 0.1 M). The reaction was stirred under argon at 0 °C, and anhydrous DIPEA (10 µL, 0.056 mmol, 2.2 eq.) was added. The reaction mixture was stirred at 0 °C for 1 h before the cooling bath was removed, and the solution was allowed to warm up and stirred at room temperature for another 16 h. Upon the completion of the reaction, the mixture was diluted with EtOAc:$Et_2O$ 1:1 (20 mL) and transferred to a separatory funnel. 0.1 M $NaHCO_{3(aq)}$ (10 mL) was added to the separatory funnel, and layers were separated. The aqueous layer was extracted with EtOAc (2x20 mL). The combined organic layer was extracted with 10% $LiCl_{(aq)}$ (2x20 mL) and brine (1x20 mL). The organic layer was dried over $Na_2SO_4$, filtered, and concentrated *in vacuo*. The crude product was purified on silica gel (gradient from 0% to 10% MeOH in $CH_2Cl_2$ with 1% ammonia, 7 M in MeOH) to afford the desired **AM-4** (10.3 mg, 0.018 mmol, 59% yield) as an orange oil. For long-term storage, the compound was kept as a trifluoroacetic acid or a formic acid salt. Trifluoroacetic acid or formic acid (1.3 eq) in acetonitrile (99:1 MeCN:TFA/formic acid) was added to the compound and concentrated *in vacuo* to afford the salt in order to avoid the formation of *N*-oxides. The dilution of the acid was necessary to avoid the decomposition of the product.

**Characterization**:

**$^1$H NMR** (600 MHz, $CD_3OD$, exists as 8:2 rotamers, * denotes minor rotamer) δ 7.89 (dt, *J* = 7.8, 1.9 Hz, 2H), 7.84 – 7.79 (m, 2H*), 7.71 (dq, *J* = 7.7, 2.8 Hz, 2H*), 7.71 – 7.66 (m, 1H), 7.62 – 7.49 (m, 3H+3H*), 7.44 (d, *J* = 7.7 Hz, 1H*), 7.35 (td, *J* = 7.8, 1.8 Hz, 1H), 7.24 (d, *J* = 2.0 Hz, 1H), 7.02 (d, *J* = 7.6 Hz, 1H), 6.83 (dd, *J* = 8.1, 1.9 Hz, 1H), 6.79 – 6.74 (m, 1H*), 6.64 (d, *J* = 7.8 Hz, 1H), 6.59 (d, *J* = 8.4 Hz, 1H*), 4.64 (dd, *J* = 8.2, 1.8 Hz, 1H), 4.33 (qd, *J* = 7.2, 1.9 Hz, 2H*), 3.86 (dd, *J* = 15.4, 1.9 Hz, 1H), 3.76 (d, *J* = 15.4 Hz, 1H), 3.67 (ddt, *J* = 11.4, 6.4, 2.9 Hz, 1H+1H*), 3.23 – 3.18 (m, 3H*), 3.15 – 3.07 (m, 1H+1H*), 3.02 (d, *J* = 18.5 Hz, 1H), 2.92 (d, *J* = 1.8 Hz, 3H), 2.84 (d, *J* = 1.9 Hz, 3H*), 2.72 (td, *J* = 15.1, 5.9 Hz, 1H+1H*), 2.49 (dt, *J* = 14.9, 7.2 Hz, 2H), 2.39 (dd, *J* = 12.9, 6.6 Hz, 1H+1H*), 2.28 (td, *J* = 12.5, 4.8 Hz, 1H+1H*), 2.18 (dt, *J* = 13.1, 6.5 Hz, 1H+1H*), 1.98 (qd, *J* = 12.9, 2.7 Hz, 1H+1H*), 1.46 (dd, *J* = 12.1, 3.4 Hz, 1H+1H*), 1.38 – 1.25 (m, 1H+1H*), 0.99 (td, *J* = 13.5, 3.2 Hz, 1H+1H*), 0.92 – 0.82 (m, 1H+1H*), 0.74 (dd, *J* = 13.0, 3.7 Hz, 1H), 0.61 – 0.47 (m, 2H), 0.14 (t, *J* = 4.0 Hz, 2H).
**$^{13}$C NMR** (126 MHz, $CD_3OD$, exists as 8:2 rotamers, only resonance of major rotamer shown) δ 173.9, 154.3, 154.0, 143.2, 142.4, 138.2, 132.9, 132.4, 132.1, 130.5, 130.3, 125.3, 123.8, 123.8, 122.2, 121.0, 119.0, 101.4, 90.2, 71.5, 63.7, 60.1, 60.0, 45.5, 42.7, 31.7, 31.0, 28.8, 23.7, 23.4, 9.8, 4.5, 4.1.
* note: the NMR of this compound at room temperature is a rotameric mixture.

**HRMS (ESI)**: m/z $[M+H]^+$ calcd for $C_{35}H_{38}N_4O_4$: 579.296582, found: 579.297980 (Δ = 2.4 ppm)
**$R_f$** (4% MeOH in $CH_2Cl_2$ with 1% ammonia, 7 M in methanol): 0.27; stains yellow with $KMnO_4$; detected by UV

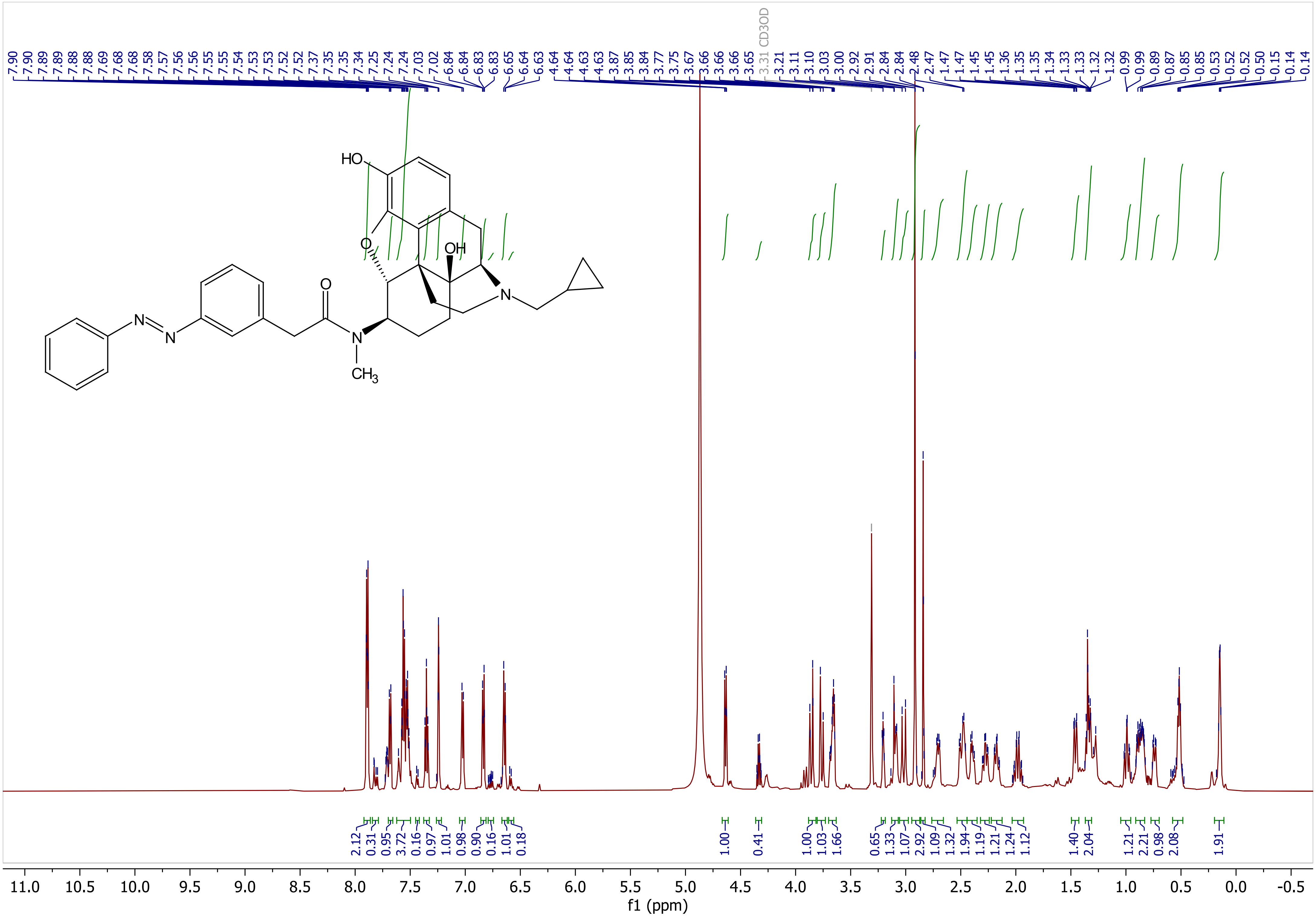

7.90
7.90
7.89
7.89
7.88
7.88
7.69
7.68
7.68
7.58
7.57
7.56
7.56
7.55
7.55
7.54
7.53
7.53
7.52
7.52
7.37
7.35
7.35
7.34
7.25
7.24
7.24
7.03
7.02
6.84
6.84
6.83
6.83
6.65
6.64
6.63
4.64
4.64
4.63
4.63
3.87
3.85
3.84
3.77
3.75
3.67
3.66
3.66
3.65
3.31 CD3OD
3.21
3.11
3.10
3.03
3.00
2.92
2.91
2.84
2.84
2.48
2.47
1.47
1.47
1.45
1.45
1.36
1.35
1.34
1.33
1.33
1.32
1.32
0.99
0.99
0.89
0.87
0.85
0.85
0.53
0.52
0.52
0.50
0.15
0.14
0.14
HO
O
OH
N
O
N
N
N
CH3
2.12
0.31
0.95
3.72
0.16
0.97
1.01
0.98
0.90
0.16
1.01
0.18
1.00
0.41
1.00
1.03
1.66
0.65
1.33
1.07
2.92
1.09
1.32
1.94
1.19
1.21
1.24
1.12
1.40
2.04
1.21
2.21
0.98
2.08
1.91
11.0 10.5 10.0 9.5 9.0 8.5 8.0 7.5 7.0 6.5 6.0 5.5 5.0 4.5 4.0 3.5 3.0 2.5 2.0 1.5 1.0 0.5 0.0 -0.5
f1 (ppm)

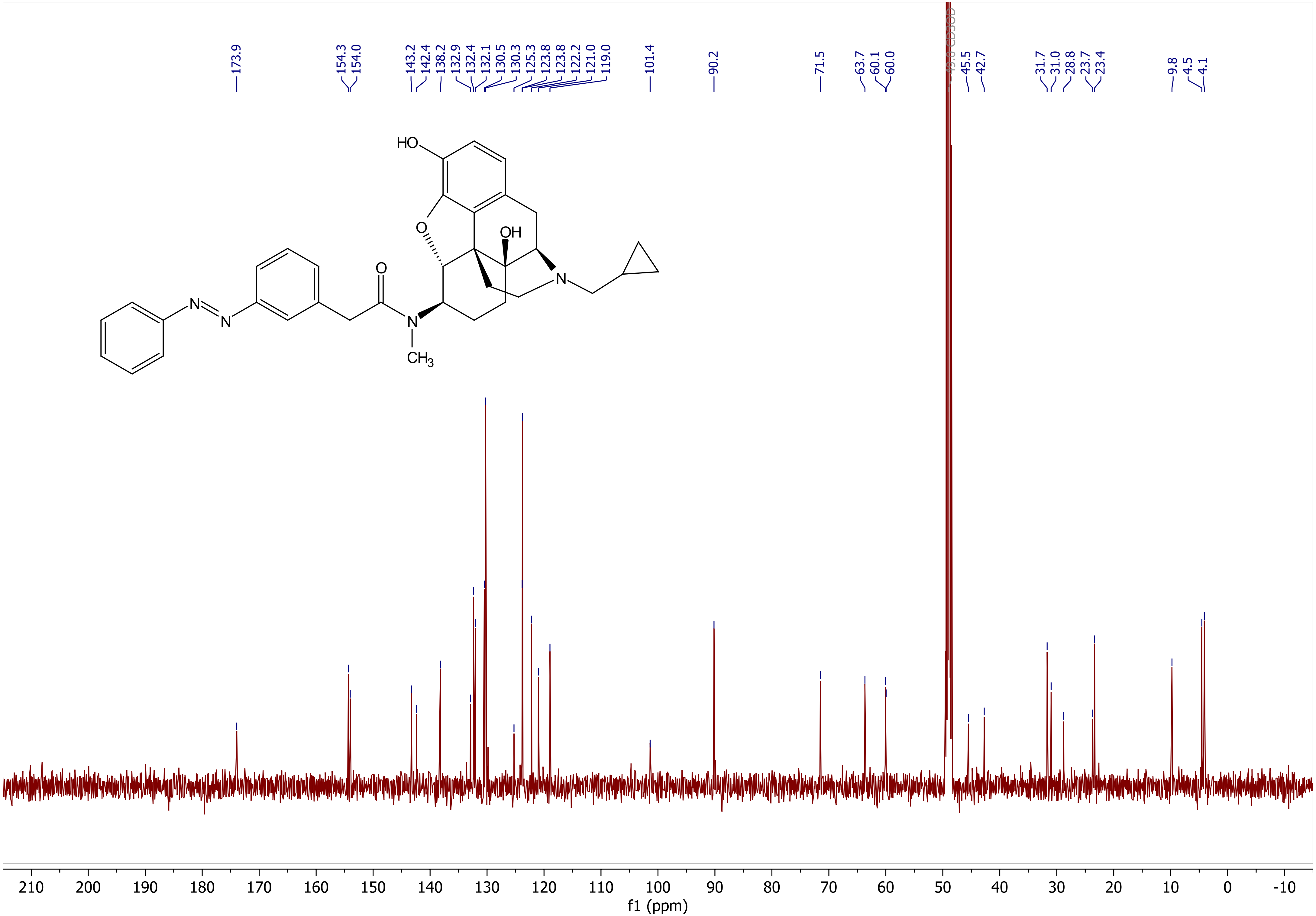
173.9
154.3
154.0
143.2
142.4
138.2
132.9
132.4
132.1
130.5
130.3
125.3
123.8
123.8
122.2
121.0
119.0
101.4
90.2
71.5
63.7
60.1
60.0
45.5
42.7
31.7
31.0
28.8
23.7
23.4
9.8
4.5
4.1
HO
O
OH
O
N
N
N
N
CH3
210 200 190 180 170 160 150 140 130 120 110 100 90 80 70 60 50 40 30 20 10 0 -10
f1 (ppm)
